# HP1 dimerization drives the condensation and segregation of H3K9me-marked chromatin

**DOI:** 10.1101/2025.11.13.687974

**Authors:** Karthik Eswara, Jennifer Semple, Francesca Rivas-Cuestas, Fernando Muzzopappa, Shakila Ali, Sonia El Mouridi, Yogesh Ostwal, Christian Frøkjær-Jensen, Fabian Erdel, Peter Meister, Wolfgang Fischle

## Abstract

Heterochromatin protein 1 (HP1) is a conserved chromatin-associated factor implicated in the establishment and maintenance of H3K9me-marked heterochromatin, potentially through phase separation–mediated condensation. Whether HP1 promotes heterochromatin condensation primarily through its dimerization or liquid–liquid phase separation (LLPS) remains unresolved. Using the *C. elegans* HP1 orthologue HPL-2 and a combined *in vitro–in vivo* approach, we systematically dissected the molecular determinants of HPL-2 function in heterochromatin condensation. Through specific mutants, we demonstrate that HPL-2 dimerization is essential for condensing H3K9me chromatin arrays *in vitro* and for maintaining HPL-2 and H3K9me heterochromatin foci in *C. elegans* embryos, while LLPS enhances or stabilizes these foci. We further show that HPL-2 dimerization is sufficient to mediate segregation of H3K9me from unmodified chromatin arrays *in vitro*, forming spatially distinct H3K9me-enriched condensates. Surprisingly, HPL-2 mutants defective in heterochromatin foci formation cause only minor transcriptional changes among genes associated with H3K9me-marked heterochromatin, implying that HP1-dependent heterochromatin foci and gene silencing are not tightly coupled *in vivo*. Nonetheless, these mutant *C. elegans* exhibit profound physiological and developmental defects. Our findings establish dimerization as the principal molecular mechanism of HP1-driven H3K9me-chromatin condensation, elucidate an auxiliary role of LLPS, and reveal the uncoupling between HP1-dependent heterochromatin and transcriptional regulation.

**GRAPHICAL ABSTRACT:** 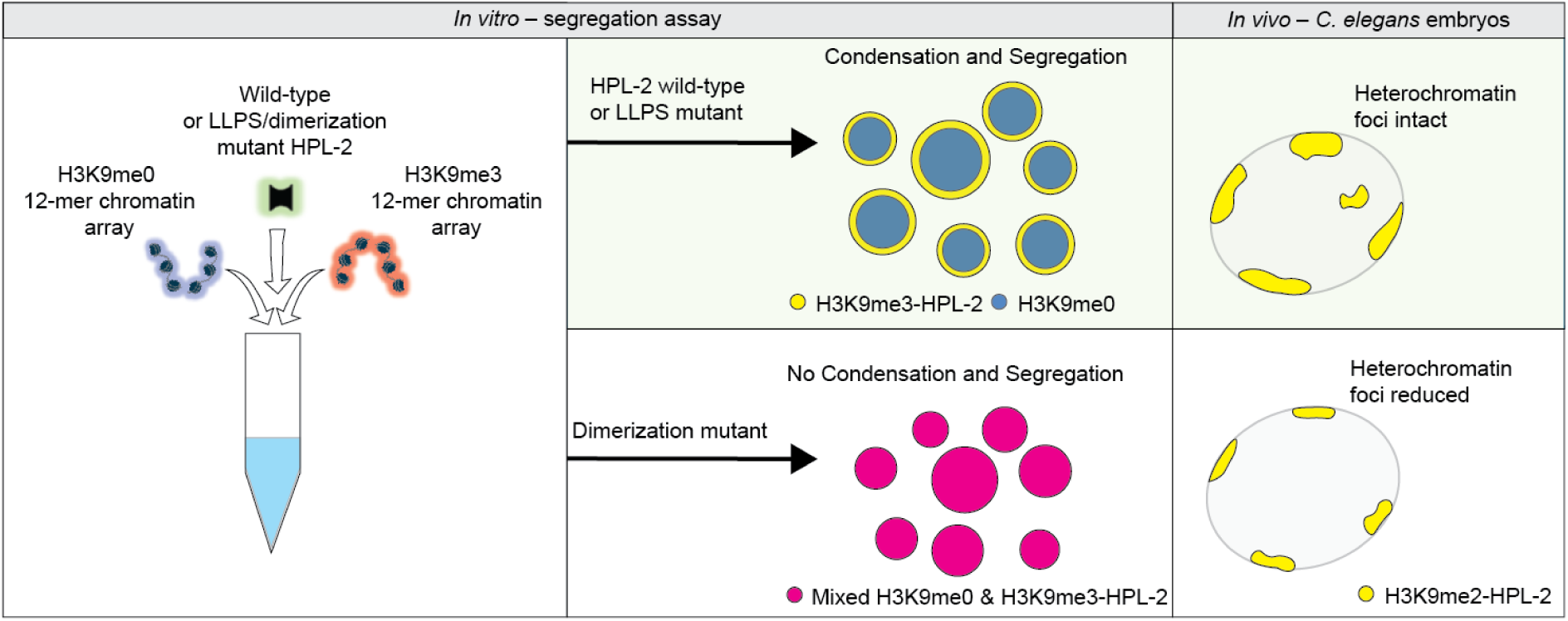

## INTRODUCTION

Constitutive heterochromatin is a defining feature of eukaryotic chromosome organisation. It is marked by histone H3 lysine 9 di- and trimethylation (H3K9me) and forming densely packed domains that maintain genome stability and silence repetitive elements [1, 2]. Despite its visibly compact organisation, the molecular basis of heterochromatin condensation remains poorly understood. Two principal models of condensation have been proposed: one based on liquid–liquid phase separation (LLPS) and the other on chromatin polymer bridging [3, 4]. Distinguishing between these mechanisms is central to understanding how chromatin architecture is established and maintained.

In the LLPS model, heterochromatin is thought to form through weak, multivalent interactions among chromatin-associated proteins and nucleic acids, producing dynamic, liquid-like condensates into which chromatin partitions [3, 5–7]. In contrast, the polymer-bridging model posits that condensation is driven by proteins that physically bridge neighbouring chromatin fibres, generating a condensed chromatin polymer network [3, 8–10]. Although both mechanisms have been proposed to generate dense chromatin domains, whether one or both operate in cells remains unresolved.

Among the factors implicated in heterochromatin condensation, Heterochromatin Protein 1 (HP1) has emerged as a key player. HP1 family proteins are conserved chromatin-associated factors that bind H3K9me2/3 through their N-terminal chromodomain (CD) and form homodimers through their C-terminal chromoshadow domain (CSD), connected by an intrinsically disordered hinge region enriched in charged residues [11–13]. This architecture equips HP1 with two potential modes of chromatin condensation once recruited to heterochromatin: (i) the capacity to self-associate through weak multivalent interactions mediated by charged residues, potentially driving LLPS of HP1 and the underlying chromatin, and (ii) the ability to bridge H3K9me nucleosomes via CSD-mediated dimerization.

Recombinant mammalian HP1α [14], *Drosophila* HP1a [15], and yeast Swi6 [16] can form liquid-like droplets whose formation is enhanced by phosphorylation or nucleic-acid binding. H3K9-methylated nucleosomes preferentially partition into these condensates. These findings inspired the idea that LLPS of HP1 proteins underlies heterochromatin condensation. In contrast, cryo-EM structures revealed that each HP1 dimer can simultaneously engage two H3K9me3-modified nucleosomes [17], providing a direct, putative physical mechanism for polymer bridging. Bridging has also been proposed for Swi6, which tetramerises *in vitro* on nucleosomes containing H3K9me and connects neighbouring nucleosomes via unoccupied chromodomain interfaces [18]. Similarly, human HP1β dynamically associates with H3K9me3-enriched chromatin arrays through dimeric bridging interactions [13]. Since HP1 can display both liquid-like behaviour and bridging activity *in vitro*, the relative contributions of these properties to heterochromatin condensation within the crowded environment of the nucleus have remained challenging to disentangle. Moreover, while heterochromatin is visually compact and transcriptionally silent [19], whether condensation itself enforces repression, or whether these are separable phenomena, remains uncertain. Resolving the molecular and physiological working mode of HP1 requires an experimental system in which its biochemical and biophysical properties can be precisely perturbed and where the impact of such perturbations on chromatin organisation and transcription can be evaluated.

Here, we investigated HP1 function in the nematode *Caenorhabditis elegans*, a tractable metazoan model for heterochromatin organisation. *C. elegans* chromosomes are partitioned into gene-rich central regions and repeat-rich arms enriched for H3K9me and the HP1 orthologue HPL-2, broadly marking constitutive heterochromatin [20–23]. While HPL-2 localizes to repressive chromatin domains and is essential for repeat silencing and normal development [20, 24–26], whether and how the factor drives heterochromatin condensation has not been defined. We combined *in vitro* biochemical reconstitution, cellular imaging, and functional genomics to dissect the mechanisms by which HPL-2 condenses chromatin *in vitro* and *in vivo*. Specifically, we tested whether heterochromatin condensation depends on dimerization mediated HPL-2 bridging or LLPS and examined how loss of condensation affects transcription within heterochromatin.

## MATERIAL AND METHODS

A complete list of plasmids, peptides, oligonucleotide primers, *C. elegans* strains, and antibodies used in this study is provided in the Supplementary Tables S1 to S6.

### *In silico* sequence and structure analysis

Protein sequences of HP1 homologs were retrieved from UniProt: *Caenorhabditis elegans* HPL-2 (G5EDE2) and HPL-1 (G5EET5), *Homo sapiens* HP1α (P45973), HP1β (P83916), HP1γ (Q13185), *Mus musculus* HP1α (Q61686), *Danio rerio* HP1 (A1L223), *Pristionchus pacificus* HPL-1 (A0A2A6CDA1), *Drosophila melanogaster* HP1a (P05205), and *Schizosaccharomyces pombe* Swi6 (P40381). Multiple sequence alignments were generated using ClustalW (v2.1) [27] within Jalview (v2.11.4.1) [28]. Predicted secondary structure elements for HPL-2 were obtained from the AlphaFold Protein Structure Database (AF-G5EDE2-F1-v4) [29] and visualized in PyMOL (v2.5) [30]. Pairwise sequence similarity scores for individual domains were calculated using the Sequence Manipulation Suite and visualized in Python (Google Colab). Protein–peptide complex structures were modelled using AlphaFold (v2.3) via ColabFold [29], with *C. elegans* HPL-2 (two copies) and an 18-residue H3 peptide as inputs. Predictions were generated under default settings using MMseqs2 for multiple sequence alignment. Models with the highest confidence (based on pLDDT and PAE scores) were retained and analysed in PyMOL. Structural features such as dimerization interfaces and aromatic cage residues were compared with published human HP1α structures (PDB: 3FDT and 3I3C) to assess conservation of interaction modes.

### Purification of recombinant HPL-2 proteins

cDNAs corresponding to the ORF of wild-type and mutant HPL-2 and encoding for a 10xHIS-tag followed by a TEV protease cleavage site at the N-terminus were cloned into pET16b, or pET-28a(+) vectors using Gibson assembly or synthetically constructed by Twist Bioscience (San Francisco, USA) (Table S1). Recombinant proteins were expressed in *E. coli* BL21-CodonPlus (DE3)-RIL cells transformed with the appropriate expression plasmid. Details of expression constructs are available upon request. Cultures were grown in 2xYT medium containing 100 µg/mL of appropriate antibiotic at 37°C to an OD₆₀₀ of 0.4–0.7. For all proteins, expression was induced with 0.5 mM isopropyl β-D-1-thiogalactopyranoside (IPTG) and continued for 16–18 h at 18°C. For histone proteins, induction was carried out with 1 mM IPTG for 4 h at 37°C. Cells were harvested by centrifugation and stored at –80°C until purification. His-tagged proteins were purified by Ni–NTA affinity chromatography. Bacterial pellets were resuspended in lysis buffer (50 mM Tris–HCl pH 8.0, 300 mM NaCl, 10 mM imidazole, 1 mM TCEP, protease inhibitor cocktail (ULTRA EDTA-free protease inhibitor, cOmplete™, Sigma) and lysed using a high-pressure homogenizer (Emulsiflex-C5, Avestin). Clarified lysates were incubated with Ni–NTA resin (HisPur™ Ni-NTA Resin, Thermo Fisher), washed with buffer containing 1 M NaCl and 10 mM imidazole, and eluted with 250 mM imidazole. His-tags were removed by overnight digestion with recombinant TEV protease (1:25 w/w) during dialysis against buffer containing 20 mM HEPES pH 8.0, 150 mM NaCl, 1 mM TCEP, and 10% (v/v) glycerol. The cleaved protein was separated from TEV protease and uncleaved material by a second Ni–NTA purification step. Protein purity and cleavage efficiency were verified by SDS– PAGE and Coomassie staining. Proteins were concentrated, flash-frozen in liquid nitrogen, and stored at –80°C.

### Fluorophore labelling of recombinant HPL-2 proteins

Purified wild-type and mutant HPL-2 proteins were dialyzed into 100 mM sodium bicarbonate buffer (pH 8.0) and labelled with Alexa Fluor™ 555 Succinimidyl Ester (Invitrogen) at a protein-to-dye molar ratio of 1:5. Reactions were incubated for 1 h at room temperature in the dark with gentle mixing. Unreacted dye was removed using Pierce™ Dye Removal Columns (Thermo Fisher) according to the manufacturer’s instructions, and labelled proteins were dialyzed into 20 mM HEPES pH 8.0, 150 mM NaCl, 1 mM TCEP, and 10% (v/v) glycerol. Protein concentration and labelling efficiency were determined spectrophotometrically by recording absorbance at 280 nm and 555 nm. Labelled proteins were aliquoted and stored at –80°C until use in phase separation assays.

### Purification of recombinant histones

Recombinant full-length and truncated histones were purified as described previously [13]. Briefly, *Xenopus laevis* histones were expressed in *E. coli*, isolated from inclusion bodies, and purified under denaturing conditions using sequential size-exclusion (Superdex 200 10/30 GL, Cytiva) and cation-exchange chromatography (XK26/20 SP-Sepharose HP, Cytiva). Purified histones were dialyzed into reducing buffer, lyophilized, and stored at −80°C.

### Site-specific histone modification by native chemical ligation

Site-specific H3K9 trimethylation was introduced using native chemical ligation (NCL) as described before [31] with some modifications. A recombinant C-terminal H3 fragment (residues 21–135) engineered to present a cysteine at the ligation junction (see Table S2) was ligated to a synthetic N-terminal H3 peptide bearing the desired K9 trimethyl modification and a C-terminal thioester. Ligation was performed in denaturing NCL buffer (100 mM potassium phosphate pH ∼7.9, 6 M guanidinium–HCl, TCEP, 20 mM TCEP, 50 mM 4-MPAA) at 20°C. The ligation product was purified by cation-exchange chromatography (HiTrap SP-Sepharose HP, Cytiva), desalted, dialyzed, lyophilized, and stored at −80°C. Product identity was confirmed by SDS–PAGE.

### Fluorophore labelling of histones

Fluorophore labelling was performed as described [32]. Lyophilized *Xenopus laevis* histone H2B containing an engineered cysteine at position 1 (H2B A1C) was dissolved in labelling buffer (20 mM Tris–HCl, pH 7.5, 6 M guanidinium–HCl, 5 mM EDTA, 0.7 mM TCEP). Dylight 405-C5-maleimide or Dylight 680-C5-maleimide (Thermo Fisher) resuspended in dimethylformamide (DMF) was added at a histone-to-dye molar ratio of 1:5, and reactions were incubated for 12 h at room temperature in the dark with continuous stirring. Excess dye was removed using Pierce™ Dye Removal Columns, and labelled histones were dialyzed into water. Protein concentration and labelling efficiency were determined spectrophotometrically by absorbance at 280 nm and 405/680 nm, following manufacturer’s guidelines. Labelled histones were aliquoted, lyophilized, and stored at –80°C for use in histone octamer assembly.

### Assembly and purification of histone octamers

Histone octamers were assembled from H2A, H2B (labelled or unlabelled), H3 (wild-type or modified), and H4 as described [13, 33, 34]. Lyophilized histones were unfolded in denaturant and combined at stoichiometric ratios (typical H2A:H2B:H3:H4 ≈ 1.2:1.2:1:1). The mixture was refolded by dialysis into high-salt refolding buffer (10 mM Tris–HCl pH 7.5, 2 M NaCl, 1 mM EDTA, 1 mM DTT) with several buffer exchanges at 4°C. Reconstituted octamers were concentrated and further purified by size-exclusion chromatography (Superdex 200 10/30 GL, Cytiva) to separate assembled octamers from H2A–H2B dimers. Pure octamer fractions as determined by SDS-PAGE and Coomassie staining were pooled, concentrated, and stored in 50% (v/v) glycerol at −20°C.

### Reconstitution of 12-mer chromatin arrays

Chromatin arrays were reconstituted by gradient salt-dialysis using purified histone octamers and 12×200-601 DNA templates as described [34]. For fluorescent arrays, labelled octamers were mixed into the octamer pool at a defined labelled:unlabelled ratio (typically 1:20). The reaction mixture of DNA and octamers was set up at high ionic strength (≥2 M NaCl) and then subjected to continuous dialysis toward low salt to promote nucleosome formation. The degree of nucleosome occupancy (expressed as *r*, octamers per DNA repeat) was optimized in small-scale test reactions (e.g., *r* ≈ 0.9–1.1) before scale-up. Assembled arrays were assessed by native agarose gel electrophoresis to verify quality and stored at 4°C; concentrations were calculated from DNA absorbance at 260 nm.

### Histone peptide and chromatin array pull-down assays

Pull-down assays were performed using either biotinylated histone peptides (Table S2) or biotinylated chromatin arrays as previously described [13]. Peptides (5 µg) or chromatin arrays (1 µg) were immobilized on 50 µL of streptavidin-coated magnetic beads (Thermo Fisher) in binding buffer (10 mM triethanolamine–HCl pH 7.5, 150 mM NaCl, 0.1% (v/v) Triton X-100, 5% (v/v) glycerol, 0.1 mM EDTA) for 4 h at 4°C. Beads were washed twice with the same buffer and incubated for 1 h at 4°C with recombinant HPL-2 mutants. After binding, beads were washed with buffer containing 300 mM NaCl and bound proteins were eluted by boiling in SDS sample buffer. Eluted material was analysed by SDS–PAGE and Coomassie staining.

### Mass photometry

Mass photometry was performed using a Refeyn OneMP instrument as described [35]. Measurements were carried out in 20 mM HEPES–NaOH pH 7.5, 150 mM NaCl, with protein concentrations of 50–75 nM. Data were acquired for 60 s and analysed using the DiscoverMP software package (Refeyn). Molecular mass calibration was performed using bovine serum albumin (monomer 66 kDa, dimer 132 kDa) and apoferritin (24-mer, 480 kDa).

### Preparation of mPEGylated microscopy plates for phase separation assays

Microscopy experiments were conducted in 384-well SensoPlates™ (Greiner Bio-One) functionalized with mPEG-silane (MW 5000; Laysan Bio) as described [32, 36]. Briefly, wells were cleaned, etched with 1 M NaOH, and incubated overnight with 20 mg/mL mPEG-silane in 95% (v/v) ethanol. Before imaging, wells were passivated with 100 mg/mL BSA for 30 min at room temperature, rinsed, and loaded immediately with 20 µL of sample. Samples were sealed to minimize evaporation.

### Disorder prediction

Intrinsic disorder across HP1 hinge regions was assessed using IUPred3 [37]. Disorder scores were normalized to sequence length, and residues with scores ≥0.5 were classified as disordered. Sequence physicochemical properties associated with intrinsically disordered regions were further analysed using the CIDER server [38]. Canonical hinge-region motifs associated with phase separation, including RKR, KRK were identified using Python scripts implemented in Google Colab.

### Protein-only LLPS

LLPS of recombinant HPL-2 mutants was examined following buffer exchange by dialysis (3.5 kD MWCO dialysis tubing, Spectra/Por) into PS150 buffer (20 mM HEPES–NaOH pH 8.0, 150 mM NaCl, 1 mM TCEP). Proteins were concentrated to twice the final assay concentration and mixed 1:1 (v/v) with PS150 buffer containing 20% (w/v) PEG 8000 to yield final reactions containing 10% (w/v) PEG 8000. Samples (20 µL) were transferred to PEGylated wells and incubated on ice for at least 60 min before imaging.

### Chromatin condensation assays

Fluorescently labelled 12-mer chromatin arrays and Alexa Fluor 555–labelled or unlabelled HPL-2 proteins were dialyzed into PS0 buffer (20 mM HEPES–NaOH pH 8.0, 1 mM TCEP). Trace labelling was achieved by mixing labelled and unlabelled protein at a 1:10 molar ratio. Reactions (20 µL) contained 1 µg of H3K9me0 or H3K9me3 chromatin arrays, HPL-2 at the indicated concentration, and 1x PS50 buffer (20 mM HEPES–NaOH pH 8.0, 50 mM NaCl, 1 mM TCEP). Samples were incubated for 1 h at room temperature prior to imaging.

For testing chromatin enrichment under LLPS-promoting conditions, Alexa Fluor 555-labelled HPL-2 proteins were first incubated in the presence of 10% (w/v) PEG8000 in PS150 buffer for 1 h on ice to allow condensate formation. Fluorescently labelled H3K9me0 or H3K9me3 chromatin arrays (1 µg) were then added to the preformed HPL-2 condensates, followed by fluorescence imaging.

### Chromatin segregation assays

Fluorescently labelled 12-mer chromatin arrays and Alexa Fluor 555–labelled or unlabelled HPL-2 proteins were dialyzed into PS0 buffer (20 mM HEPES–NaOH pH 8.0, 1 mM TCEP). Trace labelling was achieved by mixing labelled and unlabelled protein at a 1:10 molar ratio. Segregation assays were performed using equal amounts of fluorescently labelled H3K9me0 and H3K9me3 arrays. Reactions (20 µL) contained 1 µg each of H3K9me0 and H3K9me3 chromatin arrays, 100 µM HPL-2 mutants and 1x segregation buffer (25 mM Tris–OAc pH 7.5, 1 mM Mg(OAc)₂, 1 mM EDTA, 5 mM DTT, 150 mM KOAc). Samples were incubated for 1 h at room temperature before imaging.

### Imaging of condensates

LLPS, chromatin condensation, and segregation assays were imaged at the KAUST Imaging and Characterization Core Lab. Protein-only condensates were visualized in DIC mode using a Leica THUNDER microscope equipped with a 63x oil objective. Chromatin-based assays were imaged on a Leica SP8 confocal microscope equipped with a 63x oil objective (512 x 512 pixels, scan speed 400 Hz, zoom 3, line averaging 4, laser power ∼1%). All imaging was performed at room temperature.

### Turbidity assay

Turbidity measurements were done to quantify LLPS of proteins or chromatin arrays. After inducing phase separation for 1 h, 25 µL of each sample was transferred into a clear 384-well plate, and absorbance at 600 nm (A₆₀₀) was measured using an Infinite® M1000 plate reader (Tecan). Each assay was performed in at least three independent replicates, and data were analysed using GraphPad Prism.

### *C. elegans* maintenance and strain generation

*C. elegans* were cultured on nematode growth medium (NGM) plates seeded with *E. coli* OP50-1 under standard conditions at 20°C unless otherwise indicated [39]. Strains used in this study are listed in Supplementary Table S3. Endogenously GFP-tagged *hpl-2* mutant strains were generated by SunyBiotech (Fuzhou, China) using CRISPR/Cas9-mediated editing. Strain genotypes were routinely confirmed by single-worm PCR using primers listed in Table S4.

### Brood size assays

Brood size assays were performed at 20°C or 25°C as described [40]. Individual L4 hermaphrodites (n = 40 per condition) were transferred daily to fresh plates until egg laying ceased. The number of live progenies, unhatched embryos, and sterile hermaphrodites was recorded to calculate embryonic viability, sterility, and mean brood size. Data were averaged across four independent experiments.

### Lifespan assays

Lifespan assays of *C. elegans* were performed at 20°C and 25°C as described [41, 42]. Briefly, NGM plates were seeded with *E. coli* OP50-1, without 5-fluoro-2′-deoxyuridine (FUDR). For each strain, 75 worms (15 per plate) were monitored in three independent experiments. Survival was scored every 1–3 days; animals unresponsive to touch were considered dead, and censored worms (lost or escaped) were excluded. Kaplan–Meier survival curves and statistical analyses were performed using OASIS [43].

### High-temperature arrest assay

High-temperature arrest (HTA) assays of *C. elegans* were performed essentially as described [44]. L4 hermaphrodites of each genotype were transferred to either 24°C or 26°C and incubated overnight. Three independent plates per genotype, each containing 9–12 adults, were then established and adults were allowed to lay eggs for 4–5 h before removal. Progeny were maintained at the same temperature and scored two days later as either arrested larvae or adults. Data from three independent biological experiments were pooled, and the percentage of arrested larvae was calculated from the total number of progenies scored.

### Embryo preparation and immunofluorescence

Gravid *C. elegans* adults were treated with alkaline hypochlorite to isolate embryos [39]. Immunofluorescence staining of embryos was performed as described [45, 46] with minor modifications. Briefly, embryos were freeze-cracked, fixed sequentially in methanol (30 s) and 1% (v/v) formaldehyde (2 min), and washed in PBST (PBS + 0.5% (v/v) Triton X-100). Samples were blocked with 1% (w/v) BSA in PBST and incubated overnight at 4°C with primary antibodies, followed by fluorescent secondary antibodies for 4 h at 4°C. Embryos were mounted in antifade reagent for fluorescence imaging. Details of antibodies used are listed in Supplementary Table S5.

### Confocal fluorescence microscopy

Live or fixed *C. elegans* embryos were mounted on agarose pads and imaged using a Leica SP8 confocal microscope (63x oil objective) at the KAUST Imaging and Characterization Core Lab. Typical settings were: 512 x 512 pixels, 400 Hz scan speed, zoom = 3, line averaging = 4, laser power ≈ 1%, and Z-step = 0.3 µm. Nuclear foci were quantified in Fiji [47] using Otsu thresholding and watershed segmentation, and the number and total area of foci per nucleus were measured from binary masks.

### Fluorescence recovery after photobleaching (FRAP)

FRAP experiments on *C. elegans* embryos were conducted on a Zeiss LSM 880 confocal light scanning microscope equipped with a 63x oil immersion objective. Embryos were placed on a chambered LabTek coverslip and were immobilized with an agarose pad. FRAP videos were typically acquired at 128 × 256 pixels and at a scan speed corresponding to 250 milliseconds per image for 20 – 30 seconds. For each experiment, a custom script written in Python was used to segment cell nuclei and HPL-2 foci to retrieve the average intensity of the bleached foci (*I*_B_), the entire cell nucleus (*I*_REF_) and the background of the image (*I*_BG_). The intensity of the entire cell nucleus was used as an internal reference to quantify unwanted acquisition photobleaching. The FRAP curves were calculated according to:

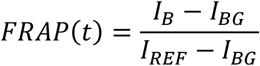

Subsequently, FRAP curves were double normalized according to:

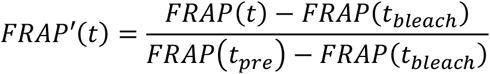

with *t*_bleach_ and *t*_pre_ representing the acquisition times of the first post-bleach and the last pre-bleach frame, respectively. Finally, FRAP curves were fitted with a diffusion model, yielding immobile fractions and apparent diffusion times [48].

### Chromatin immunoprecipitation (ChIP)

Chromatin immunoprecipitation (ChIP) experiments were performed on three independent biological replicates of mixed-stage *C. elegans* embryos for each strain, adopting a previously described protocol [49].

Gravid adult *C. elegans* were grown on NGM plates, and embryos were isolated by standard hypochlorite bleaching. Embryos were crosslinked in 2% (w/v) formaldehyde for 30 min at room temperature with gentle rotation, and the reaction was quenched by addition of glycine to a final concentration of 125 mM for 5 min. Crosslinked embryos were washed extensively with M9 buffer followed by PBS containing protease inhibitors and 1 mM PMSF, snap-frozen in liquid nitrogen, and stored at −80°C until use. Frozen embryo pellets were resuspended in FA buffer (50 mM HEPES-KOH, pH 7.5, 150 mM NaCl, 1 mM EDTA, 1% (v/v) Triton X-100, and 0.1% (w/v) sodium deoxycholate) supplemented with protease inhibitors. Embryos were disrupted on ice using a glass Dounce homogenizer (approximately 30 strokes with a type B pestle), and chromatin was fragmented to an average size of 200–800 bp using a Covaris sonicator according to the manufacturer’s recommended settings. Insoluble material was removed by centrifugation, and the chromatin concentration was determined using a Bradford assay.

For each immunoprecipitation, 1–2 mg isolated chromatin was diluted to 500 μL in FA buffer. Five percent of each sample was reserved as input. Chromatin extracts were pre-cleared with Protein A magnetic beads before overnight incubation at 4°C with antibody-bound Protein A magnetic beads. Details of antibodies used are listed in Supplementary Table S5. Beads were washed sequentially with FA buffer, FA buffer containing 1 M NaCl, FA buffer containing 500 mM NaCl, TEL buffer (0.25 M LiCl, 1% (v/v) IGEPAL CA-630, 1% (w/v) sodium deoxycholate, 1 mM EDTA, and 10 mM Tris-HCl, pH 8.0), and TE buffer. Chromatin was eluted in ChIP elution buffer (1% (v/v) SDS, 250 mM NaCl, 10 mM Tris-HCl, pH 8.0, and 1 mM EDTA), treated sequentially with RNase A and Proteinase K, reverse crosslinked overnight at 65°C, and purified using the ChIP DNA Clean & Concentrator Kit (Zymo Research).

### ChIP-qPCR

ChIP-qPCR was performed essentially as described [49]. To assess enrichment across heterochromatic and non-heterochromatic regions, primer pairs were designed for fifteen target loci and five off-target loci distributed across the five autosomes. For each autosome, three target loci and one off-target locus were selected based on previously published HPL-2 and H3K9me2 ChIP-seq datasets generated from *C. elegans* embryos (GEO accession GSE271919). Target loci were selected from autosomal arm regions exhibiting clear overlapping enrichment of both HPL-2 and H3K9me2, whereas off-target loci were selected from central chromosomal regions lacking detectable enrichment for either HPL-2 or H3K9me2. This strategy ensured that positive-control loci represented canonical HPL-2/H3K9me2-associated heterochromatin, while negative-control loci represented genomic regions devoid of HPL-2 and H3K9me2 enrichment. The genomic coordinates of the analysed loci and primers used are listed in Supplementary Table S6.

### ChIP-seq

Purified ChIP and input DNA were used for library preparation with the NEBNext® Ultra™ II DNA Library Prep Kit (Illumina®). Libraries were sequenced by Novogene (Beijing, China) on an Illumina NovaSeq X Plus platform to generate paired-end 150-bp reads, yielding approximately 15 million read pairs per sample.

### Bioinformatic analysis of ChIP-seq

Fastq reads were aligned to the WBcel235 version of the *C. elegans* genome using the nf-core/chipseq pipeline, v2.1.0, with the following optional parameters specified: ‘--read_length 150 --min_reps_consensus 3 --macs_fdr 0.05’. Log2(IP/input) tracks were created using deeptools bigwigCompare with ‘--operation log2 --binSize 10’ paremeters specified and then averaged with deeptools bigwigAverage with ‘-bs 10’ and ce11 blacklisted regions were obtained from the Boyle lab https://github.com/Boyle-Lab/Blacklist/blob/master/lists/. Consensus broad peaks found in at least three H3K9me2 chipseq samples were obtained from the nf-core/chipseq pipeline. Scripts for processing and plotting the data can be found on https://doi.org/10.5281/zenodo.22099313.

### RNA isolation and sequencing

Total RNA was isolated from frozen mixed-stage *C. elegans* embryo pellets using the GenElute™ Mammalian Total RNA Miniprep Kit (Sigma-Aldrich). RNA quality was confirmed by spectrophotometry (A260/280 = 1.8–2.2, A260/230 ≥ 2.0). Directional RNA-seq libraries were prepared after rRNA depletion and sequenced by Novogene (Beijing, China) on an Illumina NovaSeq X Plus platform (paired-end 150 bp, ∼40 million reads per sample, three biological replicates per strain).

### Bioinformatic analysis of RNA-seq

A GTF for *C. elegans* canonical genes version WS298 was downloaded from Wormbase and merged with a gtf created from annotations of repeat sequences downloaded from Dfam v3.5. Reads were aligned to the *C. elegans* genome using the nf-core [50] RNA-seq pipeline v 3.26.0 with the --extra_star_align_args ‘--outFilterMultimapNmax 100 -- winAnchorMultimapNmax 100 --alignIntronMax 100000 --outFilterMismatchNoverLmax 0.04’ option specified. For analysing canonical gene expression all Dfam repeat counts, and counts for genes overlapping rRNA clusters (I:15060299–15071033, V:17115526–17132107) or signal recognition particle RNAs (*srpr-1.1–3*) were excluded from count matrices. Differential expression analysis was performed using the nf-core differential abundance pipeline v 2.0.0 with DESeq2 shrinkage enabled (--deseq2_shrink_lfc true) and filtering thresholds specified (--filtering_min_abundance 5, --filtering_min_samples 3). Genes with adjusted *p* < 0.05 and |log₂ fold change| > 0.5 were considered significantly differentially expressed.

Plots were generated in R v4.5.2 using RStudio v2026.07.1, and ChIP–RNA integration heatmaps were produced using deepTools v3.5.6. Custom scripts are available at https://github.com/CellFateNucOrg/FischleLab_KarthikEswara_ribo0seq_repeats/. Gene arm definitions were as described [51], with both arm and tip genes classified as “arm” genes.

Genes overlapping H3K9me2, HPL-2, or LIN-61 ChIP peaks were defined as those whose gene bodies overlapped at least one peak in ChIPseq performed in mixed-stage N2 embryos. Processed peaks and bigwigs for this data were downloaded from GSE271919.

Differential gene expression of Dfam repeats was performed using a repeat-aware pipeline called Telescope that uses an expectation maximisation algorithm to assign reads to repeats [52]. Bam files produced by STAR in the nf-core/rnaseq pipeline were used as input for Telescope. The results from Telescope were then aggregated from individual genomic loci to repeat family level counts and differential expression was performed with DESeq2. Scripts for performing quantification, and plotting are available: https://doi.org/10.5281/zenodo.22099356.

## RESULTS

### HPL-2 as a model factor for studying HP1-dependent chromatin condensation

HPL-2 is one of the two *C. elegans* HP1 orthologues [22]. Several studies have established key HP1-like properties of HPL-2, including conservation of the chromodomain (CD) and chromoshadow domain (CSD) [22, 24], binding to H3K9-methylated peptides [21, 26, 53, 54], and homodimerization [24]. Since previous sequence comparisons of HPL-2 were limited to HP1 proteins from *C. elegans*, humans and *Drosophila* [22, 24], we performed domain-wise sequence alignments with diverse HP1 orthologues of a broad phylogenetic range (Figure S1A). This analysis revealed extensive conservation across the CD and CSD, including the aromatic cage residues required for H3K9me recognition and the conserved isoleucine residue implicated in CSD-mediated dimerization. Hydrophobicity-based similarity scoring further revealed conservation of the physicochemical character of these domains (Figure S1B). AlphaFold modelling of dimeric HPL-2 predicted a canonical CD aromatic cage (F19, W41, F44) that creates a hydrophobic pocket putatively capable of accommodating the methylated lysine of H3K9me (Figure S2A, bottom inset) [55]. The predicted structure closely matched the human HP1α–H3K9me3 complex (RMSD 0.323 Å; Figure S2B).

We next asked whether the H3K9me-binding activity of HPL-2 extends to the context of chromatin substrates. To address this, we reconstituted chromatin arrays carrying defined histone H3 methylation states (Figure S3, S4) and performed pull-down assays. The wild-type HPL-2 protein bound strongly to H3K9me3-modified arrays, whereas mutation of the aromatic cage residues (ACM; F19A, W41A, F44A) caused markedly reduced binding (Figure 1A; Figure S5A). Quantification revealed ∼3-fold enrichment of wild-type HPL-2 on H3K9me3 arrays relative to unmodified arrays, consistent with aromatic cage–dependent recognition of methylated histone tails (Figure 1A). HPL-2 exhibited comparable binding to H3K9me2 and H3K9me3 but showed markedly reduced affinity for unmodified H3K9me0 peptides (Figure S5B), suggesting a preference for methylated substrates without discrimination between di- and tri-methyl states.

**Figure 1.**
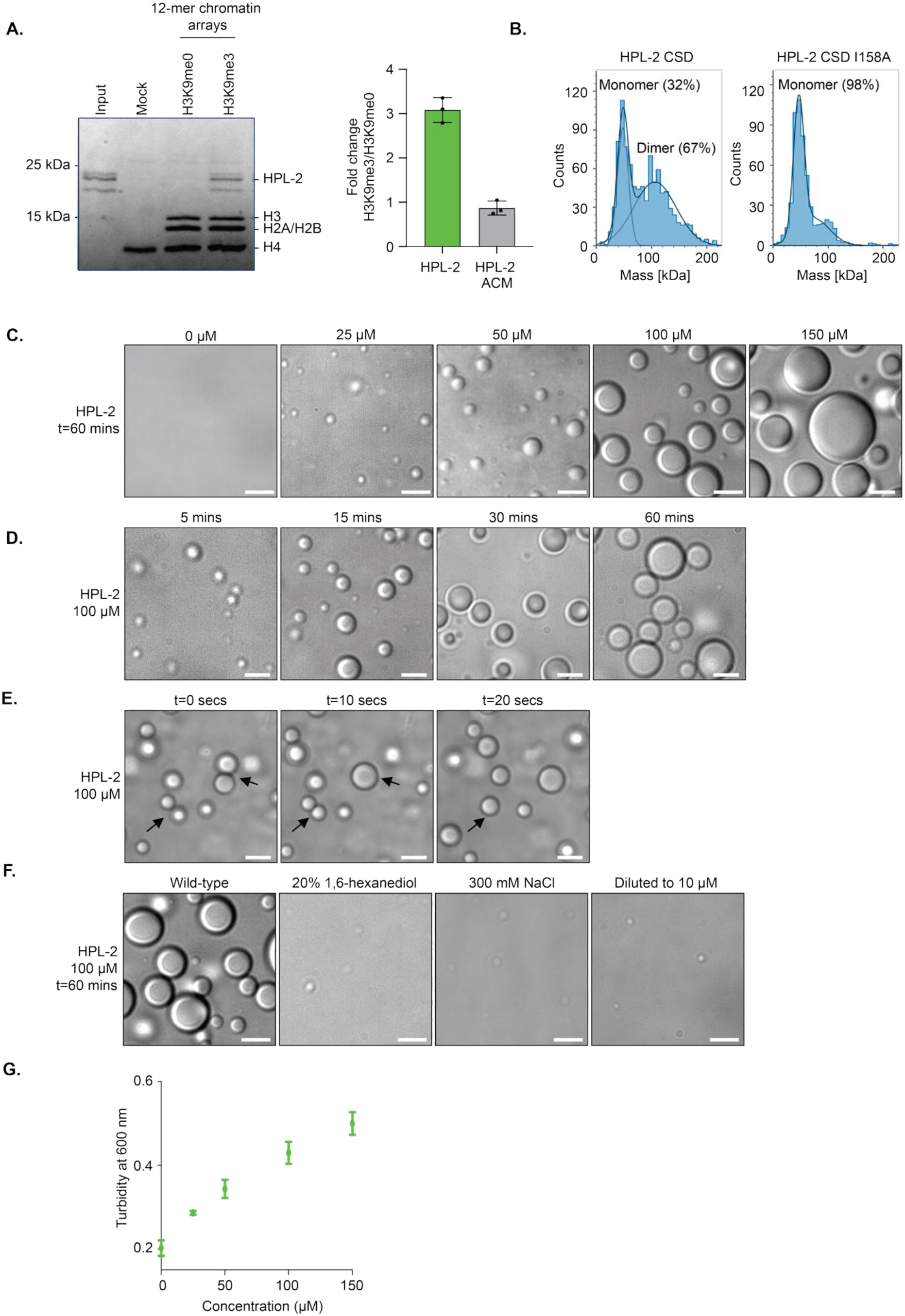
Recombinant *C. elegans* HPL-2 binds H3K9me, homodimerizes and undergoes liquid–liquid phase separation (LLPS) **(A)** Left: Recombinant HPL-2 was incubated with biotinylated 12-mer H3K9me0 or H3K9me3 chromatin arrays immobilized on streptavidin beads. Bound proteins were analysed by SDS– PAGE and stained with Coomassie blue. Positions of molecular weight markers, histones, and HPL-2 are indicated. “Mock” indicates control pull-downs with streptavidin beads to assess non-specific binding; “Input” represents 10% of total HPL-2 used. Right: Quantification of chromatin array pulldowns as shown on the left normalized to histones H2A/H2B. Data represent mean ± SD from three independent experiments. ACM, aromatic cage mutant. **(B)** Mass photometry analysis of recombinant MBP-tagged CSDs of wild-type and mutant HPL-2 proteins. Particle counts (y-axis) are plotted as a function of molecular mass (x-axis). Data are representative of three independent experiments. **(C–F)** Differential interference contrast (DIC) microscopy of HPL-2 condensates. Data are representative of three independent experiments. Scale bars: 5 μm. (C) Representative images of condensate formation at different protein concentrations. (D) Time course of condensate formation at 100 μM protein concentration. (E) Fusion dynamics of HPL-2 condensates. Arrows indicate individual fusion events. Time elapsed between fusion events is shown. (F) Effect of 20% (v/v) 1,6-hexanediol (left), 300 mM NaCl (middle), or dilution to 10 μM HPL-2 (right) on condensate stability. **(G)** Plot of turbidity measured by absorbance reading at 600 nm against different HPL-2 protein concentrations. Condensate formation was induced as in (C). Data represent mean ± SD from three independent replicates.

We next examined CSD-mediated dimerization of HPL-2. AlphaFold modelling predicted a dimeric HPL-2 CSD, structurally similar to the human HP1α CSD (RMSD 0.975 Å; Figure S2A top inset, S6A), with a conserved isoleucine residue (I158) positioned within the dimerization interface [56]. Mass photometry confirmed that wild-type HPL-2 CSD at 50-75 nM concentration exists in a dynamic monomer–dimer equilibrium, with 67% dimers and 32% monomers. At the same concentrations and conditions, the corresponding HPL-2 CSD I158A mutant protein, in contrast, was almost exclusively monomeric (Figure 1B). Using the same analysis method, the human HP1β CSD was found to predominantly form dimers (97%), while mutation of the I161 residue (HP1β CSD I161A) disrupted this interaction (Figure S6B) [57]. In chromatin array pulldown experiments, the full length HPL-2 I158A protein retained normal binding to H3K9me3 chromatin (Figure S6C,D), indicating that dimerization is dispensable for CD-mediated recognition of the histone modification *in vitro*.

Because LLPS is a key feature of HP1 proteins [14, 15], we next assessed whether HPL-2 undergoes phase separation. LLPS is often driven by weak multivalent interactions among charged residues within intrinsically disordered regions (IDRs) [58]. Consistent with this, the sequence of the HPL-2 hinge region exhibits high disorder propensity (IUPred > 0.5) and a high proportion of charged residues, like HP1 orthologues across fungi, insects, nematodes and vertebrates (Figure S7A,B). To further compare the biophysical properties of these disordered hinge regions, we analysed them using CIDER (Classification of Intrinsically Disordered Ensemble Regions) [38], which classifies IDRs based on sequence features that influence their conformational behaviour. All HP1 hinge regions analysed, except *C. elegans* HPL-1, were classified as strong polyampholytes, consistent with broadly similar sequence-ensemble properties despite extensive sequence divergence (Figure S7C). Finally, because a positively charged KRK motif has previously been implicated in HP1α phase separation [14], we examined the occurrence of KRK and the closely related RKR motif across HP1 orthologues. Although the exact motif sequence is not conserved, all proteins analysed contain at least one of these polybasic motifs (Figure S7D), suggesting that conservation of the hinge region primarily occurs at the level of physicochemical properties rather than primary amino acid sequence.

To assess whether HPL-2 undergoes LLPS *in vitro*, purified recombinant protein was incubated with a molecular crowding agent (10% (w/v) PEG8000) in a buffer containing 150 mM NaCl. Under these conditions, HPL-2 formed micron-sized, spherical condensates in a concentration-dependent manner, visualized by DIC microscopy (Figure 1C). The resulting condensates displayed hallmarks of liquid-like behaviour [59], including gradual growth and fusion over time (Figure 1D,E), and dissolved completely upon dilution (Figure 1F). HPL-2 LLPS was disrupted by treatment with 1,6-hexanediol (20% (v/v); Figure 1F), an aliphatic alcohol that interferes with weak hydrophobic and cation–π interactions required to stabilise liquid condensates [35]. LLPS was also sensitive to salt concentration, with impaired HPL-2 condensate formation observed at increased ionic strength (300 mM NaCl; Figure 1F). Consistent with DIC microscopy showing progressively larger and more abundant condensates at higher HPL-2 concentrations (Figure 1C), turbidity measurements exhibited a concentration-dependent increase in light scattering under the same experimental conditions (Figure 1G).

Together, our findings demonstrate that *C. elegans* HPL-2 possesses the defining molecular properties of HP1 proteins *in vitro*: specific recognition of H3K9me2/3 chromatin via the CD, homodimerization through the CSD, and intrinsic ability to undergo LLPS. It is thus a tractable model for dissecting the molecular working mode of HP1 proteins in heterochromatin condensation.

### Dimerization and disordered hinge region are nonredundant in HPL-2 LLPS

Having established that HPL-2 undergoes LLPS, we next dissected the molecular features driving this behaviour, focusing on the intrinsically disordered hinge region and the CSD– mediated dimerization (Figure 2A). We generated a series of mutants targeting the hinge region: completely deleting the hinge region (Δhinge), scrambling its sequence but preserving the amino acid composition (scrambled hinge), neutralizing either the conserved RKR motif (RKR neutral) [14] or a cluster of nine charged residues (9-patch neutral), fully neutralizing all its charged residues (total neutral) and two variants in which the RKR motif was repositioned within the hinge region (RKR central and RKR right) (Figure S8A,B). IUPred analysis showed that all mutants retained a similar overall disorder propensity as the wild-type HPL-2 protein (Figure S8C).

**Figure 2.**
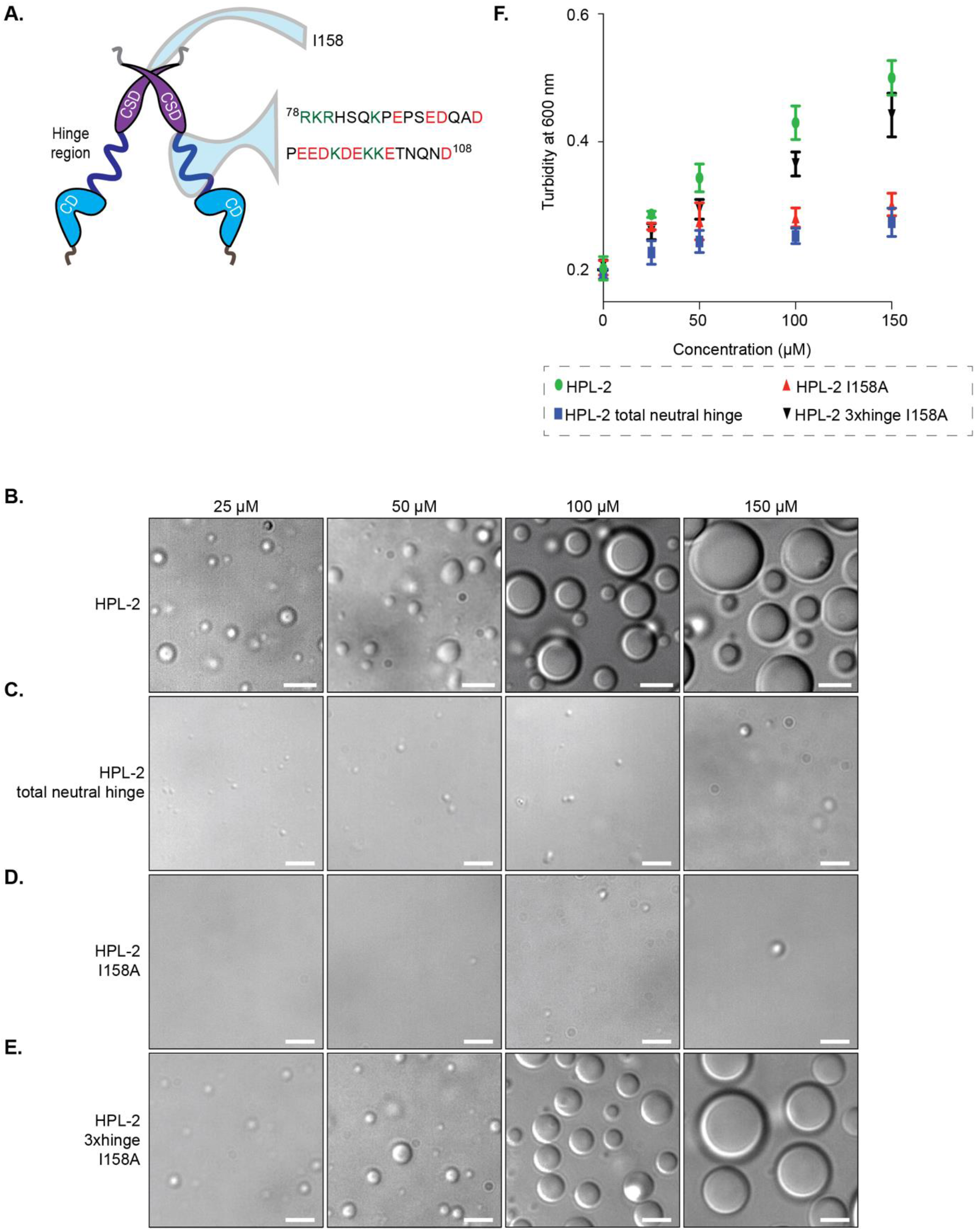
Molecular determinants of HPL-2 phase separation *in vitro*. (A) Schematic representation of dimeric HPL-2. The inset in the hinge region shows the amino acid sequence, with positively charged residues highlighted in green and negatively charged residues in red (at pH 7.4). The inset in the chromoshadow domain (CSD) shows a magnified view of an isoleucine residue (I158) critical for dimerization. **(B–E)** Differential interference contrast (DIC) microscopy of condensates formed by wild-type HPL-2 (B) or the indicated mutant proteins (C–E) at increasing protein concentrations. Images are representative of three independent experiments. Scale bars: 5 μm. **(F)** Plot of turbidity measured by absorbance reading at 600 nm against increasing concentrations of wild-type and mutant HPL-2 proteins. Condensate formation was induced as in (B–E). Data represent mean ± SD from three independent replicates.

We then examined phase separation of the HPL-2 mutant proteins using microscopy and turbidity assays and found that these could be broadly grouped by their LLPS behaviour. The HPL-2 9-patch neutral hinge mutant formed condensates comparable to the wild-type protein, indicating that this charged cluster is dispensable for LLPS (Figure S9A,G). In contrast to the wild-type protein, both the HPL-2 Δhinge and HPL-2 RKR neutral mutants formed smaller, less dynamic condensates with impaired fusion and reduced turbidity (Figure S9B,C,G), suggesting that the RKR motif is critical for maintaining condensate fluidity. The HPL-2 scrambled hinge and HPL-2 total neutral hinge mutant proteins had the most severe defects, forming sparse condensates even at high concentrations (Figure 2C; Figure S9D,G). Positioning the RKR motif to the centre of the hinge region abolished condensate formation, whereas relocation to the C-terminal end preserved phase separation comparable to the wild-type protein (Figure S9E–G). Together, these results indicate that HPL-2 LLPS depends not simply on hinge charge, amino-acid composition, or overall disorder propensity, but on the sequence patterning and positional context of charged motifs within the hinge region.

We next examined the contribution of CSD-mediated dimerization to HPL-2 LLPS. The dimerization-deficient HPL-2 I158A mutant protein failed to form condensates at any concentration tested, as indicated by microscopy and turbidity measurements (Figure 2D,F). To directly test whether dimerization is required for LLPS, we artificially restored it using the SpyTag–SpyCatcher system (Figure S10A,B) [60]. The forced dimerization fully rescued LLPS and even enhanced condensate formation beyond wild-type HPL-2 levels (Figure S10D,F). Since proteolytic cleavage of the enforced dimer abolished LLPS (Figure S10E,F), the findings establish that dimerization is necessary and, when artificially established, sufficient to drive HPL-2 condensate formation.

As both the hinge region and dimerization contribute to HPL-2 LLPS, we next asked whether loss of dimerization could be compensated by enhancing disorder content. In the background of the dimerization-deficient I158A mutation, we elongated the HPL-2 hinge region or replaced it with heterologous IDRs previously shown to drive LLPS in their native contexts—*Arabidopsis* H2B.8 [61] and human DDX4 [62] (Figure S11A,B). Duplication of the hinge region (2xhinge I158A) resulted in weak rescue, with protein condensates forming only at high concentrations (Figure S11C,F), while its triplication (3xhinge I158A) robustly restored LLPS to near wild-type levels (Figure 2E,F). The results demonstrate that increased disorder content can compensate for the loss of dimerization. Substitution of the HPL-2 hinge region with heterologous IDRs yielded distinct outcomes: the H2B.8 IDR supported condensate formation but produced irregular, solid-like droplets (Figure S11D,F), whereas the DDX4 IDR markedly enhanced LLPS, lowering the concentration threshold for droplet formation (Figure S11E).

Our findings establish that the disordered hinge region and CSD-mediated dimerization are nonredundant molecular determinants of HPL-2 phase separation. While the hinge region provides multivalent, charge-based interactions that modulate condensate dynamics, dimerization enhances intermolecular connectivity and phase separation efficiency. Importantly, we achieved uncoupling of LLPS and dimerization with the HPL-2 total neutral hinge mutant completely abolishing LLPS despite an intact dimerization interface and the HPL-2 3xhinge I158A mutant showing LLPS in absence of dimerization.

### HPL-2 dimerization mediates condensation of H3K9me3 chromatin arrays

Heterochromatin organisation relies on chromatin compaction within the nucleus [63]. Our biochemical analyses identified and mutationally isolated two key properties of HPL-2, dimerization and LLPS. To determine if these functionalities enable HPL-2 to promote chromatin condensation, we established an *in vitro* chromatin condensation assay using defined 12-mer chromatin arrays (Figure S3, S4). In this assay, fluorescently labelled chromatin arrays, uniformly containing H3K9me0 or H3K9me3, were incubated with increasing concentrations of fluorescently labelled HPL-2 (Figure 3A). Turbidity measurement and fluorescence microscopy were used to monitor condensate formation. Control reactions confirmed that neither HPL-2 alone, even at 100 µM, nor chromatin arrays alone formed condensates under the applied conditions (Figure S12A–C). Therefore, any condensation observed in this system reflects specific chromatin effects of HPL-2.

**Figure 3.**
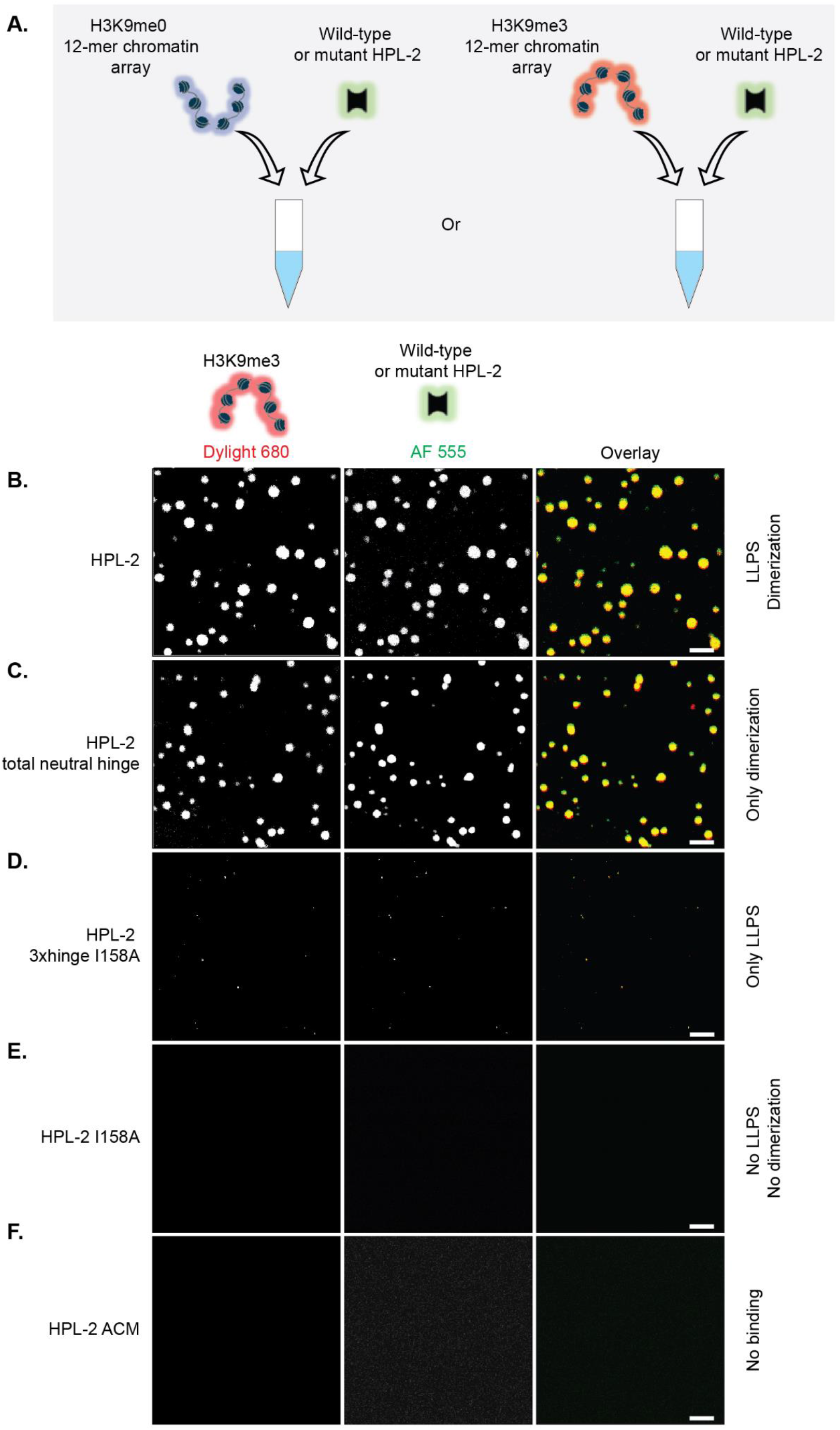
HPL-2 dimerization drives H3K9me3 chromatin condensation *in vitro*. **(A)** Scheme of the chromatin condensation assay. Fluorescently labelled chromatin arrays are incubated with increasing concentrations of fluorescently labelled HPL-2. Phase separation is monitored by turbidity measurements and fluorescence microscopy. **(B–F)** Fluorescence microscopy of Dylight 680–labelled H3K9me3 12-mer chromatin arrays in presence of wild-type (B) or the indicated HPL-2 mutant proteins (C–F) trace-labelled with Alexa Fluor 555– HPL-2. Scale bars: 5 μm. Images are representative of three independent experiments.

First, we titrated HPL-2 over a range of concentrations in the presence of constant concentrations of H3K9me0 or H3K9me3 chromatin arrays. Whereas H3K9me3 chromatin arrays displayed a robust, HPL-2-dependent increase in turbidity, only minimal effects were recorded for H3K9me0 chromatin arrays (Figure S12B,C). Fluorescence imaging confirmed the formation of spherical condensates containing both HPL-2 (AF555) and H3K9me3 chromatin (DyLight 680), indicative of co-partitioning into the same phase (Figure 3B). These results demonstrate that HPL-2 alone is sufficient to drive condensation of reconstituted chromatin arrays *in vitro*, with a strong preference for H3K9me-marked substrates.

We next examined how LLPS and dimerization individually contribute to chromatin condensation. To this end, we compared wild-type HPL-2 with different mutants: HPL-2 total neutral hinge (dimerization-competent but LLPS-deficient), HPL-2 3xhinge I158A (LLPS-competent but dimerization-deficient), HPL-2 I158A (defective in LLPS and dimerization), and HPL-2 ACM (deficient in H3K9me binding). Under the same assay conditions, none of these proteins formed condensates in the absence of chromatin (Figure S12A). As expected, the HPL-2 ACM protein failed to condense either H3K9me0 or H3K9me3 arrays (Figure 3F; Figure S12B,C), confirming that recognition of H3K9me marks is essential for chromatin condensation. The HPL-2 total neutral hinge mutant protein condensed H3K9me3 but not H3K9me0 arrays with slightly reduced efficiency relative to the wild-type protein (Figure 3C; Figure S12B,C), indicating that dimerization alone can drive H3K9me3-dependent chromatin condensation. In contrast, the HPL-2 3xhinge I158A and HPL-2 I158A mutant proteins were completely inactive, failing to condense either modified or unmodified chromatin arrays (Figure 3D,E; Figure S12B,C).

Because LLPS of HPL-2 can be promoted by molecular crowding, we next asked whether classical LLPS-promoting conditions influence chromatin condensation. We therefore promoted LLPS of the HPL-2 wild-type and 3xhinge I158A mutant proteins using 10% (w/v) PEG8000 (see Figure 2). Fluorescently labelled H3K9me0 or H3K9me3 chromatin arrays were subsequently added to assess their enrichment into the formed condensates. While H3K9me3 chromatin arrays enriched efficiently into wild-type HPL-2 condensates across the entire concentration range tested (25–100 µM), H3K9me0 arrays enriched weakly and only at the highest HPL-2 concentrations (Figure S12D). In contrast, the HPL-2 3xhinge I158A mutant showed inefficient enrichment of either H3K9me3 or H3K9me0 chromatin arrays, particularly at lower protein concentrations (Figure S12E). These results demonstrate that inducing LLPS in the dimerization-deficient mutant is insufficient to restore efficient H3K9me3 chromatin partitioning into HPL-2 condensates, further pointing to a critical role for HPL-2 dimerization in the condensation of H3K9me3 chromatin. Taken together, our findings (refer to Figure S13 for a phase map summarising the behaviour of all mutants, chromatin combinations and integrating turbidity and imaging data) uncouple the roles of dimerization and LLPS, establishing dimerization as the principal driver of HPL-2–mediated chromatin condensation *in vitro*.

### Differential condensation by HPL-2 drives segregation of H3K9me3 and H3K9me0 chromatin

Higher-order chromatin organisation involves not only condensation of individual domains but also spatial segregation of eu- and heterochromatin, a hallmark of cellular 3D genome organization [64, 65]. Since HPL-2 specifically condenses H3K9me3-modified chromatin arrays *in vitro*, we hypothesised that this property could extend to promoting selective partitioning of H3K9me3 chromatin in mixed chromatin environments. To test this idea, we established a chromatin segregation assay in which fluorescently labelled H3K9me0 and H3K9me3 12-mer chromatin arrays were co-induced to phase separate using a previously defined combination of monovalent and divalent salts (Figure 4A) [32]. In the absence of HPL-2, this approach produced uniform, homogeneous condensates containing overlapping signals from both array types thereby establishing a baseline phase-separated state independent of HPL-2 (Figure 4B). Under the conditions applied, the recombinant chromatin arrays themselves seem to lack intrinsic features that are sufficient to promote segregation.

**Figure 4.**
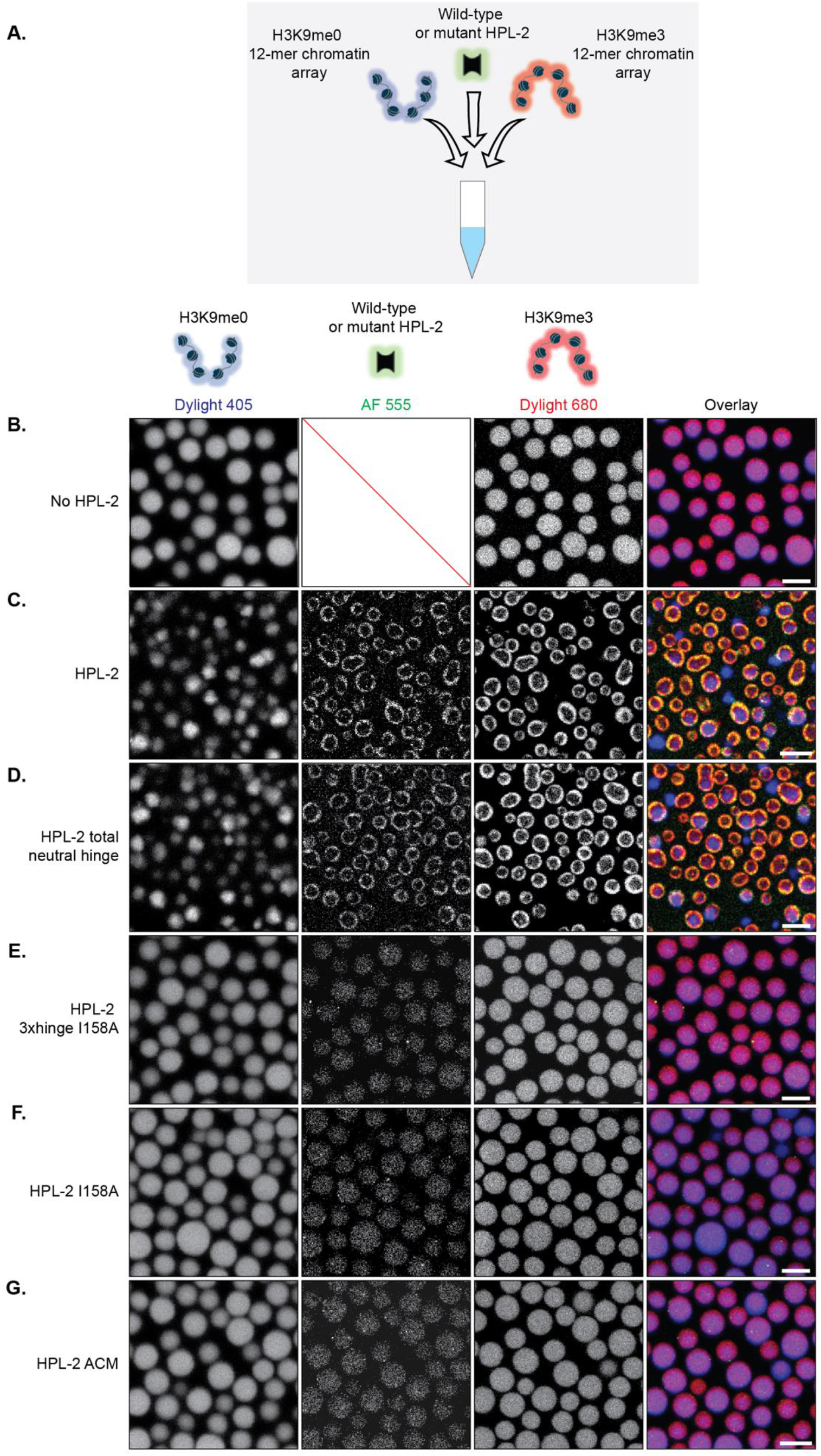
HPL-2 dimerization drives selective condensation and segregation of H3K9me3 chromatin arrays. **(A)** Scheme depicting the chromatin segregation assay. **(B–G)** Fluorescence microscopy of Dylight 405–labelled H3K9me0 and Dylight 680–labelled H3K9me3 chromatin arrays in the absence (B) or presence of wild-type (C) or the indicated HPL-2 mutant proteins (D–G) trace-labelled with Alexa Fluor 555–HPL-2. Scale bars: 5 μm. Images are representative of three independent experiments.

Addition of HPL-2 transformed the homogeneous chromatin condensates into biphasic, compartmentalised structures, in which H3K9me0 chromatin arrays preferentially occupied the central core, whereas H3K9me3 chromatin arrays and HPL-2 (AF555) co-partitioned into a condensed shell at the periphery (Figure 4C). Swapping fluorophore labels between the two chromatin types did not alter the segregation pattern (Figure S14A), confirming that this effect was modification-dependent rather than an artefact of fluorophore labelling. Quantification showed that HPL-2 partitioned more than twofold higher into the H3K9me3-rich compartment compared to the H3K9me0-rich compartment (Figure S14B), demonstrating preferential association with modified chromatin. Together with the condensation assays, these findings establish that HPL-2 not only condenses H3K9me3 chromatin more efficiently than unmodified chromatin but also drives its spatial segregation. We term this phenomenon “differential condensation-induced segregation” (DCIS).

To dissect the mechanistic basis of DCIS, we performed segregation assays using our HPL-2 mutant proteins. HPL-2 total neutral hinge mutant (dimerization-competent but LLPS-deficient) retained the ability to drive DCIS. The mutant protein produced biphasic condensates in which H3K9me3 arrays segregated from H3K9me0 arrays (Figure 4D), although the degree of segregation was moderately reduced compared to wild-type HPL-2, with ∼1.5-fold preferential partitioning into the H3K9me3 compartment (Figure S14B). This reduced enrichment correlates with the slightly decreased condensation efficiency of this mutant (Figure 3C; Figure S12B,C). By contrast, HPL-2 3xhinge I158A (LLPS-competent but dimerization-deficient), HPL-2 I158A (defective in LLPS and dimerization), and HPL-2 ACM (deficient in H3K9me binding) were completely incapable to induce DCIS (Figure 4E–G). In the presence of these mutant proteins, both H3K9me0 and H3K9me3 arrays remained uniformly mixed within condensates, with partitioning ratios close to 1 (Figure S14B). Collectively, these results reveal that HPL-2 dimerization is required to drive efficient condensation and spatial segregation of H3K9me3 chromatin arrays *in vitro*. It is tempting to speculate that the HPL-2 induced spatial segregation of chromatin through a differential condensation mechanism *in vitro* might be involved in the establishment and maintenance of cellular heterochromatin domains [66].

### HPL-2 mutant *C. elegans* show physiological and vulval defects

The 3D organization of eukaryotic genomes is essential for developmental homeostasis, and its disruption often leads to physiological abnormalities [67]. Having defined the molecular determinants enabling HPL-2–mediated chromatin condensation and segregation *in vitro*, we next examined their functional relevance *in vivo*. Previous studies, including ours, have shown that *C. elegans* HPL-2 and the malignant brain tumor (MBT)-domain protein LIN-61 act in parallel heterochromatin pathways [26], displaying extensive overlap with H3K9me2-enriched genomic regions [68]. Although single mutants exhibit relatively mild developmental phenotypes, combined loss of HPL-2 and LIN-61 results in synthetic germline and vulval defects [26]. We therefore analysed HPL-2 separation-of-function mutants in a *lin-61Δ* sensitized genetic background, enabling defects in HPL-2 function to be readily detected *in vivo*.

We generated *C. elegans* strains carrying *gfp*-tagged *hpl-2* alleles at the endogenous locus (i.e. under native control), each selectively perturbing a defined molecular feature: (i) *hpl-2::gfp* (control); (ii) *hpl-2 total neutral hinge::gfp* (dimerization-competent but LLPS-deficient); (iii) *hpl-2 3xhinge I158A::gfp* (LLPS-competent but dimerization-deficient); and (iv) *hpl-2 I158A::gfp* (LLPS- and dimerization-deficient). Of note, *hpl-2* has two splicing variants, *hpl-2a* and *hpl-2b* [22]. The insertion of a sequence encoding a GFP-tag at the C-terminus of *hpl-2* eliminated expression of the minor *hpl-2b* splicing variant, enabling exclusive analysis of the predominant HPL-2A protein (Figure S15A,B) [23]. For simplicity, we refer to the corresponding *C. elegans* strain and corresponding protein from here on as *hpl-2::gfp* and HPL-2::GFP. Western blotting of the different mutant embryos confirmed comparable expression of the different HPL-2::GFP proteins (Figure S15C,D).

Following established schemes [40, 41], we measured brood size, embryonic viability, sterility, and lifespan of the different *hpl-2* mutant *C. elegans* strains (*lin-61Δ* background) at 20°C (Figure S16, Table S7) and 25°C (Figure 5A–C, Table S7). In addition, we examined the high-temperature arrest (HTA) phenotype, a previously described consequence of HPL-2 loss at elevated temperature [44], by scoring larval arrest at 24°C and 26°C (Table S8). The single deletion mutant strains *hpl-2Δ* and *lin-61Δ* displayed high embryonic viability (∼95%) and lifespan comparable to the wild-type (N2) *C. elegans* strain at both temperatures. As reported before [26], mild sterility and modest brood size reduction was seen in *hpl-2Δ* mutants at 25°C, while showing robust HTA at 26°C. In contrast the *lin-61Δ; hpl-2Δ* double deletion mutant exhibited severe reproductive and lifespan defects at both 20°C and 25°C [26], establishing a sensitized reference for mutant analyses. The lifespan and brood size of both the *lin-61Δ*; *hpl-2::gfp* control strain and the *lin-61Δ*; *hpl-2 total neutral hinge::gfp* strain closely resembled those of the *lin-61Δ* single mutant strain at both 20°C and 25°C, indicating that the neutralization of charged residues in the hinge region only minimally affects HPL-2 function when dimerization remains intact. Consistent with these observations, both strains displayed only low frequencies of larval arrest at 26°C, comparable to the *lin-61Δ* mutant. By contrast, *lin-61Δ*; *hpl-2 3xhinge I158A::gfp* and *lin-61Δ*; *hpl-2 I158A::gfp C. elegans* showed pronounced phenotypes at elevated temperatures (25°C), including 20–30% sterility, markedly reduced brood size, and shorter lifespan compared to the control strain. These dimerization-defective mutants also exhibited modestly increased larval arrest at 26°C relative to the control and neutral hinge strains.

**Figure 5.**
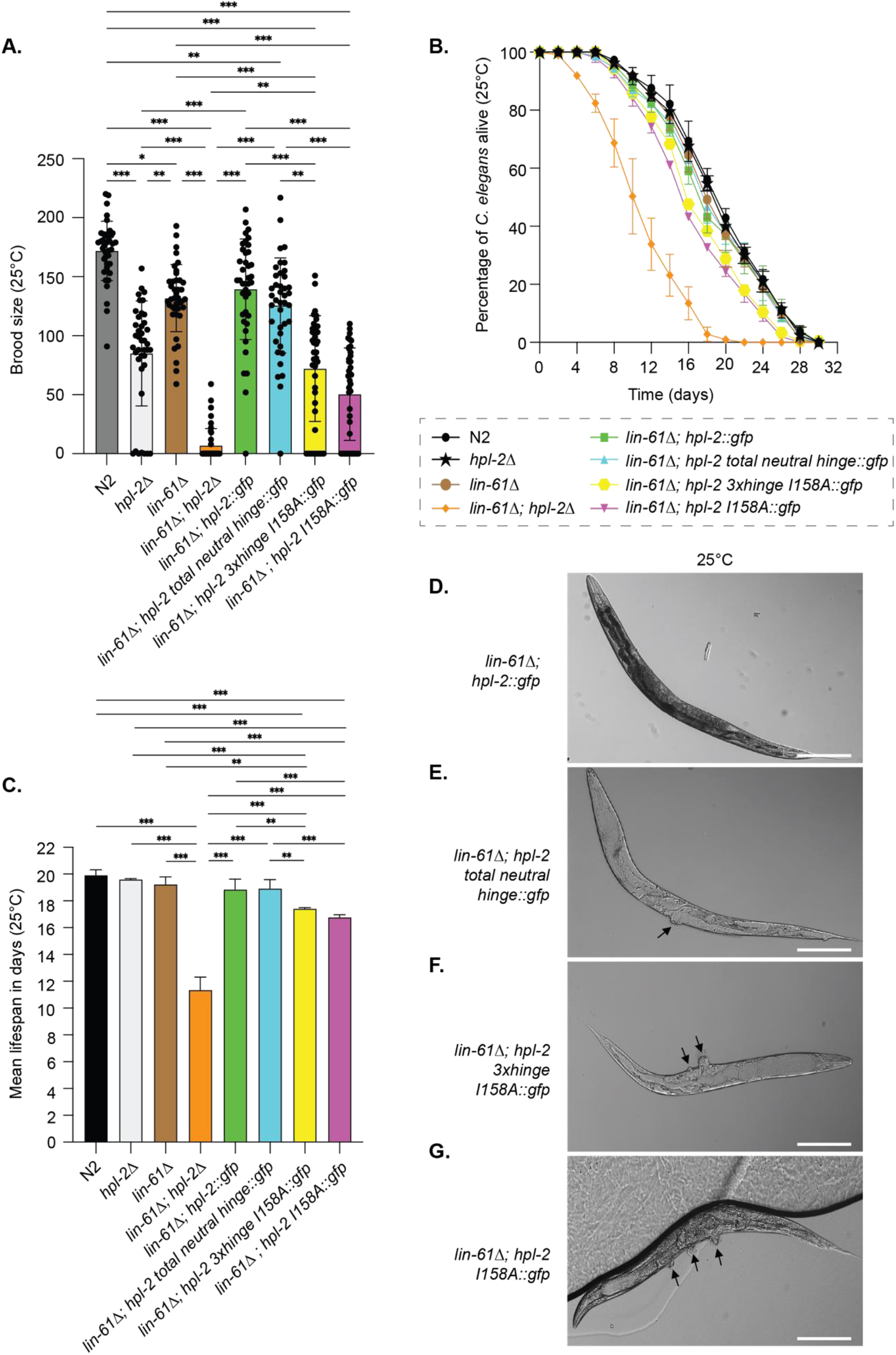
Effects of HPL-2 mutations on *C. elegans* fertility, lifespan, and vulval development at 25°C. **(A)** Mean brood size of non-sterile *C. elegans* expressing the indicated, HPL-2::GFP wild-type or mutant proteins in the background of *lin-61Δ* at 25°C. Data are representative of 40 worms analysed in four independent experiments. Error bars indicate standard deviation. *P<0.05, **P < 0.01, ***P < 0.001 by Kruskal–Wallis test with post hoc Dunn’s test. **(B)** Lifespan curves of *C. elegans* expressing HPL-2::GFP wild-type or mutant proteins in the background of *lin-61Δ* at 25°C. The fraction of surviving worms (in percent) is plotted over time; Day 0 corresponds to the L4-to-adult molt. Data represent 225 worms from three independent experiments. Colour codes for genotypes are shown below the curves. **(C)** Mean lifespan (in days) of *C. elegans* expressing HPL-2::GFP wild-type or mutant proteins in the background of *lin-61Δ* at 25°C. Survival functions were estimated using the Kaplan–Meier method, and mean survival times were calculated as the area under the survival curve. Error bars denote standard error of the mean (SEM). *P < 0.05, **P < 0.01, ***P < 0.001 by Mantel–Cox log-rank test. **(D– G)** Representative differential interference contrast (DIC) microscopy images of *C. elegans* of the indicated genotypes at 25°C. Arrows denote vulval protrusions indicative of a synMuv phenotype. Scale bars: 100 μm.

We then analysed vulval development, a sensitive readout of chromatin-dependent gene regulation in *C. elegans* [69]. All mutant strains showed normal vulva development at 20°C (Figure S16D–G, Table S9). However, at the elevated temperature of 25°C, abnormal vulval development was observed in specific *hpl-2* mutant strains. Consistent with our previous findings [26], *lin-61Δ; hpl-2Δ* double mutants developed ectopic pseudovulval protrusions (>50% of animals) at 25°C (Table S9), indicative of a synthetic multivulva (synMuv) phenotype arising from a failure to restrict vulval cell fates [70]. In contrast, single *lin-61Δ* or *hpl-2Δ* mutant and control *lin-61Δ*; *hpl-2::gfp C. elegans* had normal vulva development (Figure 5D, Table S9). Strikingly, *lin-61Δ*; *hpl-2 3xhinge I158A::gfp* and *lin-61Δ*; *hpl-2 I158A::gfp C. elegans* exhibited severe synMuv phenotypes (∼40% of animals), similar to the *lin-61Δ; hpl-2Δ* double mutant at 25°C, whereas *lin-61Δ*; *hpl-2 total neutral hinge::gfp C. elegans* showed only mild defects (∼9%) (Figure 5E–G, Table S9). Collectively, these findings suggest that dimerization is an essential determinant of HPL-2 physiological function *in vivo*.

### HPL-2 dimerization, but not hinge-mediated multivalency, is required for HPL-2 nuclear foci formation in *C. elegans* embryos

Condensed heterochromatin in animal and plant cells often forms microscopically distinct foci, such as chromocenters in *Mus*, *Drosophila*, and *Arabidopsis* nuclei [71, 72]. Similarly, *C. elegans* embryos exhibit discrete heterochromatic foci that provide a tractable mesoscale readout of chromatin condensation *in vivo* [73]. To dissect the contributions of HPL-2 dimerization–mediated bridging and hinge region-mediated LLPS to heterochromatin organization, we first analysed the formation of HPL-2 foci in nuclei of *C. elegans* embryos (100–200 cells stage) expressing wild-type or mutant HPL-2 proteins in the sensitised *lin-61Δ* background using live imaging.

Genome-wide profiling has shown that HPL-2 is enriched on autosomal arms corresponding to large-scale heterochromatic domains [21, 23, 68, 74]. Live imaging of *lin-61Δ*; *hpl-2::gfp* embryos revealed distinct nuclear foci of GFP signal enrichment (Figure 6A), with a median of ∼5 per nucleus occupying ∼5% of the total nuclear area (Figure 6E,F). These foci were retained in the *lin-61Δ*; *hpl-2 total neutral hinge::gfp* strain, with a modest but non-significant reduction in their number (∼3 per nucleus) and area (∼3%) (Figure 6B,E,F). By contrast, nuclei of embryos expressing dimerization-deficient HPL-2 mutant proteins (*lin-61Δ; hpl-2 I158A::gfp* and *lin-61Δ; hpl-2 3xhinge I158A::gfp*) did not show discernible foci (Figure 6C–F), demonstrating that HPL-2 dimerization is essential for its foci formation. Hinge-mediated multivalency might serve to enhance or stabilize these structures.

**Figure 6.**
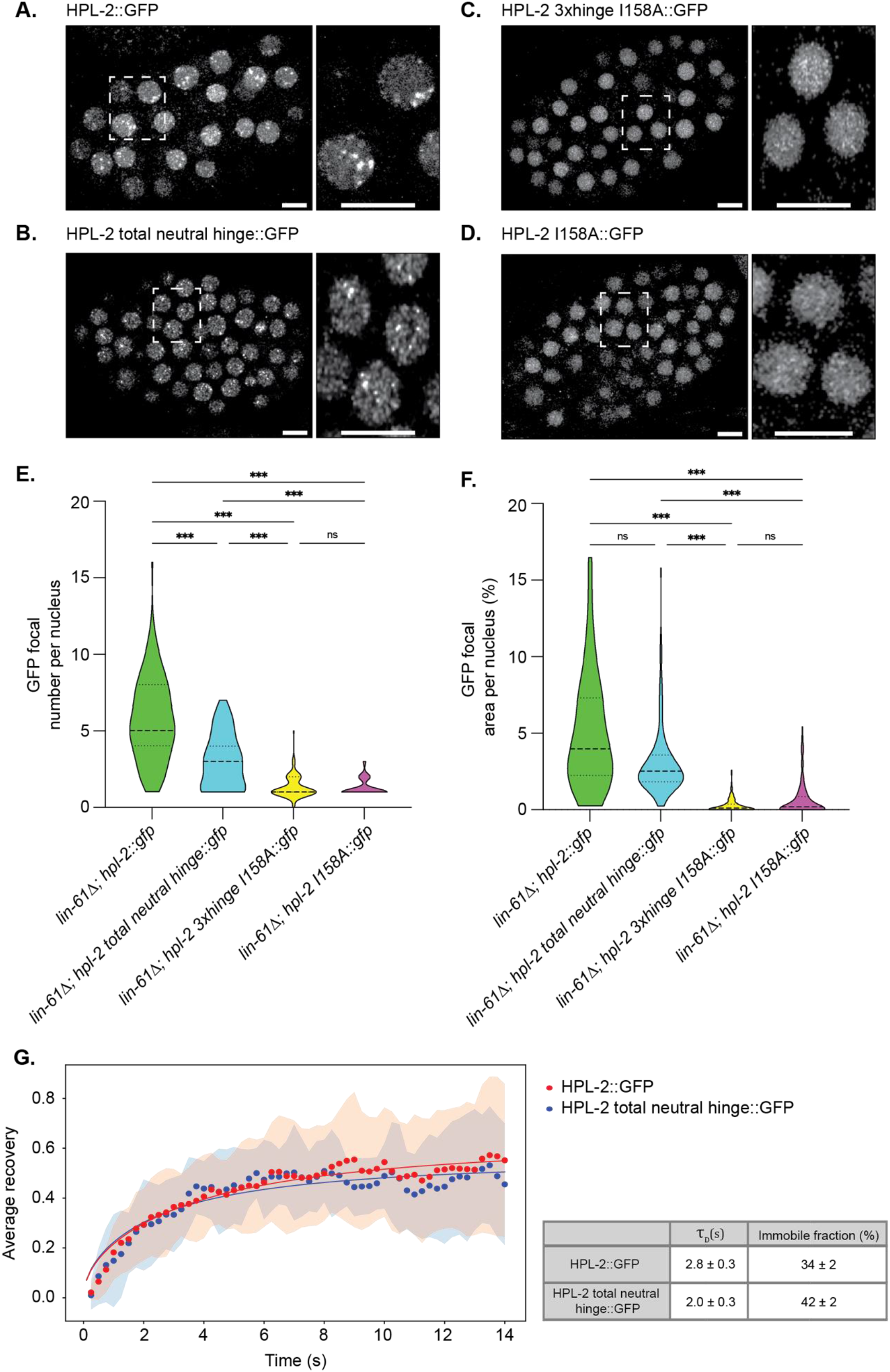
HPL-2::GFP foci formation is impaired in *C. elegans* embryos expressing HPL-2 dimerization-deficient mutant proteins. **(A–D)** Maximum intensity z-projected fluorescence microscopy images of live *C. elegans* embryos expressing HPL-2::GFP wild-type (A) or mutant proteins (B–D) in the background of *lin-61Δ*. Insets show magnified views of nuclei indicated by dashed boxes. Scale bars: 5 μm. Images are representative of three independent experiments. **(E, F)** Quantification of HPL-2::GFP foci in the nucleus of embryos of the indicated *C. elegans* strains from panel (A–D), expressed as the number of foci per nucleus (E) and as the percentage of focal area per nucleus (F). Data are shown as violin plots; dotted lines indicate the 25th and 75th percentiles, and the dashed line marks the median. N = 115 nuclei per genotype from three independent experiments. *P < 0.05, ***P < 0.001 by Kruskal–Wallis test with post hoc Dunn’s test. (G) Fluorescence recovery after photobleaching (FRAP) of nuclear foci formed by HPL-2::GFP and HPL-2 total neutral hinge::GFP in live *C. elegans* embryos in the background of *lin-61Δ*. Data points represent the mean of 15 (HPL-2::GFP) and 13 (HPL-2 total neutral hinge::GFP) measurements. Error bars show the standard deviation. The fit parameters for both curves are listed in the table.

To assess whether hinge-mediated multivalency influences the dynamics of HPL-2 foci *in vivo*, we performed fluorescence recovery after photobleaching (FRAP) on HPL-2::GFP nuclear foci in *lin-61Δ; hpl-2::gfp* and *lin-61Δ; hpl-2 total neutral hinge::gfp* embryos (Figure 6G). HPL-2::GFP foci in both strains recovered rapidly after photobleaching and exhibited highly similar recovery kinetics. Fitting the recovery curves yielded comparable characteristic recovery times *τ* = 2.8 s for HPL-2::GFP and *τ* = 2.0 s for HPL-2 total neutral hinge::GFP foci and similar immobile fractions (34% and 42%, respectively). Thus, despite forming slightly fewer and smaller nuclear foci, the HPL-2 total neutral hinge mutant retains exchange dynamics comparable to those of wild-type HPL-2.

To determine whether the loss of HPL-2 nuclear foci resulted from impaired chromatin recruitment rather than defective condensation, we quantified chromatin occupancy of GFP-tagged HPL-2 proteins by ChIP-qPCR. Fifteen heterochromatic target loci and five negative-control loci were selected based on HPL-2 and H3K9me2 ChIP-seq datasets from mixed-stage embryos, with target loci corresponding to regions of overlapping HPL-2 and H3K9me2 enrichment and off-target loci corresponding to chromosomal centre regions lacking enrichment for either factor (Table S6). As expected, GFP ChIP of HPL-2::GFP from *lin-61Δ; hpl-2::gfp* embryos showed robust enrichment at heterochromatic target loci (median 0.56% input) but negligible signal at off-target loci, whereas GFP ChIP from N2 embryos (which lack a GFP-tagged protein) and IgG controls showed only background enrichment (Figure S17A,B). The HPL-2 total neutral hinge::GFP protein exhibited comparable chromatin occupancy (median 0.54% input) at the target loci (Figure S17A,B). The GFP tagged dimerization-deficient mutants showed only a modest reduction in occupancy compared to HPL-2::GFP, with median enrichments of 0.44% input for HPL-2 I158A::GFP (∼21% reduction) and 0.40% input for HPL-2 3xhinge I158A::GFP (∼29% reduction), while remaining strongly enriched over background (Figure S17A,B). We note that despite retaining approximately 70–80% of wild-type chromatin occupancy, both dimerization-deficient mutants display an almost complete loss of HPL-2 nuclear foci (>90% reduction in foci area; Figure 6F). Apparently, the loss of HPL-2 foci is not caused by impaired chromatin recruitment but likely reflects defects in HPL-2-mediated chromatin condensation.

### HPL-2 dimerization is required for organization of H3K9me2-marked heterochromatin foci independently of H3K9me2 deposition

To assess whether mutation of HPL-2 dimerization and/or LLPS affect heterochromatin organization, we examined nuclear H3K9me2 foci, which mark canonical heterochromatic domains in *C. elegans* embryos [75]. Because HPL-2 predominantly co-distributes with chromatin areas containing H3K9me2 rather than H3K9me3 [21], and because loss of LIN-61 reduces H3K9me3 levels via impaired recruitment of the H3K9 methyltransferase SET-25 [76], H3K9me2 foci provide an informative readout in this context.

Western blotting of *C. elegans* embryos confirmed that global H3K9me2 levels were unchanged in the strains expressing different mutant HPL-2 proteins in *lin-61Δ* background (Figure S18A,B). To determine whether the genomic distribution of H3K9me2 was altered, we additionally performed H3K9me2 ChIP-seq. Genome-wide H3K9me2 profiles across consensus peaks were highly similar between wild-type N2 and GFP-tagged HPL-2 mutant strains in *lin-61Δ* background with only a minor reduction in H3K9me2 enrichment observed in the *lin-61Δ; hpl-2 I158A::gfp* mutant (Figure 7A and Figure S18C). In contrast, H3K9me2 signal was nearly absent in the *set-32; met-2 set-25* (triple deletion of H3K9 methyltransferases) [77] mutant and was not detected in the IgG control.

**Figure 7.**
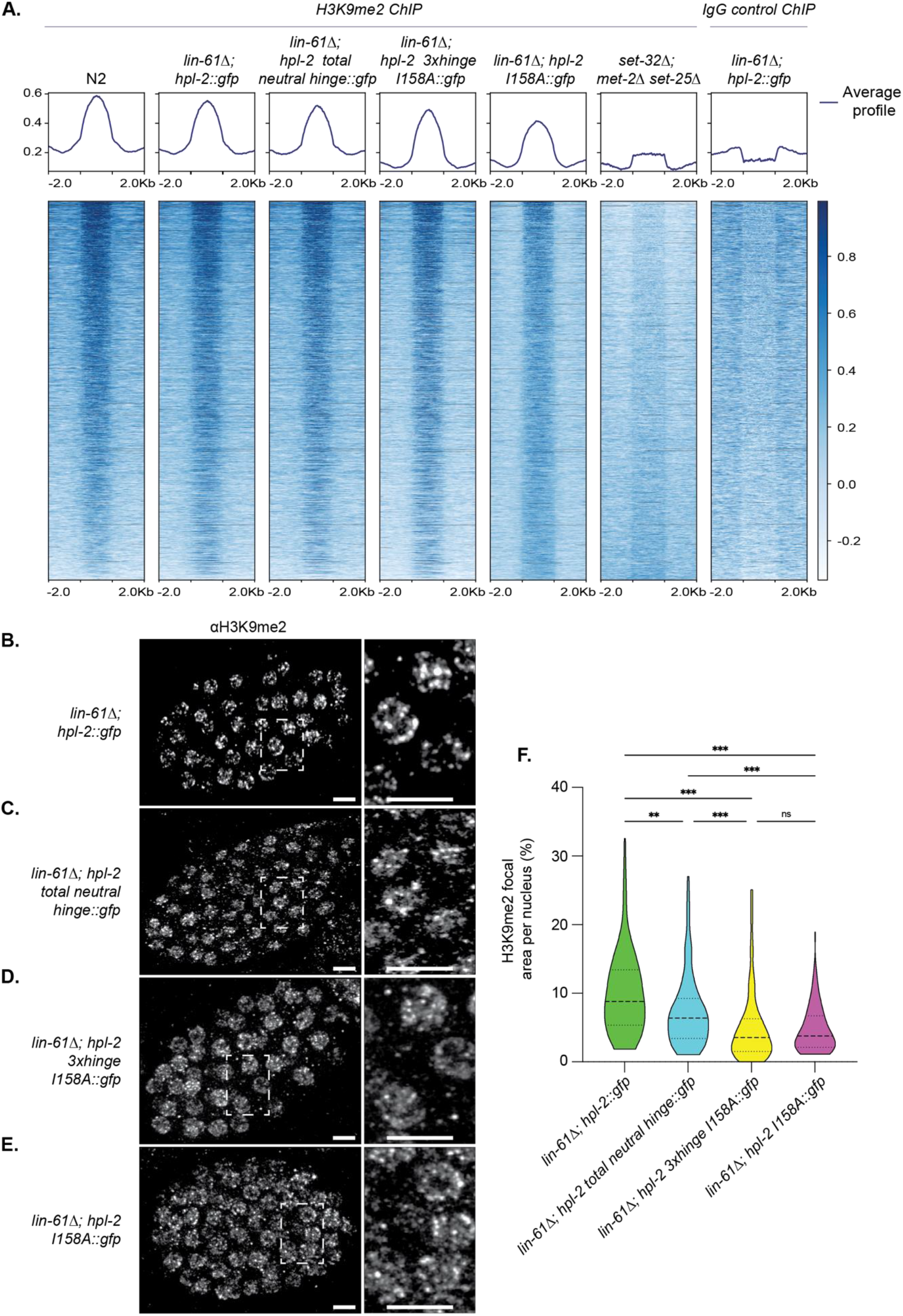
H3K9me2 nuclear foci are impaired in *C. elegans* embryos expressing HPL-2 dimerization-deficient mutant proteins. **(A)** Heat maps showing input-normalized H3K9me2 ChIP-seq signal (log_2_ fold change over input) across consensus H3K9me2 peaks in different wild-type and mutant *C. elegans* strains. Consensus peaks were defined as regions present in at least three biological replicates and are ordered according to H3K9me2 signal intensity in *lin-61Δ; hpl-2::gfp* embryos. Heat maps depict the average log_2_ fold change from three biological replicates over a 6-kb window centred on a 2kb representation of each broad peak with adjoining regions (−2 kb to +2 kb). Average enrichment profiles across all consensus peaks are shown above the heat maps. The *set-32; met-2 set-25* mutant that is deficient in H3K9 KMTs [77] and IgG ChIP served as negative controls. **(B–E)** Maximum-intensity z-projected immunofluorescence images of *C. elegans* embryos stained with antibodies against H3K9me2 expressing the indicated HPL-2 mutant proteins in the background of *lin-61Δ*. Insets show magnified views of nuclei indicated by dashed boxes. Scale bars: 5 μm. Images are representative of three independent experiments. **(F)** Quantification of H3K9me2 foci from panels (B–E), expressed as the percentage of focal area per nucleus. Data are shown as violin plots; dotted lines indicate the 25th and 75th percentiles, and the dashed line marks the median. N = 115 nuclei per genotype from three independent experiments. *P < 0.05, ***P < 0.001 by Kruskal–Wallis test with post hoc Dunn’s test.

Immunofluorescence imaging of *C. elegans* embryos revealed prominent H3K9me2 foci in the *lin-61Δ*; *hpl-2::gfp* control strain, occupying ∼10% of the total nuclear area (Figure 7A,E). The *lin-61Δ*; *hpl-2 total neutral hinge::gfp* strain displayed slightly reduced foci (∼8% nuclear area; Figure 7B,E), indicating that hinge-mediated multivalency might enhance but is not essential for heterochromatin foci formation. In contrast, both mutant strains expressing dimerization-deficient HPL-2 proteins (*lin-61Δ; hpl-2 I158A::gfp* and *lin-61Δ; hpl-2 3xhinge I158A::gfp*) showed a severe phenotype, with H3K9me2 foci reduced to ∼5% of the total nuclear area, half that of the control strain (Figure 7C–E). Together, these data suggest that HPL-2 dimerization is critical for organizing H3K9me2-marked heterochromatin *in vivo*.

### Loss of HPL-2-mediated chromatin condensation has only modest effects on repetitive element and gene expression in *C. elegans* embryos

Heterochromatin is commonly associated with transcriptional silencing, with its condensed state thought to restrict access of RNA polymerase II and associated factors to DNA [78–80]. To test whether impaired HPL-2–mediated chromatin condensation alters transcription, we performed total RNA-seq on *C. elegans* embryos expressing wild-type or mutant HPL-2 proteins in the *lin-61Δ* background. Principal component analysis of all RNA-seq samples, showed clustering of biological replicates and also genotypes (Figure S19A), indicating that the technical and/ or developmental variation within replicates was limited.

Because H3K9me-marked heterochromatin is critical for silencing repetitive elements [2], we first analysed the RNA-seq data using a repeat-aware quantification pipeline [52]. Across all major classes of repetitive elements, including Class I (RNA) transposons, Class II (DNA) transposons, inverted repeats (IRs), and satellite repeats, expression changes remained modest, with only 24 of the 177 quantified repeat families showing significant upregulation in any of the mutant strains (Figure 8A). The *lin-61Δ; hpl-2 total neutral hinge::gfp* strain exhibited little or no repetitive element derepression, whereas mild increases were observed in the dimerization-deficient mutants. These changes were primarily restricted to a subset of DNA transposons, including members of the Mariner and Merlin families, and a small number of LTR retrotransposons within the Gypsy and Bel-Pao superfamilies.

**Figure 8.**
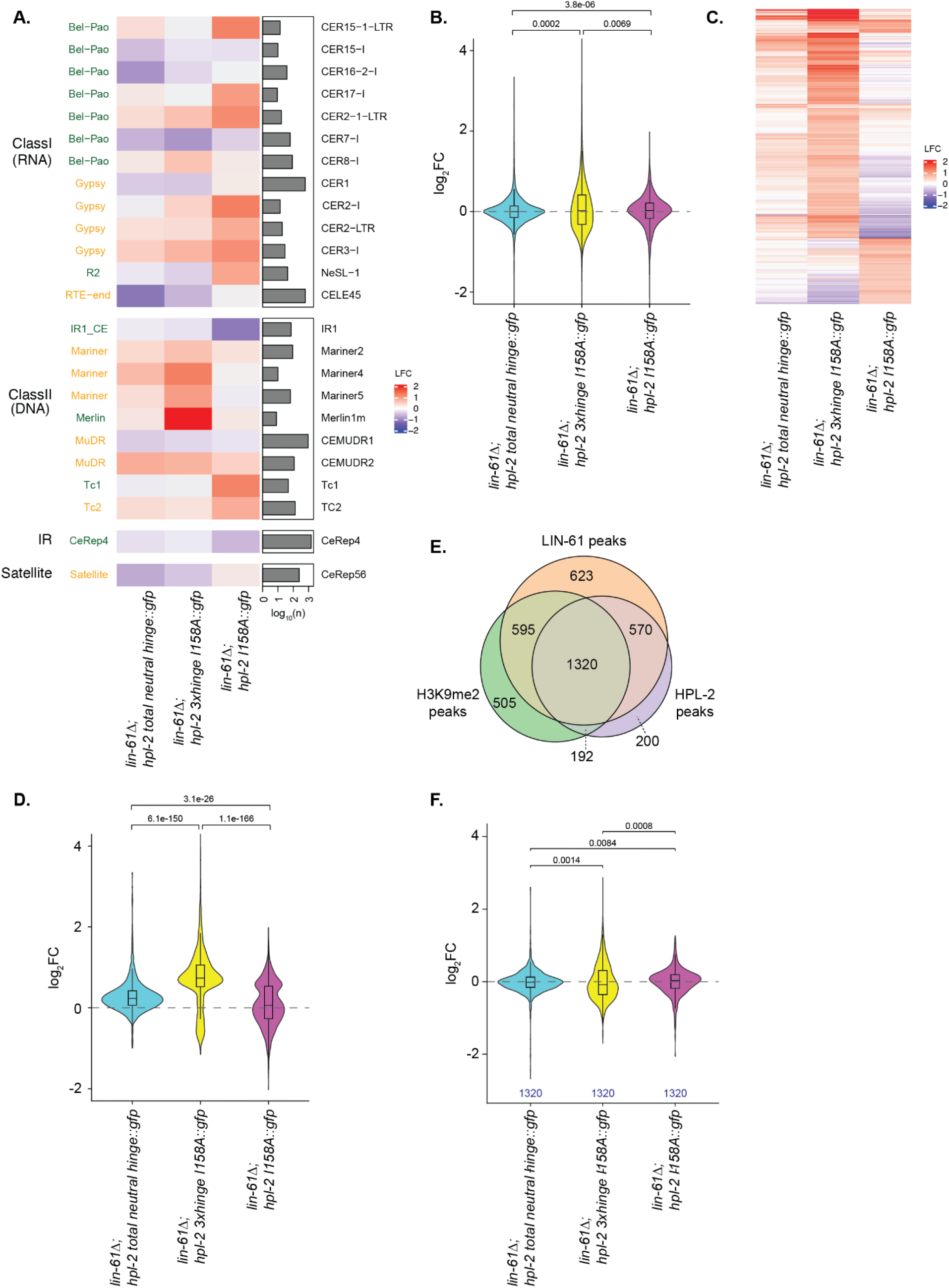
Transcriptional status is largely maintained at canonical heterochromatin in *C. elegans* embryos expressing HPL-2 dimerization-deficient mutant proteins. **(A)** Heatmap showing the log₂ fold change (log₂FC) of repetitive element families in embryos of the indicated mutant *C. elegans* strains relative to *lin-61Δ; hpl-2::gfp*. Repetitive elements are grouped into Class I (RNA transposons), Class II (DNA transposons), inverted repeats (IR), and satellite repeats. The number of genomic copies for each repetitive element family is indicated on the right. **(B)** Violin plot quantification of log₂FC for all expressed genes (those not excluded by DESeq2 automatic filtering) located on the autosomal arms in embryos of the indicated mutant *C. elegans* strains, using *lin-61Δ; hpl-2::gfp* as the control. The violin plots show the distribution of log₂FC values, with the embedded boxplot indicating the interquartile range and median. Whiskers represent the data range. **(C-D)** Genes significantly upregulated in at least one sample (padj < 0.05; log₂FC > 0.5), relative to *lin-61Δ; hpl-2::gfp*, are shown as a heatmap with rows (genes) ordered by hierarchical clustering (C) and quantified as violin plots with embedded boxplots showing the interquartile range and median (D). **(E)** Venn diagram of genes on the autosomal arms with H3K9me2/HPL-2/LIN-61 ChIP signals in the gene body. The number of genes in each category overlapping at least 1 ChIP peak of a particular dataset are specified. **(F)** Violin plot quantification of RNAseq log₂FC in embryos of mutant *C. elegans* strains using *lin-61Δ; hpl-2::gfp* as the control. 1,320 genes located on the arms of autosomes bearing overlapping H3K9me2/HPL-2/LIN-61 ChIP peaks in the gene body were considered. The FDR-adjusted p values shown in (B, D, F) were obtained by a two-sided Wilcoxon rank sum test of all possible two-way comparisons of samples. Only significant (padj<0.05) comparisons are shown.

We then focused on gene expression from the predominantly heterochromatic arms of the autosomes [20]. Of the 23,018 annotated genes located on autosomal arms (7,687 protein-coding and 15,331 non-coding), most showed very low expression and were therefore excluded from further analysis. This filtering retained 6,747 genes (6,069 protein-coding and 678 non-coding) as expressed. RNA-seq analysis of N2, *hpl-2Δ*, *lin-61Δ*, and *lin-61Δ*; *hpl-2Δ* embryos showed that loss of *hpl-2* alone resulted in limited transcriptional changes, whereas combined loss of *hpl-2* and *lin-61* caused broader transcriptional alterations relative to N2, supporting the use of the *lin-61Δ* background as a sensitized genetic context to investigate the transcriptional consequences of HPL-2 mutant proteins (Figure S19B). Differential expression relative to the control *lin-61Δ; hpl-2::gfp* strain revealed both up- and downregulated genes across the *C. elegans* strains expressing different mutant HPL-2 proteins. A slight overall bias toward upregulation was seen in the strains expressing HPL-2 dimerization-deficient mutants (*lin-61Δ; hpl-2 I158A::gfp* and *lin-61Δ; hpl-2 3xhinge I158A::gfp*) (Figure 8B).

Of the 6,747 expressed genes, 1,675 (1,561 protein-coding and 114 non-coding) were significantly upregulated in at least one mutant strain. Hierarchically clustered expression profiles revealed broadly similar patterns of transcriptional change between the *lin-61Δ; hpl-2 total neutral hinge::gfp* and *lin-61Δ; hpl-2 3xhinge I158A::gfp* strains. Although the magnitude of upregulation was markedly greater in the latter (Figure 8C, D), like the increased transcriptional upregulation observed in the *lin-61Δ; hpl-2Δ* double mutant (Figure S19C). By contrast, the *lin-61Δ; hpl-2 I158A::gfp* strain displayed both up- and downregulation at distinct subsets of genes, suggesting a dominant-negative effect on transcriptional control. Most of the genes upregulated in the *lin-61Δ; hpl-2 3xhinge I158A::gfp* strain pertained to transmembrane transport, stress response and neuronal function (Figure S19D).

To further explore the genomic context of gene derepression, we focused specifically on genes located on autosomal arms that showed H3K9me2 enrichment and were co-occupied by HPL-2 and LIN-61. This analysis identified 1,320 such loci (Figure 8E). Strikingly, transcriptional fold-changes for this subset remained largely stable across all HPL-2 mutants, even in *C. elegans* strains expressing HPL-2 dimerization-deficient mutants where we had found H3K9me2 foci area reduced by more than two-fold (Figure 8F). Together, our analyses indicate that loss of HPL-2-mediated chromatin condensation has only modest effects on both gene expression and repetitive element silencing. Heterochromatin condensation and transcriptional repression appear therefore partially uncoupled in *C. elegans* embryos.

## DISCUSSION

### Bridging via HPL-2 dimerization condenses H3K9me chromatin

Our combined *in vitro* and *in vivo* findings establish a mechanistic framework for HPL-2, suggesting that bridging is at the core of its function in heterochromatin condensation (Figure 9). While structural studies have verified that mammalian HP1 can bridge adjacent dinucleosomes, we infer from our *in vitro* condensation system that such bridging must extend to more distal nucleosomes (intra-fibre) and especially on different chromatin fibres (inter-fibre). With the short 12-mer chromatin arrays, condensate formation seems unlikely when only considering intra-strand bridging interactions. The relative low binding affinities (micromolar range) and rebinding kinetics reported for HP1/nucleosome complexes [13, 81] further suggest that the resulting interaction network (mesh) is highly dynamic, with bridging interactions constantly forming and reorganizing, resulting in a plastic state of condensed chromatin (Figure 9A).

**Figure 9.**
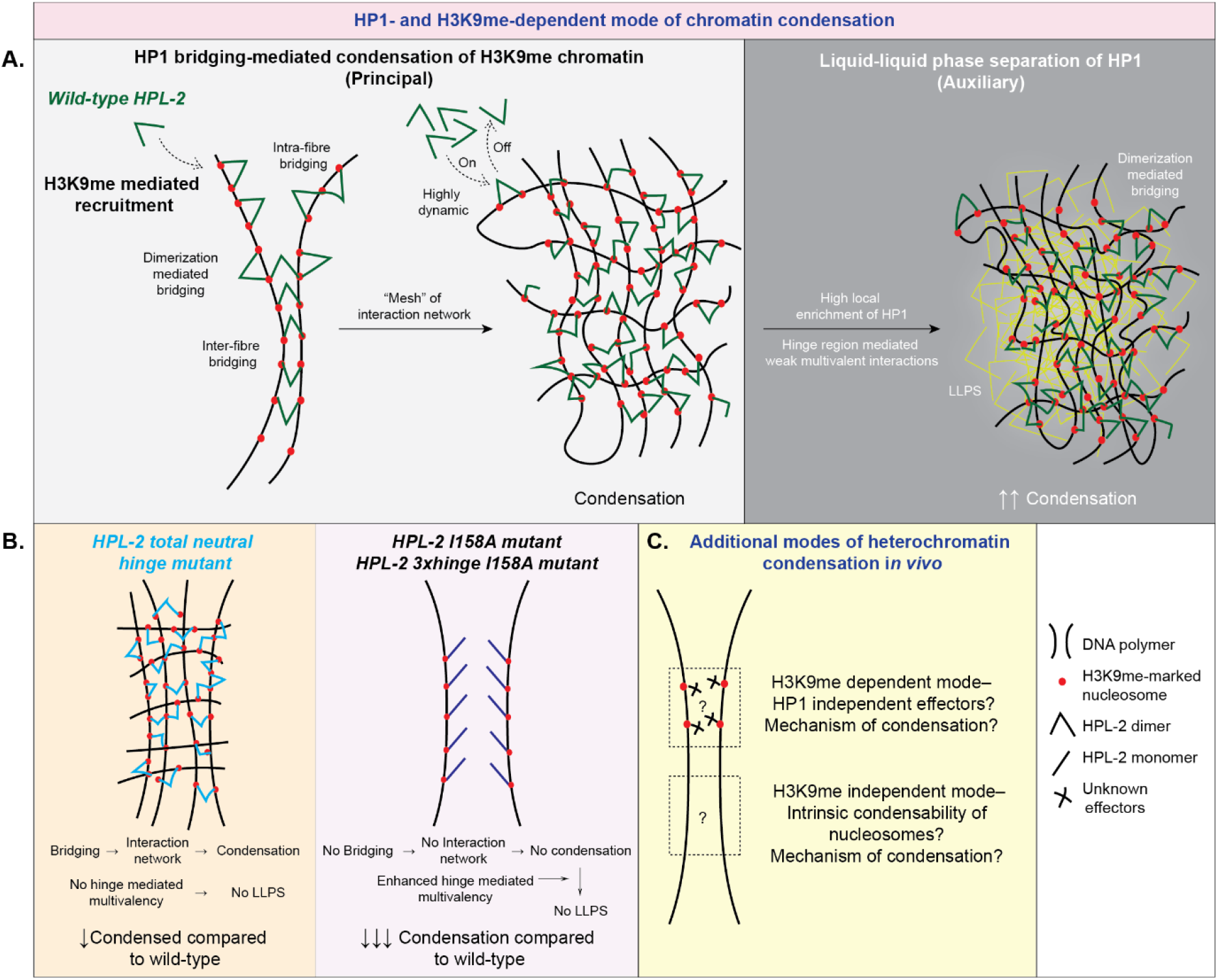
Mechanism of HPL-2-mediated H3K9me-marked heterochromatin condensation. **(A)** HPL-2 is recruited to H3K9me via CD–H3K9me interactions. Dimerization of HPL-2 induces intra- and inter-fibre bridging between H3K9me nucleosomes, forming a network (“mesh”) of crosslinked chromatin. This constitutes the principal mechanism of HP1–mediated heterochromatin condensation (left). The transient nature of CD–H3K9me keeps the condensed chromatin in a dynamic, viscoelastic state. The high local enrichment enables hinge region mediated LLPS of HPL-2, resulting in augmented and modulated heterochromatin condensation (right). **(B)** Dimerization enables bridging-mediated condensation, but lack of multivalent interactions mediated by the hinge region prevents LLPS of the HPL-2 neutral hinge mutant protein, resulting in reduced H3K9me chromatin condensation (left). Loss of dimerization in the HPL-2 I158A mutant protein prevents bridging-mediated condensation of H3K9me chromatin. In the absence of high local protein concentration, LLPS of HPL-2 I158A is not possible, even when multivalency mediated by the hinge region is enhanced in the HPL-2 3xhinge I158A mutant protein (right). **(C)** Heterochromatin is condensed via multiple, partially redundant pathways, including H3K9me- and HP1-independent pathways.

While at the mesoscale, condensed chromatin has been reported to exhibit “solid-like” behaviour [82], fibre entanglement is expected to prevent true liquid-like behaviour above the scale of nucleosome clutches (a zig-zag stretch of 1–2 kb of the 10 nm fibre) [83, 84]. Consistent with this, our FRAP measurements of HPL-2::GFP nuclear foci show rapid partial recovery (τ ≈ 2.8 s) alongside a substantial immobile fraction (∼34%; Figure 6G), a signature more consistent with a dynamic, viscoelastic condensate than with a freely mixing liquid. We therefore propose that HPL-2 dimerization maintains condensed chromatin above the sol-gel transition, in a viscoelastic state that remains more permissive to molecular exchange than a fully solid assembly, potentially enabling heterochromatin to respond dynamically to developmental cues, cell cycle progression, or stress [85, 86]. This view aligns with the emerging model that chromatin exhibits viscoelastic behaviour consistent with both liquid- and solid-like properties depending on the relevant time (seconds vs minutes–hours) and length (stretches of nucleosome vs entire chromosomes) scales [8, 63, 87].

Beyond HPL-2-mediated bridging, chromatin can also condense through intrinsic electrostatic interactions, including those mediated by the positively charged histone tails and, *in vivo*, linker histone H1, both of which have well-established roles in promoting contacts and crosslinking between chromatin fibres [63]. The relative contributions of these intrinsic interactions, HP1 dimerization-mediated bridging and LLPS are expected to be sensitive to the ionic and physicochemical environment. Our chromatin condensation assays were performed at relatively low ionic strength (50 mM NaCl), whereas HPL-2 LLPS was examined at 150 mM NaCl, with molecular crowding further promoted by PEG8000. Increasing ionic strength can screen charge-dependent interactions underlying both chromatin self-association and HPL-2 LLPS, while molecular crowding can independently favour LLPS. Hence, the competition between bridging and LLPS is likely to shift with buffer composition rather than being an invariant property of HPL-2. Nevertheless, even under LLPS-promoting conditions, the LLPS-competent but dimerization-deficient HPL-2 3xhinge I158A mutant protein remained impaired in H3K9me3 chromatin enrichment (Figure S12D,E), indicating that enhanced LLPS cannot substitute for dimerization-dependent bridging under the conditions examined. Thus, while ionic strength, molecular crowding and additional chromatin-associated factors may shift the balance between these mechanisms *in vivo*, our data identify dimerization-mediated bridging as the dominant HPL-2-dependent contribution across the distinct physicochemical conditions examined.

### Modulatory role of HPL-2 LLPS

The chromatin-condensation behavior of HP1 proteins is depending on the local protein concentration. In mouse fibroblasts, an average nuclear HP1α concentration of ∼1 μM has been reported [88], while fluorescence-based measurements of HP1α enrichment in chromocenters indicated local concentrations of up to ∼3 μM [89]. These concentrations are significantly lower than the concentrations required for HP1α LLPS *in vitro* [14], which approximate the level obtained when assuming the binding of one HP1α dimer per closely spaced nucleosome (100 μM). While the endogenous concentration of HPL-2 is not known, we propose that bridging-mediated chromatin condensation locally enriches HPL-2 to concentrations at which weak multivalent interactions mediated by the hinge region may become relevant for LLPS, thereby further enhancing or stabilising chromatin condensation (Figure 9A). The modest reduction in the focal area occupied by the HPL-2 total neutral hinge::GFP mutant protein in embryo nuclei relative to the wild-type is consistent with such an auxiliary contribution of LLPS *in vivo* (Figure 6B,F). We also note that mutation or repositioning of specific residues within the hinge region, including the RKR motif, markedly alters HPL-2 condensate formation *in vitro* (Figure S9C,E,F). This is consistent with growing evidence that sequence-encoded determinants, including short linear motifs (SLiMs) [90], emerging as critical regulators of biological condensate specificity [91]. We therefore interpret HPL-2 LLPS as an auxiliary mechanism that enhances or modulates dimerization-dependent, bridging-mediated chromatin condensation (Figure 9B).

### H3K9me in HP1 chromatin recruitment and condensation

In our *in vitro* chromatin condensation systems, recruitment is exclusively mediated by binding of the HPL-2 CD to H3K9me (Figure 3F). This interaction mode has been well characterized through biochemical, biophysical, and structural analyses of various HP1 orthologues [13, 55]. While genomic analyses in diverse cellular systems, including *C. elegans*, consistently reveal strong overlap between H3K9me2/3-marked heterochromatin and HP1 proteins [20], the exact mode of HP1 recruitment to heterochromatin *in vivo* is less clear. For example, the interaction of HPL-2 with the PxVxL motif–containing protein LIN-13, which co-localises with HPL-2 on the heterochromatic arms of *C. elegans* chromosomes, has been proposed to constitute a H3K9me-independent mode of recruitment [24, 92]. Additionally, the hinge region of HP1 proteins can associate non-specifically with nucleic acids, potentially stabilizing its chromatin binding [12]. Notably, work in *Drosophila* suggests that HP1 recruitment and higher-order chromatin organization are mechanistically separable: HP1 remains chromatin-associated under conditions of reduced H3K9 methylation, yet fails to form higher-order clusters, indicating that HP1 recruitment alone is insufficient to establish higher-order heterochromatin organization [93]. While multiple, overlapping pathways likely contribute to HPL-2 localization at heterochromatin *in vivo*, genome wide mapping studies and fluorescence microscopy of *C. elegans* embryos mutant in the H3K9 methylation system support a significant role of the modification in the recruitment of the protein to its target sites and subsequent condensation of chromatin [21].

Although our findings show that HPL-2 dimerization is crucial for heterochromatin condensation, H3K9me2 foci are not completely lost in embryos expressing dimerization-deficient proteins (Figure 7C,D). Persistence of heterochromatin condensation in systems impaired in HP1 function was also seen in HP1α knockout mice that maintain chromocenters [89] and in *Drosophila* tissues that show persistence of self-association among pericentromeric regions in the absence of HP1a [94]. Apparently, heterochromatin condensation relies on multiple, partially redundant pathways (Figure 9C). Such pathways may not only account for the residual H3K9me2 foci observed in the dimerization-deficient mutants but could also contribute indirectly to the changes observed upon disruption of HPL-2 dimerization. Thus, while our *in vitro* reconstitution experiments establish a direct requirement for HPL-2 dimerization in H3K9me3-dependent chromatin condensation, the corresponding *in vivo* phenotypes likely reflect the combined contributions of HPL-2 and other chromatin-organizing mechanisms. Some of these seem to rely on H3K9me, others might be independent thereof. This view is supported by the observation that the nuclear area occupied by HPL-2 foci (∼5%) is significantly less than that occupied by H3K9me2 foci (∼10%). Candidate systems that could work via H3K9me but independent of HPL-2 include the H3K9 methyltransferase MET-2 and its intrinsically disordered cofactor LIN-65 [92, 95]. The H3K9me-independent pathways of heterochromatin organization might be fundamental and not requiring accessory factors as it was shown recently that native nucleosomes from mammalian cells intrinsically encode features of genome organization [96].

### Differential condensation-induced segregation (DCIS) in heterochromatin domain formation

The mechanism of spatial segregation of euchromatin and heterochromatin within the eukaryotic interphase nucleus represents a long-standing question in genome organisation [63, 66]. *In vitro,* HPL-2 dimerization induces DCIS of H3K9me from unmodified chromatin arrays (Figure 4). Similar findings were recently independently reported in an *in vitro Drosophila* HP1a–chromatin system, suggesting that DCIS is an intrinsic, conserved property of the HP1–H3K9me axis [97]. Extrapolating from this reductionist framework, we propose that HP1-induced DCIS provides an integral mechanism for the segregation and formation of H3K9me-marked heterochromatin domains *in vivo*. Chromosome-scale modelling and simulations incorporating HP1–H3K9me distribution patterns and HP1 dimerization indeed recapitulate heterochromatin segregation, consistent with the observation of DCIS [98].

### Condensation and transcriptional repression are uncoupled at C. elegans heterochromatin

Although heterochromatin condensation is classically linked to transcriptional silencing [79, 99, 100], our data show that reduction of canonical heterochromatin compaction in HPL-2 dimerization-deficient mutant embryos does not trigger widespread gene derepression (Figure 8F), indicating that heterochromatin condensation and transcriptional repression are mechanistically separable. This observation is in alignment with recent evidence from human and *Drosophila* cells showing that nucleosomes in both eu- and heterochromatic regions remain dynamic and accessible [101], and that disruption of heterochromatin clustering produces only mild transcriptional effects [102]. Further, HP1-driven heterochromatin coalescence was shown to have only mild effects on gene regulation in mammalian cells [103]. In *C. elegans*, repression at H3K9me-marked loci likely relies on additional pathways such as histone deacetylation [53], MES-dependent H3K27me3 deposition [104], and RNAi-mediated silencing [105] that act independently of compaction. Together, these mechanisms likely preserve silencing even when condensation is perturbed, suggesting that HP1-mediated chromatin organization primarily maintains genome architecture.

### Biological significance of heterochromatin condensation

If HPL-2–mediated condensation of heterochromatin exerts limited influence on gene expression, what is its physiological significance? Increasing evidence suggests structural rather than transcriptional functions [106]. HP1 proteins act as chromatin cross-linkers that influence nuclear morphology and chromosome behaviour [107, 108], and defects in these structural roles often result in nuclear blebs, hallmarks of pathologies including progeria, cardiovascular disease, and cancer [109]. In *C. elegans*, ageing is accompanied by nuclear blebbing and loss of peripheral heterochromatin [110]. The shortened lifespan we observe in *hpl-2* dimerization-deficient *C. elegans* might therefore reflect compromised nuclear structure and accelerated ageing resulting from reduced chromatin condensation (Figure 5B,C). The vulval defects in *hpl-2* dimerization mutants emerge predominantly under elevated temperature (Figure 5D–G), suggesting that weakened condensation renders heterochromatin more susceptible to stress-induced remodelling. We propose that HPL-2-mediated bridging fortifies heterochromatin against environmental perturbation, and that its loss exposes the genome to temperature-sensitive reorganisation [111]. In this sense, HP1-driven heterochromatin condensation might act as a structural safeguard that preserves nuclear integrity and genome stability.

## ACKNOWLEDGEMENTS

We thank members of the Frøkjær-Jensen, Erdel, Meister and Fischle laboratories for helpful discussions. N2 and PFR40 *C. elegans* strains were provided by the CGC, which is funded by NIH Office of Research Infrastructure Programs (P40 OD010440). We thank Dr. Elizabeth Miller for generating the *hpl-2b* deletion strain used in this study. We are grateful to Dr. Alfredo De Biasio and Dr. Quoc Phong Nguyen (both at KAUST) for providing access to and training in mass photometry. During preparation and revision of the manuscript, the authors used ChatGPT (OpenAI) and Claude (Anthropic) for language editing and to improve grammar.

## AUTHOR CONTRIBUTIONS

K.E. and W.F. conceived the project with input from Y.O. K.E. executed all *in vitro* experiments. F. R-C. performed *in silico* studies and imaging analysis. K.E and S.A. did all *C. elegans* related experiments with S.E.M. and C. F-J. generating different *C. elegans* strains. F.M. performed FRAP experiments, F.M. and F.E. analysed FRAP experiments. J.S. and P.M. analysed the genomic data. K.E. wrote the original draft of the manuscript which was reviewed and edited by W.F. The final version of the manuscript was read and approved by all authors.

## Supplementary Data statement

Supplementary Data are available at NAR online.

## DATA AVAILABILITY

ChIP-seq data have been deposited to Gene Expression Omnibus (GEO) under the accession number GSE339336. RNA-seq data have been deposited in the EMBL-EBI European Nucleotide Archive (ENA) under the project accession PRJEB102446 and GEO under the accession number GSE342086. The corresponding gene-level raw count matrix is available on Figshare (https://doi.org/10.6084/m9.figshare.30559331). Scripts used for ChIP-seq and RNA-seq analyses have been archived on Zenodo (https://doi.org/10.5281/zenodo.22099313 and https://doi.org/10.5281/zenodo.22099356, respectively).

