## Supplementary Tables and Figures for "HP1 dimerization drives the condensation and segregation of H3K9me-marked chromatin"

#### Table S1. Plasmids

| Name | Vector | Resistance | Application | Source |
|---|---|---|---|---|
| H2A | pET-3a | Ampicillin | Octamer assembly | K. Luger, University of Colorado Boulder[1] |
| H2B | pET-3a | Ampicillin | | |
| H3 | pET-3a | Ampicillin | | |
| H4 | pET-3a | Ampicillin | | |
| H3 Δ1-20 A21C | pET-3a | Ampicillin | Native chemical ligation | Fischle laboratory[2] |
| H2B A1C | pET-3a | Ampicillin | Fluorophore labelling | This study |
| HPL-2A | pET-16b | Ampicillin | *In vitro* assays | |
| HPL-2A I158A | pET-16b | Ampicillin | *In vitro* assays | |
| HPL-2 9-patch neutral hinge | pET-28a(+) | Kanamycin | *In vitro* assays | |
| HPL-2 Δhinge | pET-28a(+) | Kanamycin | *In vitro* assays | |
| HPL-2 RKR neutral hinge | pET-28a(+) | Kanamycin | *In vitro* assays | |
| HPL-2 scrambled hinge | pET-28a(+) | Kanamycin | *In vitro* assays | |
| HPL-2 total neutral hinge | pET-28a(+) | Kanamycin | *In vitro* assays | |
| HPL-2 I158A SpyTag | pET-28a(+) | Kanamycin | *In vitro* assays | |
| HPL-2 I158A SpyCatcher | pET-28a(+) | Kanamycin | *In vitro* assays | |
| HPL-2 2xhinge I158A | pET-28a(+) | Kanamycin | *In vitro* assays | |
| HPL-2 3xhinge I158A | pET-28a(+) | Kanamycin | *In vitro* assays | |
| HPL-2 H2B.8 hinge I158A | pET-28a(+) | Kanamycin | *In vitro* assays | |
| HPL-2 DDX4 hinge I158A | pET-28a(+) | Kanamycin | *In vitro* assays | |
| MBP-HPL-2 CSD | pET-28a(+) | Kanamycin | Mass photometry | |
| MBP-HPL-2 CSD I158A | pET-28a(+) | Kanamycin | Mass photometry | |

#### Table S2. Histone peptides

| Name | Histone (aa) | Peptide sequence | Application | Source |
|---|---|---|---|---|
| H3K9me3 | H3 (1-20) | ARTKQTARK(me3)ST GGKAPRKQL(thioester) | Native protein ligation | NMR laboratories, Max Planck Institute for Biophysical Chemistry, Göttingen |
| H3K9me0 | H3 (1-20) | ARTKQTARK(me0)ST GGKAPRKQL(biotin) | Biotinylated peptides used for peptide pulldowns | |
| H3K9me2 | H3 (1-20) | ARTKQTARK(me2)ST GGKAPRKQL(biotin) | | |
| H3K9me3 | H3 (1-20) | ARTKQTARK(me3)ST GGKAPRKQL(biotin) | | |

### Table S3. *C. elegans* strains

| Strain | Genotype | Designation | Source |
|---|---|---|---|
| N2 | *wild-type* | N2 | CGC [3] |
| PFR40 | *hpl-2(tm1489) III* | *hpl-2Δ* | CGC [4] |
| WFK2 | *lin-61(tm2649) I* | *lin-61Δ* | [5] |
| WFK3 | *lin-61(tm2649) I; hpl-2(tm1489) III* | *lin-61Δ; hpl-2Δ* | This study, Crossing |
| PHX9175 | *lin-61 (tm2649) I; hpl-2a(syb3274) III* | *lin-61Δ*; *hpl-2::gfp* | This study, SunyBiotech |
| PHX9304 | *lin-61(tm2649) I; hpl-2a(syb3274 syb9304) III* | *lin-61Δ*; *hpl-2 total neutral hinge::gfp* | This study, SunyBiotech |
| PHX9193 | *lin-61(tm2649) I; hpl-2a(syb3274 syb6241) III* | *lin-61Δ*; *hpl-2 I158A::gfp* | This study, SunyBiotech |
| PHX9332 | *lin-61(tm2649) I; hpl-2a(syb3274 syb6241 syb9332) III* | *lin-61Δ*; *hpl-2 3xhinge I158A::gfp* | This study, SunyBiotech |
| PHX9185 | *lin-61(tm2649) I; hpl-2a(syb9185) III* | *lin-61Δ*; *hpl-2 total neutral hinge* | This study, SunyBiotech |
| PHX10290 | *hpl-2a(syb10290) III* | *hpl-2 3xhinge I158A* | This study, SunyBiotech |
| WFK4 | *lin-61(tm2649) I; hpl-2a(syb10290) III* | *lin-61Δ*; *hpl-2 3xhinge I158A* | This study, Crossing |
| WFK5 | *hpl-2a(I158A) III* | *hpl-2 I158A* | This study, CRISPR/Cas9 editing |
| WFK6 | *lin-61(tm2649) I; hpl-2a(I158A) III;* | *lin-61Δ*; *hpl-2 I158A* | This study, Crossing |

### Table S4. Primers for *C. elegans* genotyping

| Gene | Mutation | WT (bp) | Mutant (bp) | Primer (5' – 3') |
|---|---|---|---|---|
| *hpl-2* | *tm1489* deletion | 1142 | 470 | CTGCTTGCCTTCCAGTGAGA |
| | | | | GTGGAAGGGCTTGAAGGTG |
| | *syb3274* insertion | 824 | 1550 | GAATCTTGTGGTCTCGTGA |
| | | | | TTTGGACCTTGTGTAGAGA |
| | *syb9304* insertion | 434 | - | CAGCCAGAAGCCCGAACCT |
| | | | | CCATTTCAAAACTCGAGC |
| | | - | 434 | TTCTCAAGCACCAGCTCCA |
| | | | | CCATTTCAAAACTCGAGC |
| | *syb6241* substitution | 597 | - | CAGCTATCCGAGCCAAGTTAT |
| | | | | ACGTTGTGGCGAATTTTGAA |
| | | - | 597 | CAGCTATCCAAGTCAGGTGGCT |
| | | | | ACGTTGTGGCGAATTTTGAA |
| | *syb9332* or *syb10290* insertion | 833 | 1019 | TCTCATACAGTGGCAAGGAT |
| | | | | CCATTTCAAAACTCGAGC |
| *lin-61* | *tm2649* deletion | 1322 | 650 | CGATAAGACCTACTTGTGGG |
| | | | | TTGAACTCATCTGGCGGAAC |

**Table S5. Antibodies**

| Name | Host | Application | Dilution/Volume | Supplier |
|---|---|---|---|---|
| **Primary antibodies** | | | | |
| αH3 | Rabbit | Western blot | 1:2500 | Abcam (ab1791) |
| αH3K9me2 | Mouse | Western blot<br>Immunofluorescence<br>ChIP | 1:2500<br>1:250<br>4 µL per reaction | Abcam (ab1220) |
| αH3K9me3 | Rabbit | Western blot | 1:2500 | Abcam (ab8898) |
| αGFP | Rabbit | Western blot<br>Immunofluorescence<br>ChIP | 1:1000<br>1:250<br>4 µL per reaction | Invitrogen (A-11122) |
| IgG | Mouse | ChIP | 4 µL per reaction | Diagenode (C15400001) |
| IgG | Rabbit | ChIP | 4 µL per reaction | Diagenode (C15410206) |
| **Secondary antibodies** | | | | |
| αMouse IgG, 680RD | Goat | Western blot | 1:5000 | LICOR (925-68070) |
| αRabbit IgG, 680RD | Goat | Western blot | 1:5000 | LICOR (925-68071) |
| αMouse IgG, 800CW | Goat | Western blot | 1:5000 | LICOR (925-32210) |
| αRabbit IgG, 800CW | Goat | Western blot | 1:5000 | LICOR (925-32211) |
| αRabbit IgG, Alexa Fluor 488 | Donkey | Immunofluorescence | 1:250 | Invitrogen (A-21206) |
| αMouse IgG, Alexa Fluor 647 | Goat | Immunofluorescence | 1:250 | Invitrogen (A-32728) |

**Table S6. Genomic loci analysed by ChIP-qPCR.**

| Chromosome loci (bp) | Targets | Primer (5' – 3') |
|---|---|---|
| chrI:1444214-1444402 | Target 1 | CAGCGCACACTTTTCACATT<br>CGACCAGAACTCGCTCTTTT |
| chrI:1435588-1435774 | Target 2 | GTTTCCGCAAATCGTGTTCT<br>TGGTTTGCAAAAATAAATTACAAAA |
| chrI:14113157-14113336 | Target 3 | AAATATCGAAAATAACGTTGAAAAA<br>TTTGAGTTAAAAATCACGAAAAACC |
| chrII:2920879-2921085 | Target 4 | CCGACGATTTTCTGAGGAAT<br>TCGTCGAAAACAGTGATGAAA |
| chrII:2973992-2974173 | Target 5 | TTTCACGGACATTGAAAGGA<br>CTCCCTCGTCTAGCCTCTCC |
| chrII:13447717-13447912 | Target 6 | TCTGGTAAATCGGGTAGATCAAA<br>GCCCCACTGAGAATCTTGTT |
| chrIII:1215307-1215517 | Target 7 | TCGGCAGACTCATTAAGAAAAA<br>AGAACCTGAACCCCGAAACT |
| chrIII:1418802-1418983 | Target 8 | AAATCGGAGCCAAATCTCG<br>CAGTGATTTTTGTGCGAAACA |
| chrIII:12612443-12612630 | Target 9 | TCAGCCTTGAAAGTCCCTTTT<br>AGATTGTGGAGACGCCAGAG |

| | | |
|---|---|---|
| chrIV:2602065-2602270 | Target 10 | TTCCATTTTCCCGCCTATAA |
| | | CGCTCTCATAAGGTCGTCATC |
| chrIV:2774597-2774799 | Target 11 | CTCTTTCCAAAACCATCGAAA |
| | | TTGACACAAAGAACCCTCACA |
| chrIV:16491612-16491791 | Target 12 | GAGACCCTCATTTCCCCATC |
| | | TTTGAGAGAGCGCAGACAGA |
| chrV:5049502-5049686 | Target 13 | TGCTTGCGTACCTTTTCTCC |
| | | TCAGCGACCAACAGTGATGT |
| chrV:4827080-4827263 | Target 14 | GGGCTTTTGCATCAACTTTC |
| | | TTTTCGCCAAAAATAAAGCA |
| chrV:19643086-19643271 | Target 15 | TAACGAGCGTGGGATCAAAT |
| | | CCCTCCCTCACCTCGTACA |
| chrI:7778160-7778348 | Off target 1 | TTGAAATTTGCCCTGAACATT |
| | | AAATCTGCCTGTGGGAAGAC |
| chrII:7120608-7120796 | Off target 2 | TCAGGGGCGAGATTATTTGT |
| | | CATCAGCAAGCAGCTCAACT |
| chrIII:6783755-6783965 | Off target 3 | TTTGCGATTCTTTGTCACTCA |
| | | TCTGCCGTACTTCCGCTATT |
| chrIV:10027826-10028005 | Off target 4 | CTGTTTTTAAACTCAAACACGTGTAAC |
| | | TCAAGCCTATACTTTGCCAAAAA |
| chrV:10027662-10027860 | Off target 5 | CTCTTCGCTCGACTCGTTCT |
| | | TAGAGGGGATGCAGAGGGTA |

**Table S7. Effects of HPL-2 mutations on *C. elegans* embryonic viability and sterility**

Scoring of embryonic viability and sterility in *C. elegans* expressing HPL-2::GFP wild-type or mutant proteins in the background of *lin-61Δ* at 20°C or 25°C. Data are representative of 40 worms scored per strain from four independent experiments. SD, standard deviation.

| **Strain** | **20°C** | | **25°C** | |
|---|---|---|---|---|
| | **Viability**±SD (%) | **Sterility (%)** | **Viability**±SD (%) | **Sterility (%)** |
| N2 | 97.5±0.5 | 0 | 96.2±0.9 | 0 |
| *hpl-2Δ* | 95.3±0.9 | 7.5 | 94.5±1.2 | 17.5 |
| *lin-61Δ* | 96.4±0.5 | 0 | 94.3±1.0 | 0 |
| *lin-61Δ; hpl-2Δ* | 83.8±3.1 | 12.5 | 82.0±2.7 | 75 |
| *lin-61Δ; hpl-2::gfp* | 96.1±0.7 | 0 | 94.0±0.8 | 2.5 |
| *lin-61Δ; hpl-2 total neutral hinge::gfp* | 95.4±0.7 | 0 | 94.2±0.8 | 2.5 |
| *lin-61Δ; hpl-2 3xhinge I158A::gfp* | 95.2±0.6 | 0 | 93.3±2.0 | 20 |
| *lin-61Δ; hpl-2 I158A::gfp* | 95.3±0.7 | 2.5 | 94.0±1.1 | 30 |

**Table S8. Effects of HPL-2 mutations on larval arrest at elevated temperature**

Scoring of high temperature arrest (HTA) phenotype in *C. elegans* expressing HPL-2::GFP wild-type or mutant proteins in the background of *lin-61Δ* at 24°C or 26°C. n, total number of worms scored in three independent experiments.

| Genotype | Percentage of arrested larvae at 24°C (n) | Percentage of arrested larvae at 26°C (n) |
|---|---|---|
| N2 | 0.0 (543) | 0.0 (497) |
| *hpl-2Δ* | 0.0 (582) | 80.2 (524) |
| *lin-61Δ* | 0.0 (621) | 12.8 (598) |
| *lin-61Δ; hpl-2::gfp* | 0.0 (558) | 10.5 (601) |
| *lin-61Δ; hpl-2 total neutral hinge::gfp* | 0.0 (587) | 14.7 (622) |
| *lin-61Δ; hpl-2 3xhinge I158A::gfp* | 0.0 (612) | 17.2 (643) |
| *lin-61Δ; hpl-2 I158A::gfp* | 0.0 (583) | 21.4 (672) |

* *The lin-61Δ; hpl-2Δ double mutant could not be analysed because of severe embryonic lethality at 26°C, precluding HTA scoring.*

**Table S9. Effects of HPL-2 mutations on *C. elegans* vulval development**

Scoring of vulval defects of *C. elegans* expressing HPL-2::GFP wild-type or mutant proteins in the background of *lin-61Δ* at 20°C or 25°C. n, total number of worms scored in two independent experiments. SD, standard deviation.

| Strain | 20°C | | 25°C | |
|---|---|---|---|---|
| | Vulval defects±SD (%) | n | Vulval defects±SD (%) | n |
| N2 | 0±0 | 794 | 0±0 | 859 |
| *hpl-2Δ* | 0±0 | 809 | 0.9±0.2 | 794 |
| *lin-61Δ* | 0±0 | 804 | 0.3±0.0 | 797 |
| *lin-61Δ; hpl-2Δ* | 0.6±0.2 | 818 | 53.4±4.1 | 814 |
| *lin-61Δ; hpl-2::gfp* | 0±0 | 800 | 0.6±0.2 | 823 |
| *lin-61Δ; hpl-2 total neutral hinge::gfp* | 0±0 | 812 | 9.1±3.2 | 798 |
| *lin-61Δ; hpl-2 3xhinge I158A::gfp* | 0±0 | 835 | 40.4±5.0 | 842 |
| *lin-61Δ; hpl-2 I158A::gfp* | 0.5±0.3 | 825 | 37.8±3.2 | 840 |

### Supplementary figures

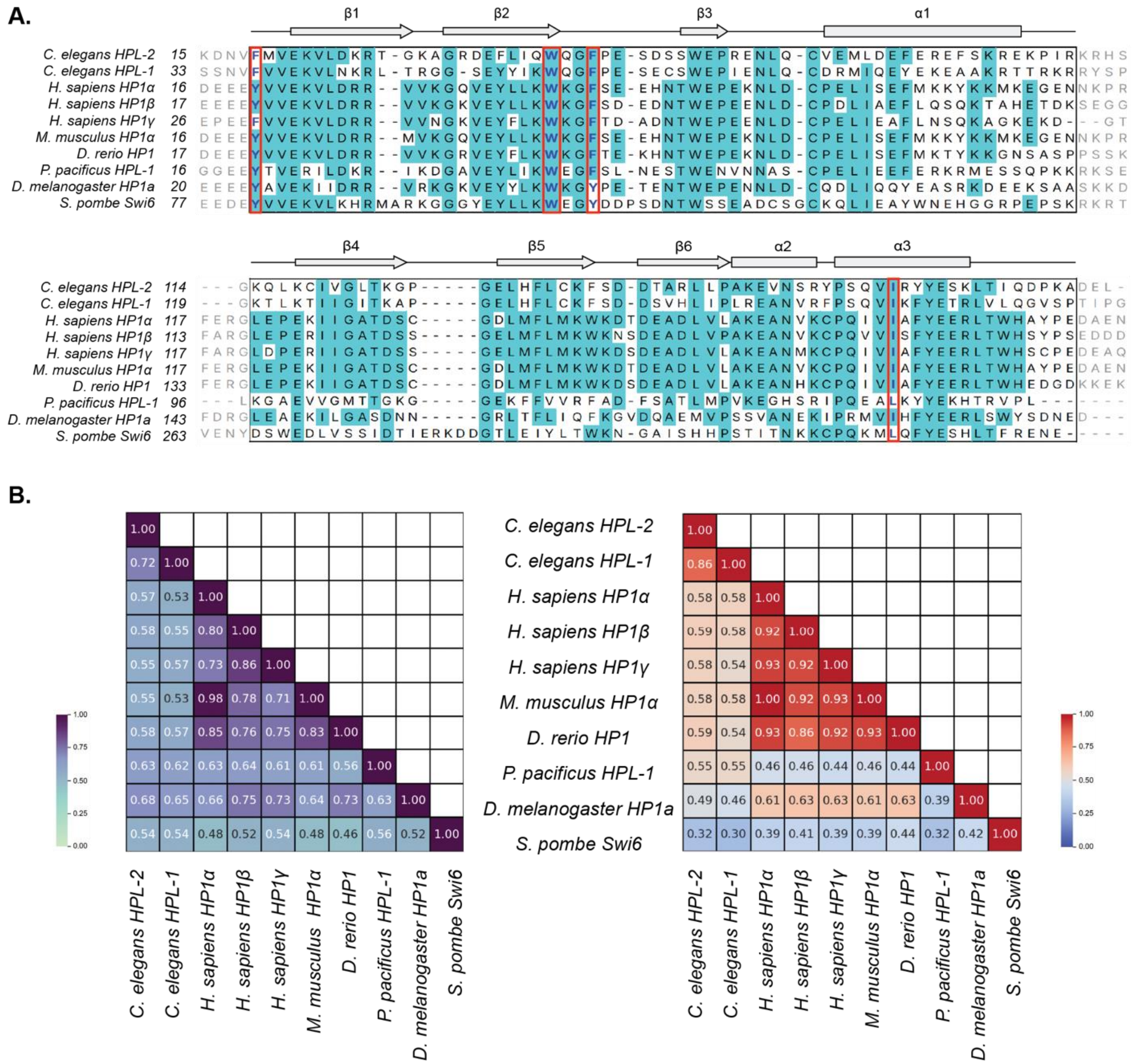

**Figure S1. Sequence conservation of HPL-2 and HP1 orthologues.**

**(A)** Multiple sequence alignment of HP1 proteins from different species. Secondary structure elements of HPL-2 are annotated based on AlphaFold prediction. Top, chromodomain; bottom, chromoshadow domain. Residues with >50% identity across orthologues are highlighted. Red boxes indicate conserved aromatic cage residues within the chromodomain and the conserved isoleucine residue important for chromoshadow domain dimerization. **(B)** Pairwise similarity scores of different HP1 proteins for the chromodomains (left) and chromoshadow domains (right), based on residue-level hydrophobicity.

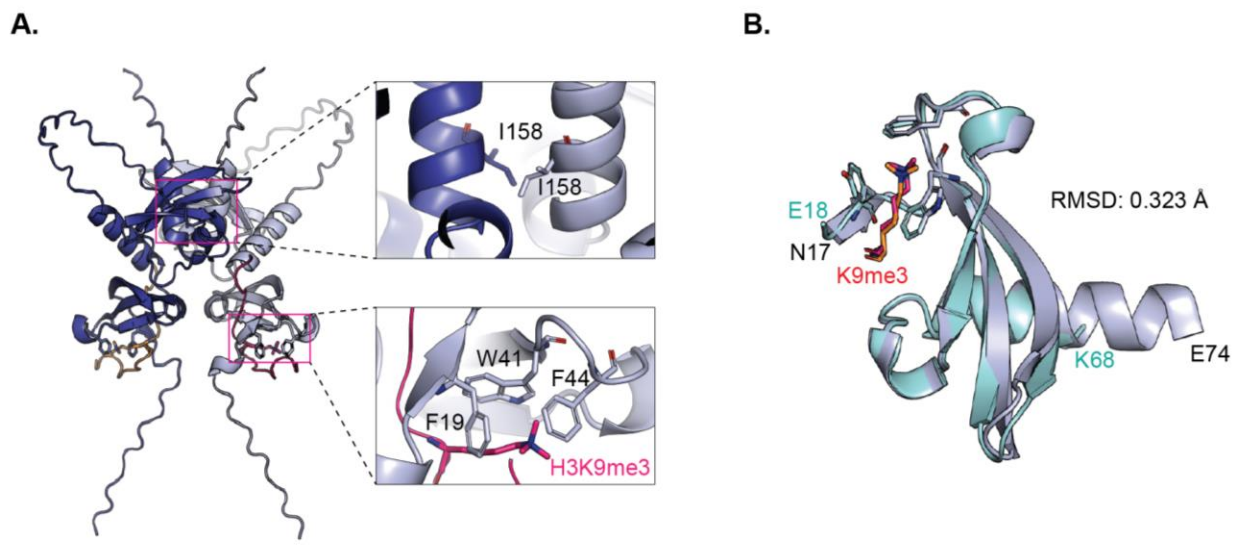


**Figure S2. Structural models of HPL-2 dimerization and H3K9me3 recognition**

**(A)** AlphaFold-predicted dimeric structure of *C. elegans* HPL-2 (blue and grey) docked with H3 tails (orange and pink). Insets show a magnified view of the isoleucine-mediated dimerization interface (top) and interaction between aromatic cage residues (grey) and H3K9me3 (pink) (bottom). **(B)** Structural alignment of AlphaFold-predicted *C. elegans* HPL-2 CD (grey) with human HP1α CD (green) bound to H3K9me3 peptide (pink) (PDB: 3FDT). N17 and E18 mark the N-termini, while E74 and K68 mark the C-termini of the chromodomains of HPL-2 and HP1α, respectively.

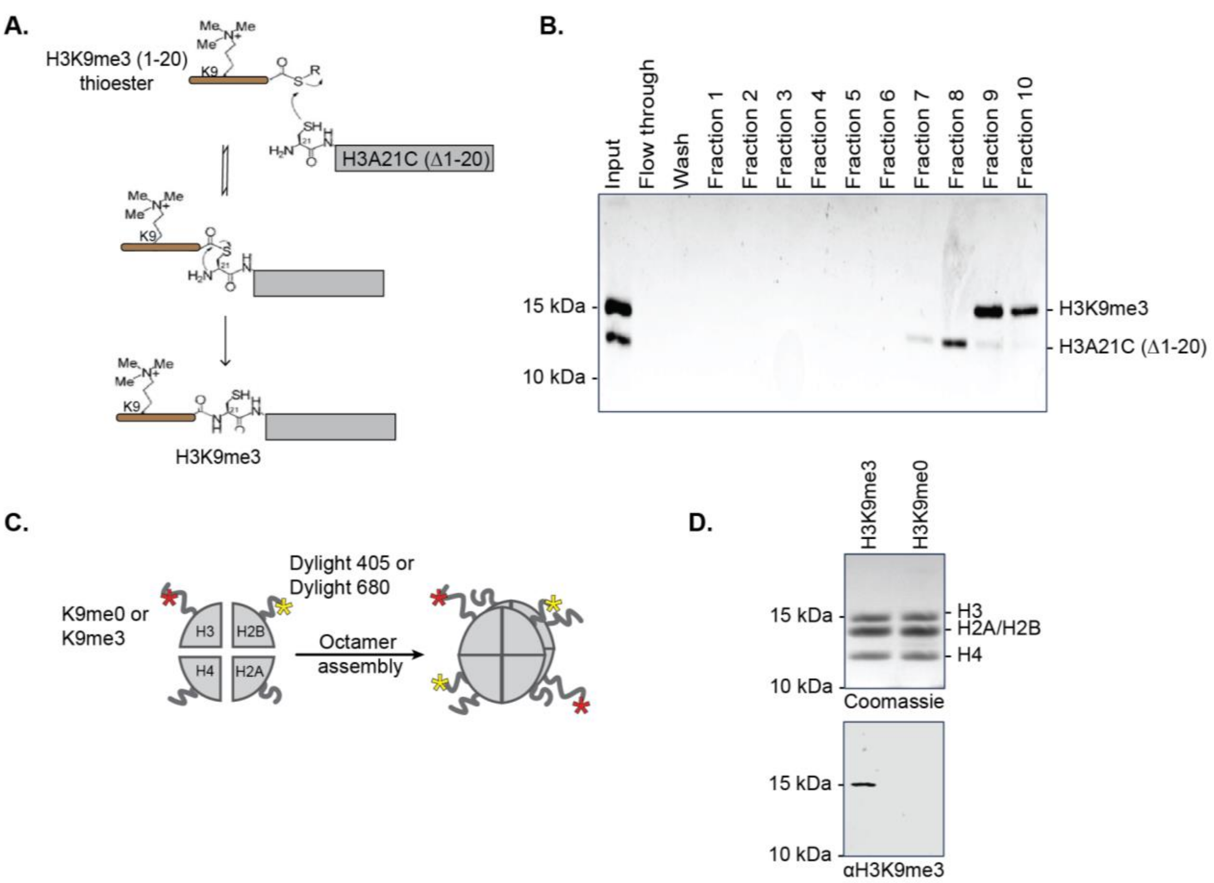


**Figure S3. Assembly of recombinant histone octamers**

**(A**) Scheme of native chemical ligation (NCL) for generating H3K9me3 histone. In the first step, an N-terminal synthetic H3 peptide comprising the first 20 amino acids and carrying the K9me3 modification, with a C-terminal thioester, is coupled to the thiol group of cysteine 21 in recombinantly expressed H3A21C (Δ1–20) via reversible trans-thioesterification. The reaction proceeds through a spontaneous and irreversible rearrangement of the thioester to form a native peptide bond. Scheme adapted from [6]. **(B)** SDS-PAGE analysis of NCL reaction separated by ion-exchange chromatography (IEX). Input, NCL reaction products; flowthrough, material that did not bind the IEX resin; wash, non-specific eluates from IEX column; fractions n, material eluting from the IEX with increasing salt concentration. The running positions of molecular weight markers are indicated on the left and those of unligated H3A21C (Δ1–20) (unligated H3Δ) and full-length, ligated H3K9me3 (ligated H3) are indicated on the right. **(C)** Scheme of octamer assembly. H3K9me0 histones with Dylight 405-labelled H2B or H3K9me3 histones with Dylight 680-labelled H2B, together with H2A and H4, were assembled into histone octamers. **(D)** H3K9me3 and H3K9me0 octamers assembled from recombinant *Xenopus laevis* histones were run on SDS-PAGE and stained with Coomassie Blue (top). Western blot analysis of octamers using an antibody specific for H3K9me3 (bottom). The running positions of molecular weight markers are indicated on the left and those of core histone proteins on the right.

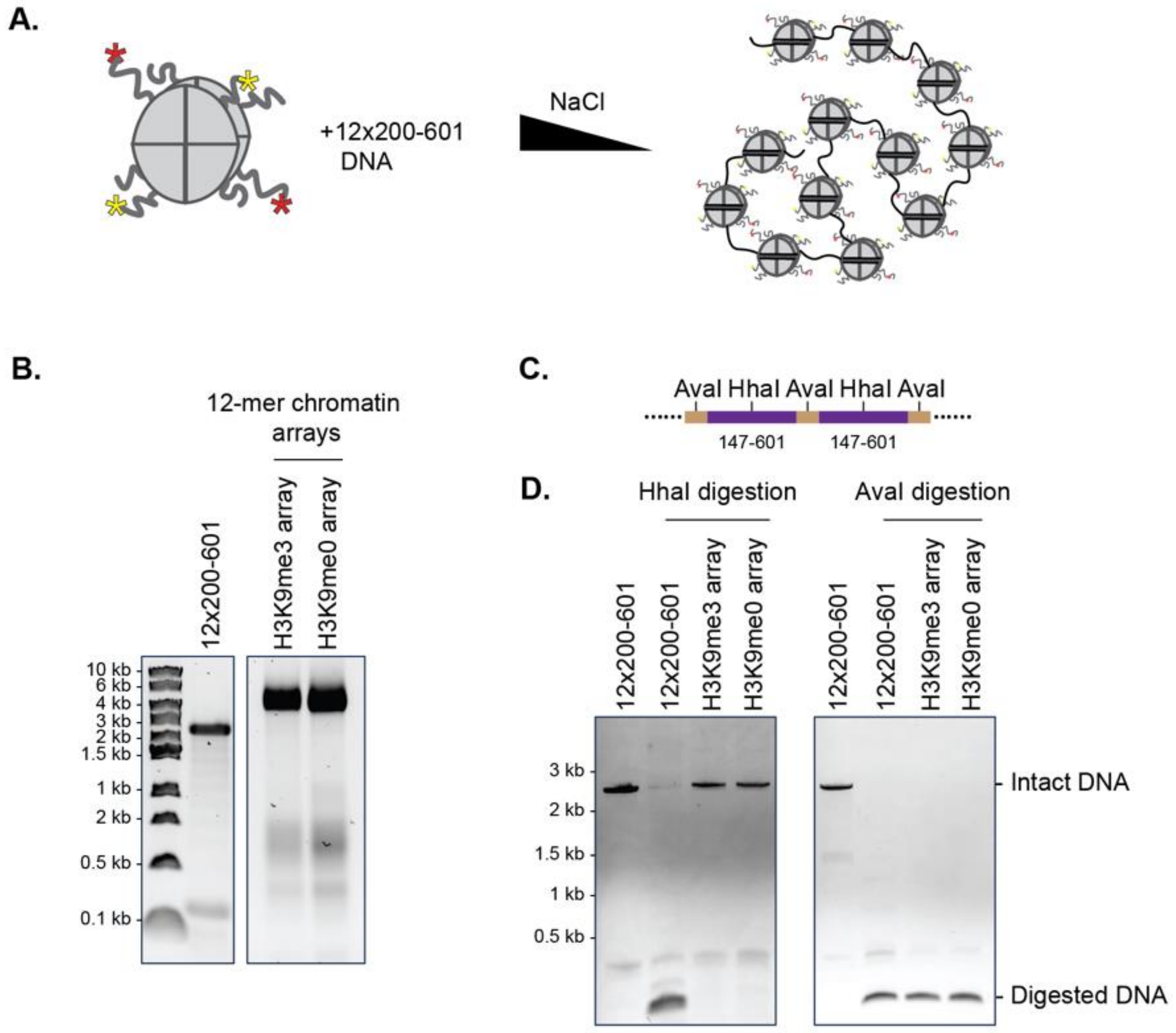


**Figure S4. Reconstitution of recombinant, H3K9me3 chromatin arrays**

**(A)** Scheme of reconstitution of chromatin arrays by salt-gradient dialysis using octamers and DNA containing 12 repeats of the 147 bp nucleosome positioning 601 sequence with 53 bp linker DNA (12x200-601). **(B)** 12x200-601 control DNA and reconstituted chromatin arrays were run on a native agarose gel and stained with ethidium bromide. The running positions of molecular size markers are indicated on the left. **(C)** Scheme of AvaI and HhaI restriction endonuclease recognition sites in the 12x200-601 DNA. The Widom 601 147 bp nucleosome positioning sequence is highlighted in purple, and the 53 bp linker region is highlighted in brown. **(D)** Reconstituted chromatin arrays were digested with AvaI (right) or HhaI (left) and the DNA isolated from the reactions was run on native agarose gels that were stained with ethidium bromide. The running positions of molecular size markers are indicated on the left and of major DNA species in the reactions on the right.

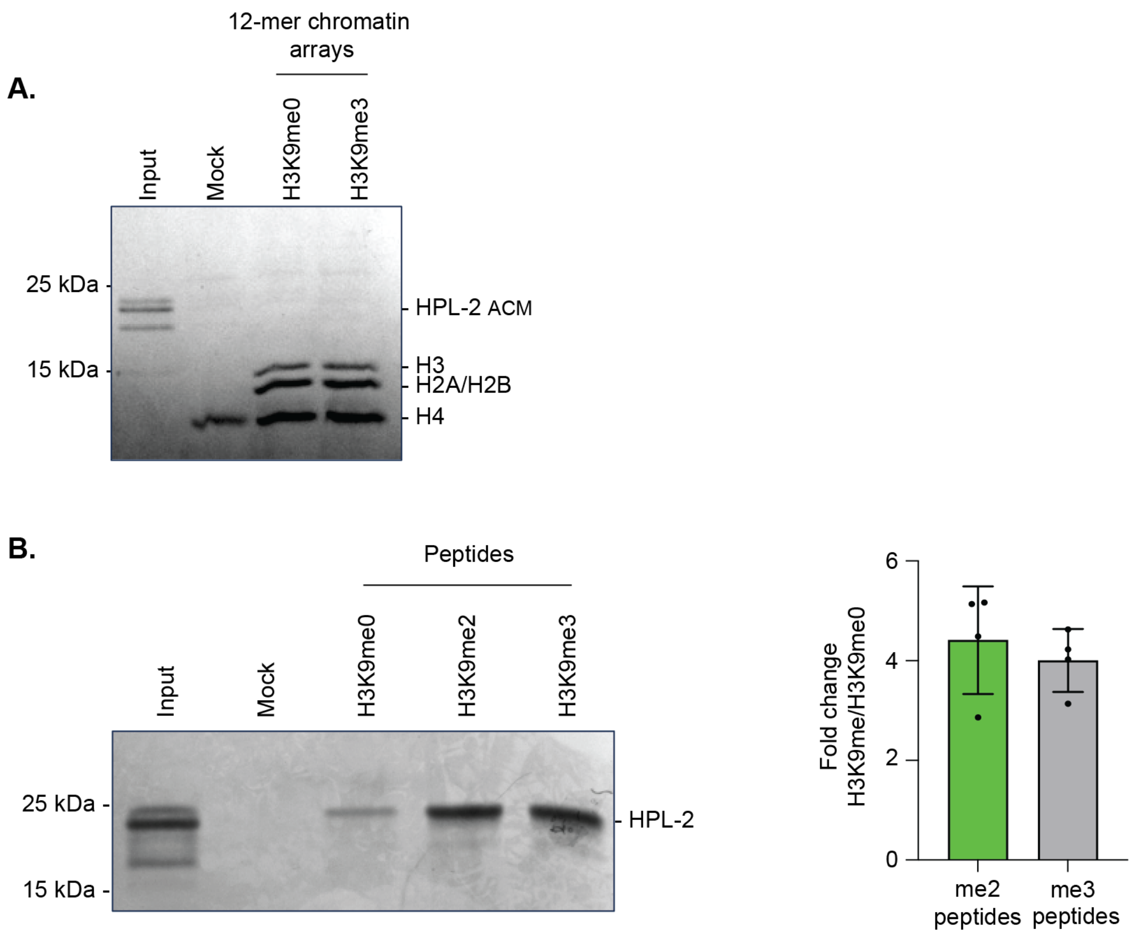


**Figure S5. Analysis of HPL-2 CD–H3K9me interaction**

**(A)** Recombinant HPL-2 ACM was incubated with biotinylated 12-mer H3K9me0 or H3K9me3 chromatin arrays immobilized on streptavidin beads. Bound proteins were analysed by SDS–PAGE and stained with Coomassie blue. Running positions of molecular weight markers, histones, and HPL-2 ACM are indicated. “Mock” indicates control pull-downs with streptavidin beads to assess non-specific binding; “Input” represents 10% of total HPL-2 ACM used. **(B)** Left: Recombinant HPL-2 was incubated with biotinylated H3K9me0, H3K9me2 or H3K9me3 peptides immobilized on streptavidin beads. Bound proteins were analysed by SDS–PAGE and stained with Coomassie blue. Running positions of molecular weight markers and HPL-2 are indicated. “Mock” indicates control pull-downs with streptavidin beads to assess non-specific binding; "Input" represents 10% of the total HPL-2 used. Right: Quantification of peptide pulldowns as shown on the left. Data represent mean ± SD from four independent experiments.

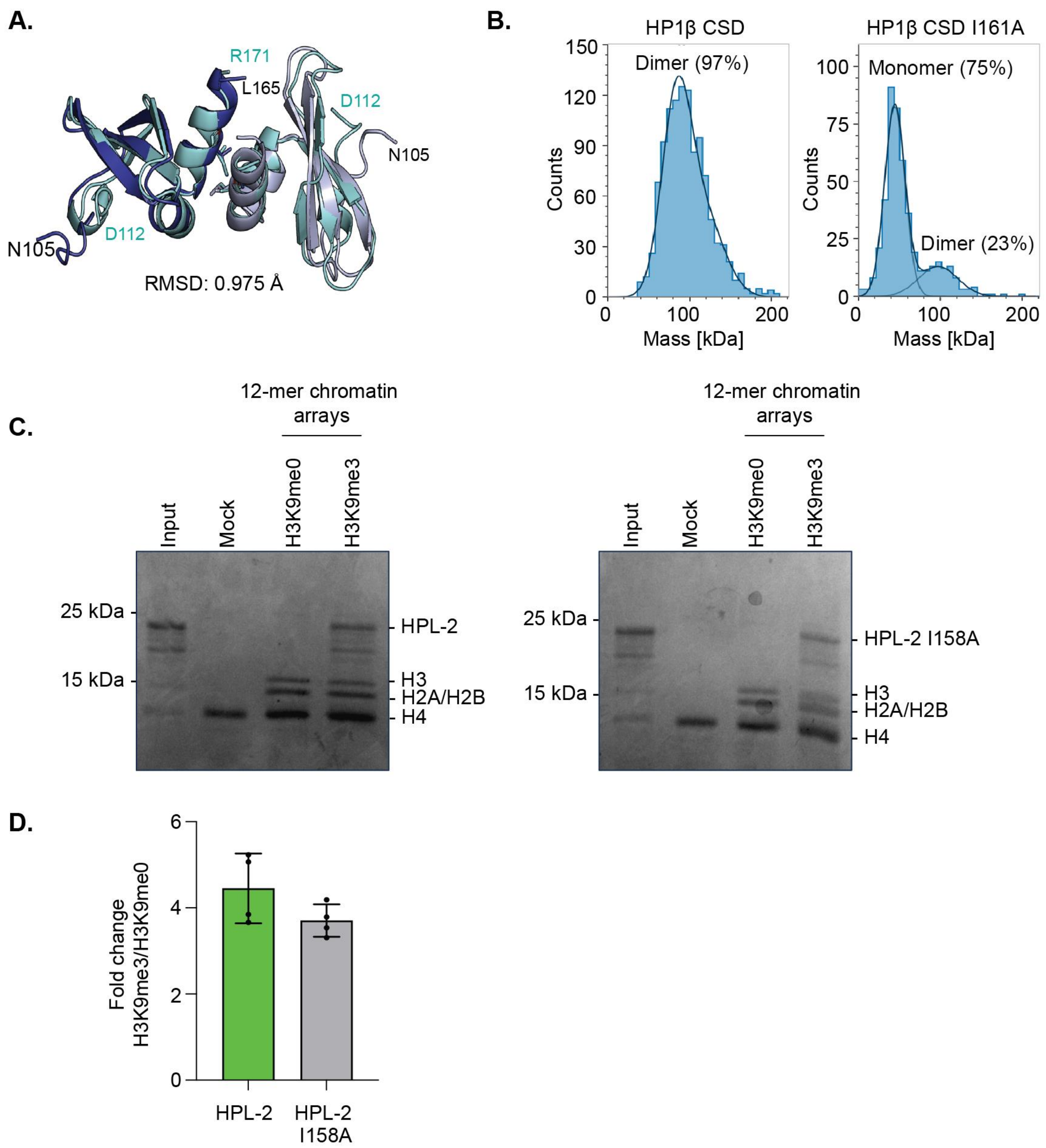


**Figure S6. Analysis of HPL-2 CSD dimerization**

(**A**) Structural alignment of the AlphaFold-predicted *C. elegans* HPL-2 CSD dimer (grey) with the human HP1α CSD dimer (green) (PDB: 3I3C). N105 and D112 mark the N-termini, while L165 and R171 mark the C-termini of the chromoshadow domains of HPL-2 and HP1α, respectively. **(B)** Mass photometry analysis of recombinant MBP-tagged CSDs of wild-type and mutant human HP1β proteins. Particle counts (y-axis) are plotted as a function of molecular mass (x-axis). Data are representative of three independent experiments. **(C)** Recombinant HPL-2 (left) or HPL-2 I158A mutant proteins (right) were incubated with biotinylated 12-mer H3K9me0 or H3K9me3 chromatin arrays immobilized on streptavidin beads. Bound proteins were separated by SDS–PAGE and stained with Coomassie blue. The running positions of molecular weight markers are indicated on the left and those of core histones and HPL-2 proteins on the right. “Mock” indicates control pull-downs with streptavidin beads to assess non-specific binding; “Input” represents 10% of total protein used. **(D)** Quantification of chromatin array pulldowns from (C) normalized to histones H2A/H2B. Data represent mean ± SD from three independent experiments.

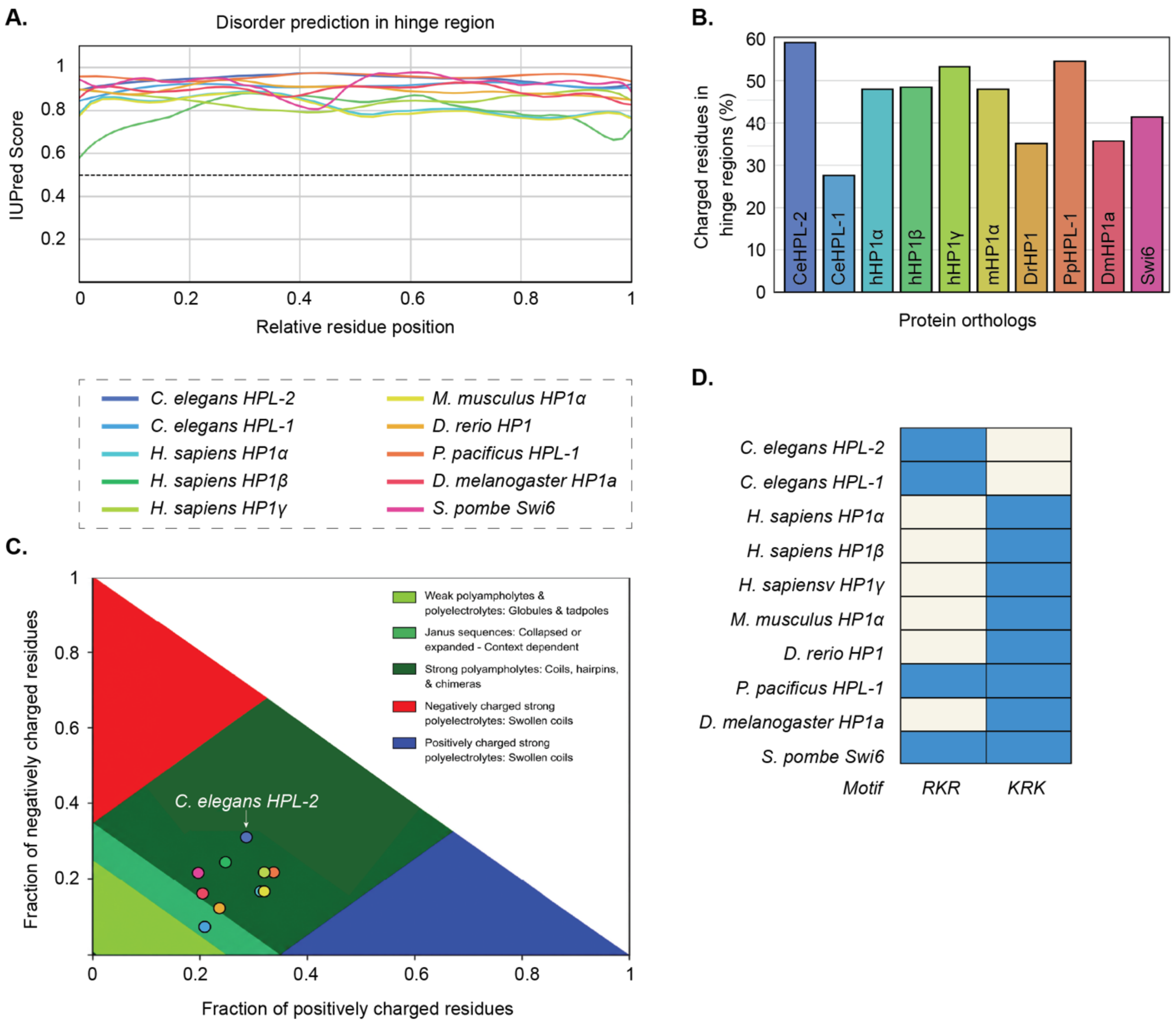


**Figure S7. Intrinsic disorder and charge distribution in the hinge regions of HP1 orthologs (A)** IUPred3 disorder prediction scores across the hinge regions of HP1 ortholog proteins Profiles are plotted against normalized residue positions (x-axis). Dashed line indicates a disorder score of 0.5. Legend indicates protein identities by colour. **(B)** Percentage of charged residues (Asp, Glu, Lys, Arg) in the hinge region of HP1 ortholog proteins, calculated at pH 7.4. **(C)** CIDER (Classification of Intrinsically Disordered Ensemble Regions) analysis of HP1 hinge regions, classifying intrinsically disordered sequences according to their sequence-ensemble properties. See legend below Figure S6A for annotation of protein identities. **(D)** KRK and RKR polybasic motifs identified within the hinge region of each HP1 orthologue. Colour scheme: blue, present; white, absent.

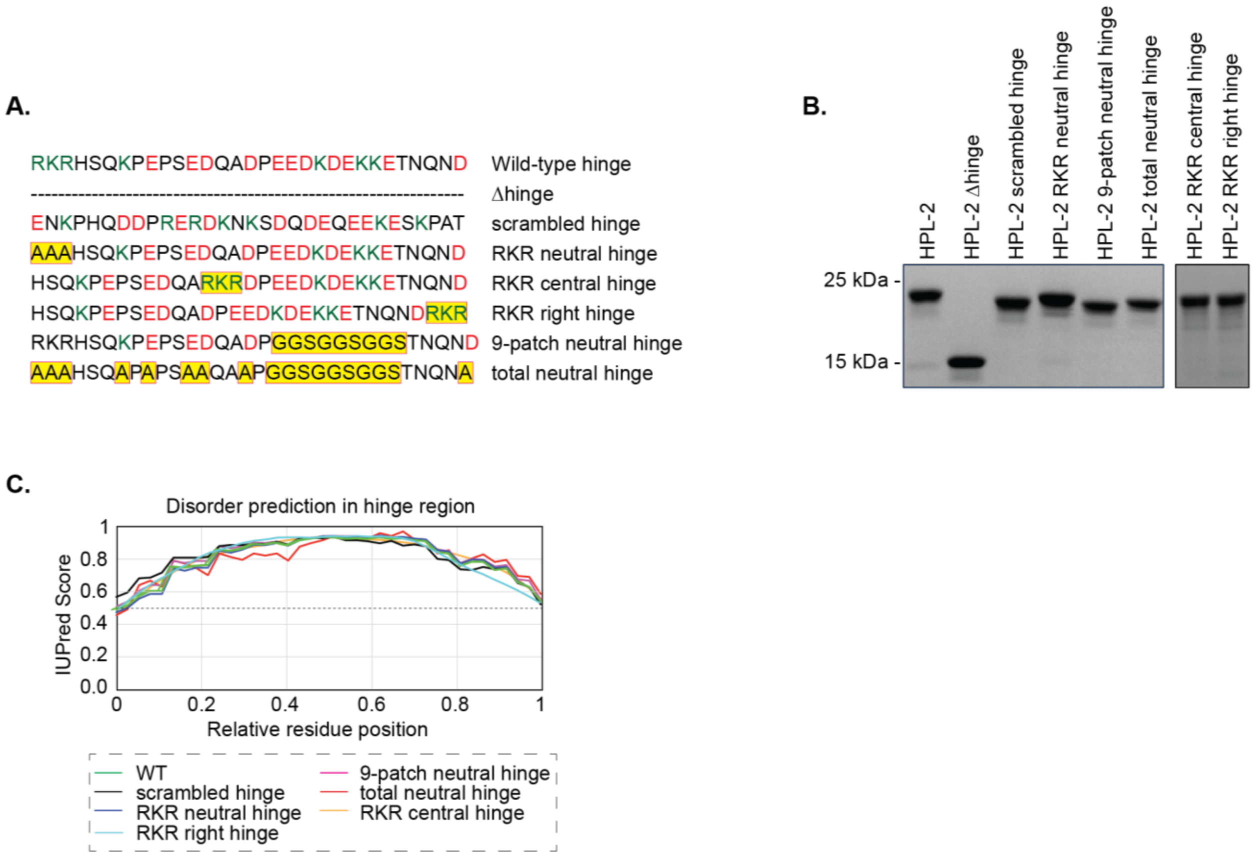


**Figure S8. Design and characterization of HPL-2 hinge region mutants**

**(A**) Annotation of different mutations introduced in the hinge region of HPL-2. Charge neutralised residues and repositioned RKR motifs are highlighted in yellow. **(B)** Recombinant HPL-2 wild-type and mutant proteins were run on SDS–PAGE and stained with Coomassie blue. **(C)** IUPred3 disorder prediction scores across the hinge region of HPL-2 wild-type and mutant proteins. Profiles are plotted against normalized residue positions (x-axis). Dashed line indicates a disorder score of 0.5.

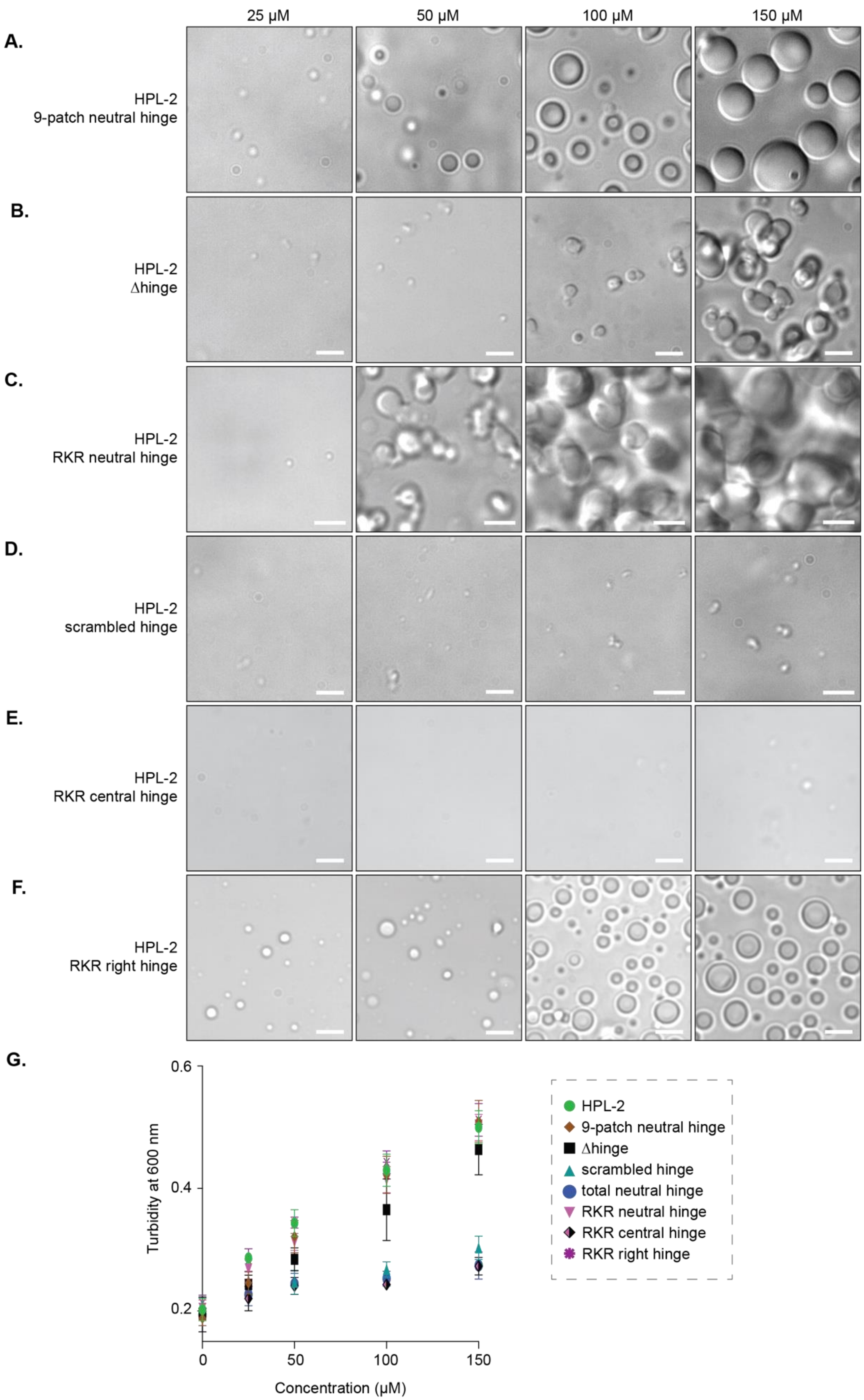

25 μM
50 μM
100 μM
150 μM
A.
HPL-2
9-patch neutral hinge
B.
HPL-2
Δhinge
C.
HPL-2
RKR neutral hinge
D.
HPL-2
scrambled hinge
E.
HPL-2
RKR central hinge
F.
HPL-2
RKR right hinge
G.
Turbidity at 600 nm
0.6
0.4
0.2
0
50
100
150
Concentration (μM)
HPL-2
9-patch neutral hinge
Δhinge
scrambled hinge
total neutral hinge
RKR neutral hinge
RKR central hinge
RKR right hinge

**Figure S9. Phase separation properties of HPL-2 proteins mutant in the hinge region**

**(A–F)** Differential interference contrast (DIC) microscopy of condensates formed by the indicated HPL-2 mutant proteins at increasing concentrations. Data are representative of three independent experiments. Scale bars: 5 μm. **(G)** Plot of turbidity measured by absorbance reading at 600 nm against increasing concentrations of wild-type and mutant HPL-2 proteins. Condensate formation was induced as in (A–F). Data represent mean ± SD from three independent replicates.

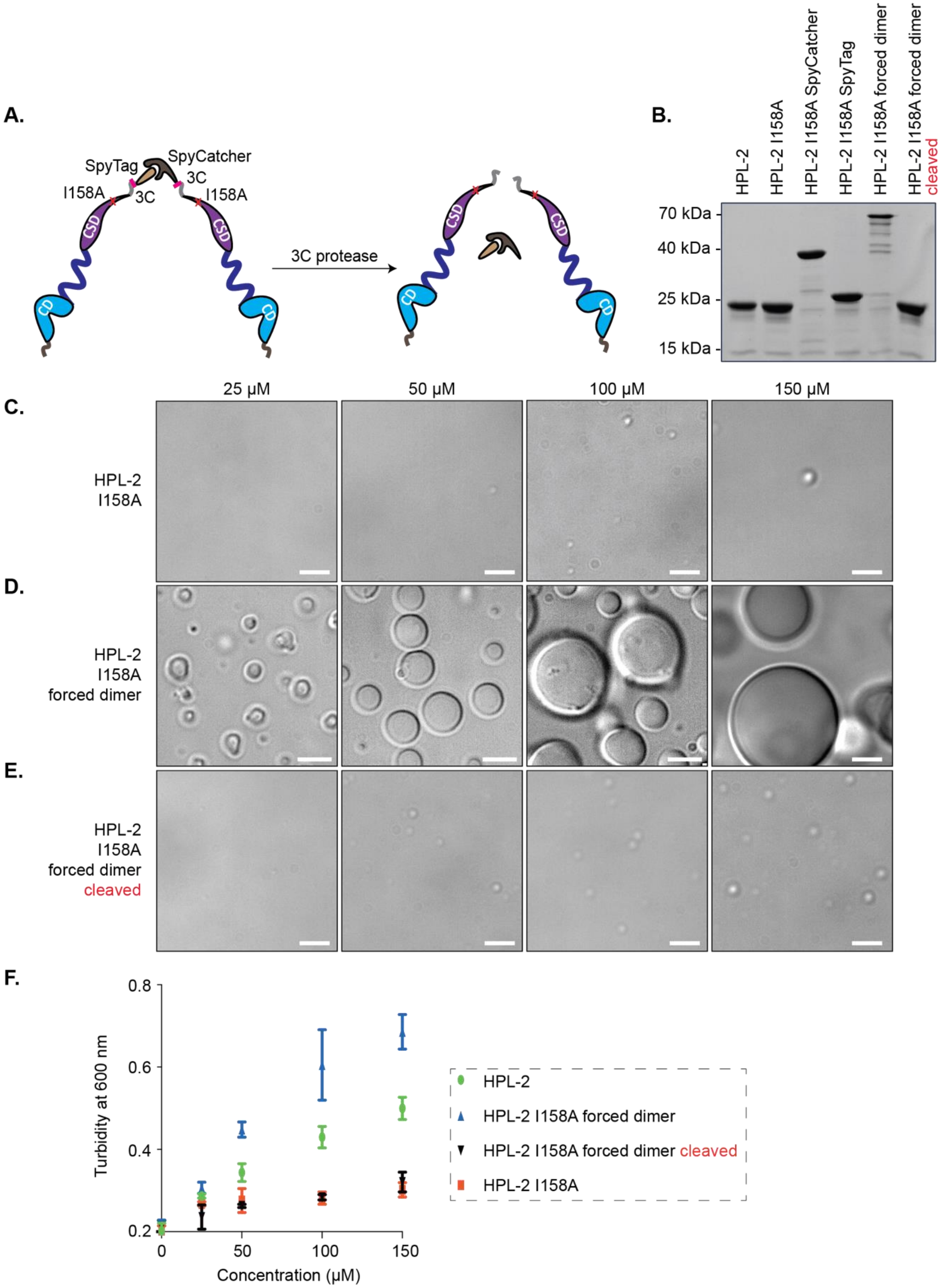

A.
SpyTag
SpyCatcher
I158A
3C
3C
I158A
CSD
CD
3C protease
B.
HPL-2
HPL-2 I158A
HPL-2 I158A SpyCatcher
HPL-2 I158A SpyTag
HPL-2 I158A forced dimer
HPL-2 I158A forced dimer cleaved
70 kDa
40 kDa
25 kDa
15 kDa
25 μM
50 μM
100 μM
150 μM
C.
HPL-2 I158A
D.
HPL-2 I158A forced dimer
E.
HPL-2 I158A forced dimer cleaved
F.
Turbidity at 600 nm
Concentration (μM)
0.2
0.4
0.6
0.8
0
50
100
150
HPL-2
HPL-2 I158A forced dimer
HPL-2 I158A forced dimer cleaved
HPL-2 I158A

**Figure S10. Forced dimerization restores phase separation of the HPL-2 I158A mutant protein**

**(A)** Scheme of the forced dimerization approach. HPL-2 I158A was C-terminally tagged with either SpyTag or SpyCatcher, preceded by a 3C protease recognition sequence. Equimolar amounts of tagged proteins were incubated for 1 hour at room temperature to induce covalent dimerization (HPL-2 I158A forced dimer). 3C protease cleaves the dimer and releases individual subunits (HPL-2 I158A forced dimer cleaved). **(B)** The indicated proteins were run on SDS-PAGE and stained with Coomassie blue. The running positions of molecular weight markers are indicated on the left. **(C–E)** Differential interference contrast (DIC) microscopy of HPL-2 mutant proteins and assemblies at increasing protein concentrations. Labelling as in (B). Images are representative of three independent experiments. Scale bars: 5 µm. (**F)** Plot of turbidity measured by absorbance reading at 600 nm against increasing concentrations of wild-type and mutant HPL-2 proteins. Condensate formation was induced as in (C–E). Data represent mean ± SD from three independent replicates.

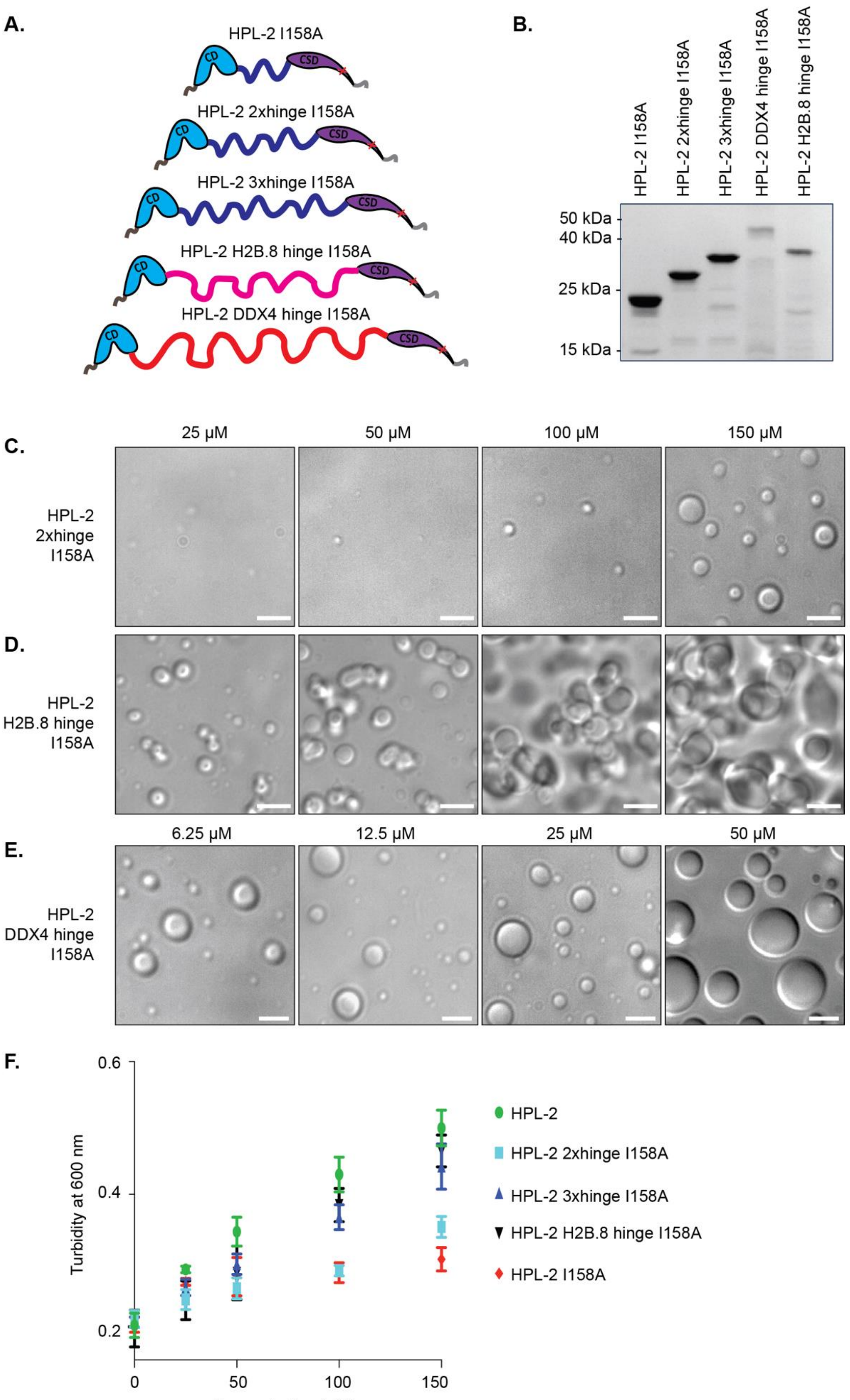
A.
HPL-2 I158A
HPL-2 2xhinge I158A
HPL-2 3xhinge I158A
HPL-2 H2B.8 hinge I158A
HPL-2 DDX4 hinge I158A
CD
CSD
B.
HPL-2 I158A
HPL-2 2xhinge I158A
HPL-2 3xhinge I158A
HPL-2 DDX4 hinge I158A
HPL-2 H2B.8 hinge I158A
50 kDa
40 kDa
25 kDa
15 kDa
C.
25 µM
50 µM
100 µM
150 µM
HPL-2 2xhinge I158A
D.
HPL-2 H2B.8 hinge I158A
E.
6.25 µM
12.5 µM
25 µM
50 µM
HPL-2 DDX4 hinge I158A
F.
Turbidity at 600 nm
0.6
0.4
0.2
0
50
100
150
Concentration (µM)
HPL-2
HPL-2 2xhinge I158A
HPL-2 3xhinge I158A
HPL-2 H2B.8 hinge I158A
HPL-2 I158A

**Figure S11. Analysis of mutants restoring phase separation of HPL-2 I158A**

**(A)** Scheme of different engineered HPL-2 proteins containing the I158A mutation. **(B)** The indicated proteins were run on SDS-PAGE and stained with Coomassie blue. The running positions of molecular weight markers are indicated on the left. **(C–E)** Differential interference contrast (DIC) microscopy of condensates formed by the indicated HPL-2 mutant proteins at increasing concentrations. Data are representative of three independent experiments. Scale bars: 5 μm. **(F)** Plot of turbidity measured by absorbance reading at 600 nm against increasing concentrations of HPL-2 wild-type and mutant proteins. Condensate formation was induced as in (C–E). Data represent mean ± SD from three independent replicates.

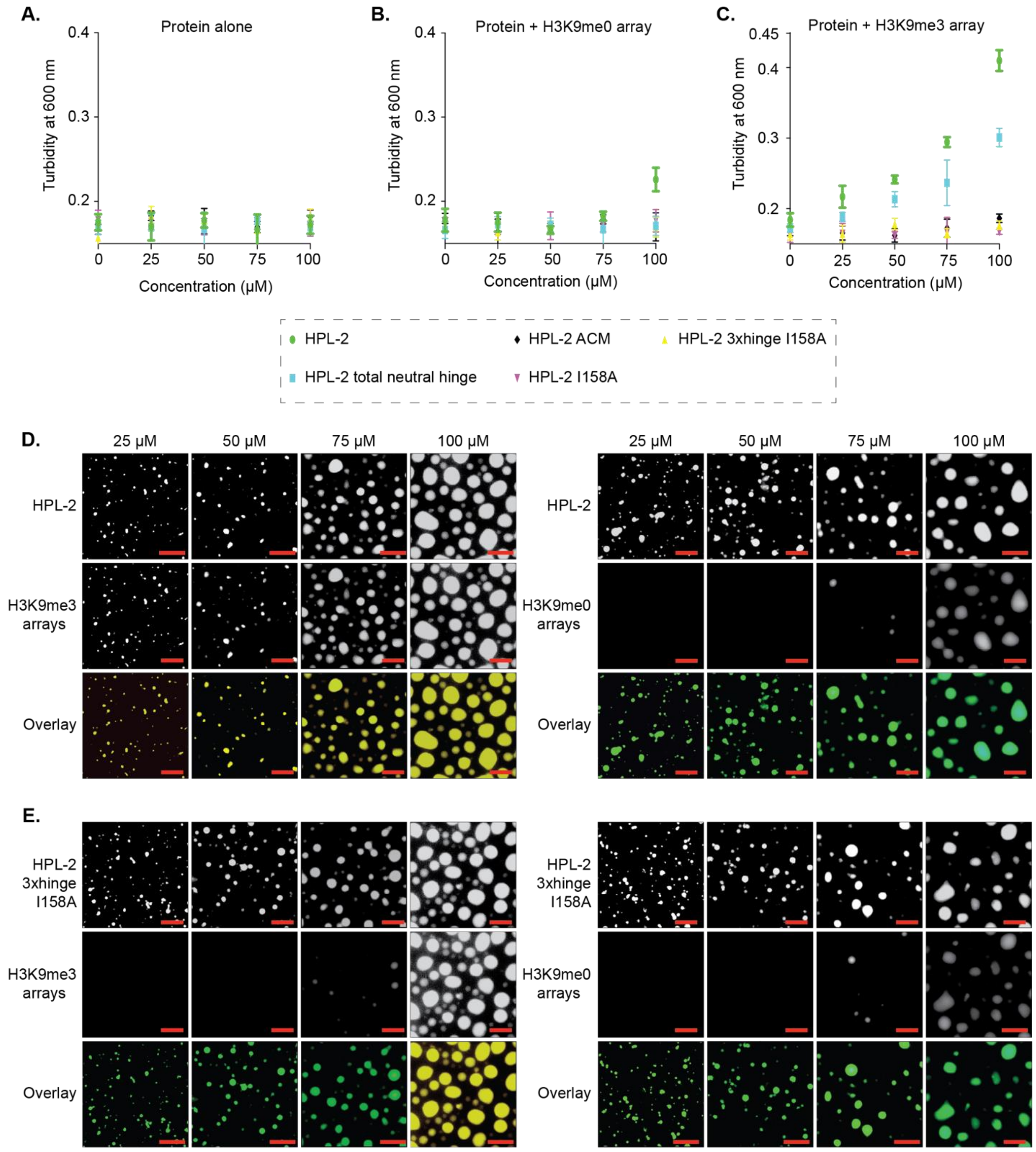


**Figure S12. Chromatin condensation and partitioning properties of HPL-2 wild-type and mutant proteins**

**(A–C)** Plot of turbidity measured by absorbance reading at 600 nm against increasing concentrations of HPL-2 wild-type and mutant proteins without chromatin arrays (A), in the presence of constant concentration of H3K9me0 chromatin arrays (B) or H3K9me3 chromatin arrays (C). Data represent mean ± SD from three independent replicates. **(D–E)** Representative fluorescence microscopy images of condensates of wild-type HPL-2 (D) or HPL-2 3xhinge I158A (E) formed in the presence of 10% (w/v) PEG8000 following addition of H3K9me0 or H3K9me3 chromatin arrays at the indicated protein concentrations. Data are representative of three independent experiments. Scale bars: 5 μm.

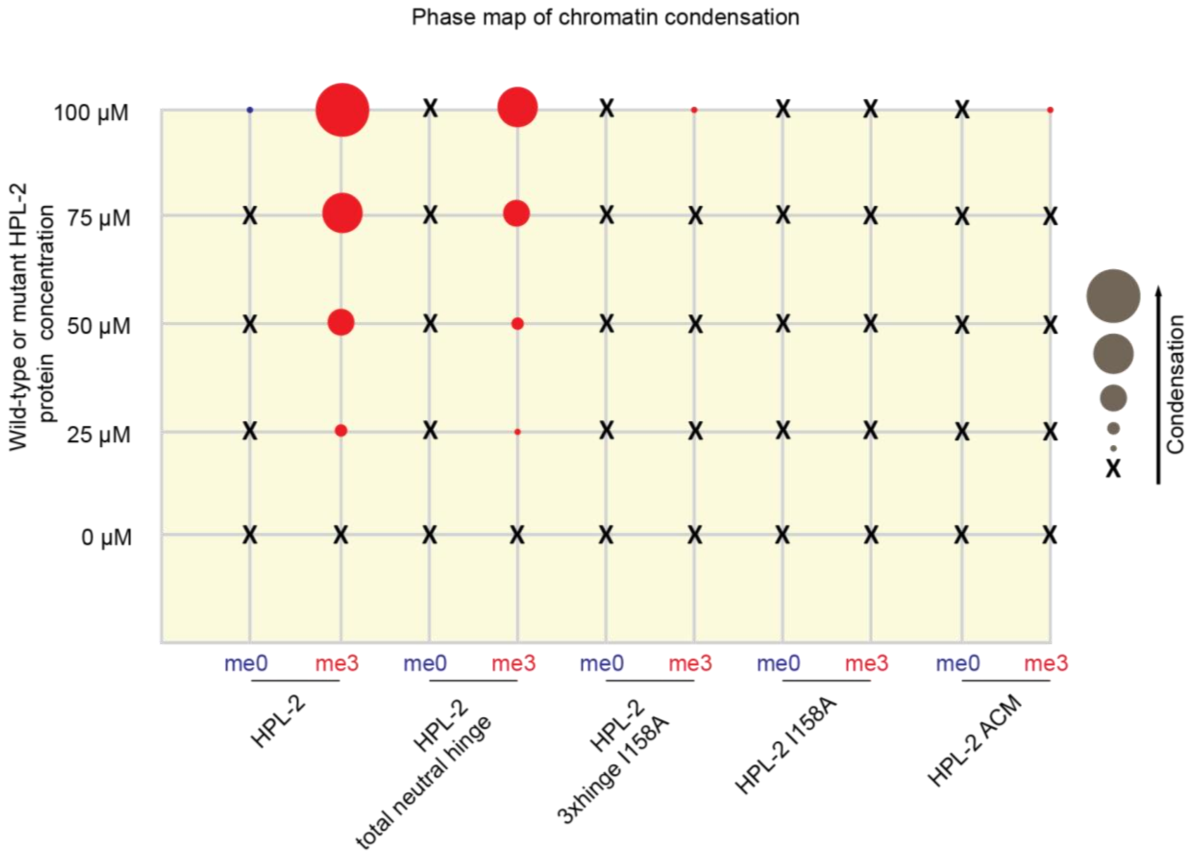


**Figure S13. Phase map summarizing HPL-2-mediated chromatin condensation**

Phase map of chromatin condensation by HPL-2 wild-type and mutant proteins at different concentrations and in presence of H3K9me0 or H3K9me3 chromatin arrays. Circle size indicates the degree of condensation (turbidity assay); 'x' denotes lack of condensation as determined by microscopy.

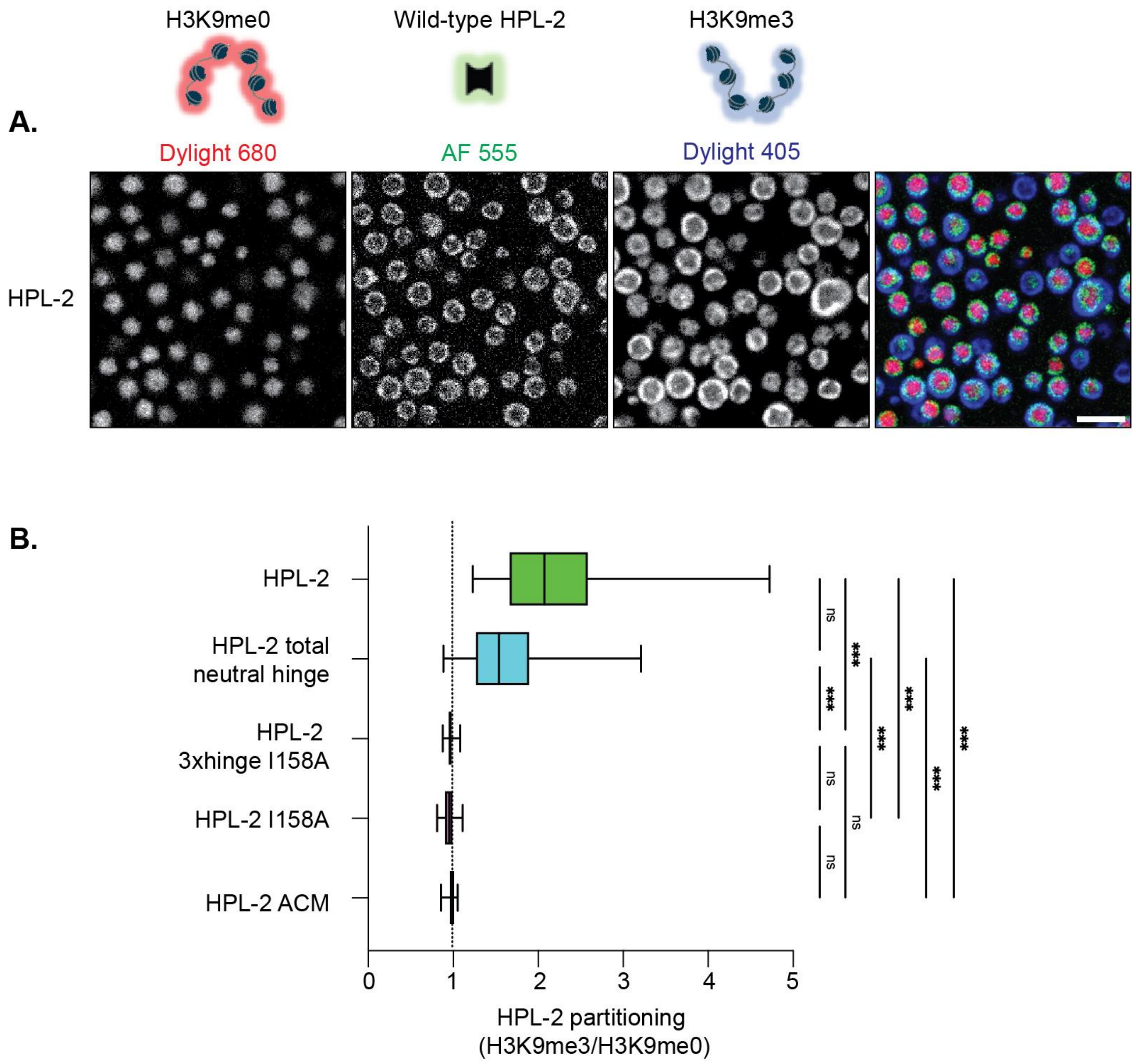


**Figure S14. Analysis of H3K9me3 chromatin segregation by HPL-2 wild-type and mutant proteins**

**(A)** Label-swap control: fluorescence microscopy of Dylight 680–labelled H3K9me0 and Dylight 405–labelled H3K9me3 chromatin arrays in the presence of wild-type HPL-2 protein trace-labelled with Alexa Fluor 555–HPL-2. Scale bars: 5 μm. Images are representative of three independent experiments. **(B)** Quantification of partitioning of HPL-2 wild-type or mutant proteins into the H3K9me3 chromatin region relative to the H3K9me0 chromatin region. Boxes extend from the 25th to the 75th percentile, with the median indicated by the vertical line. Whiskers represent the minimum and maximum values. N = 50 individual condensates from three independent experiments. *** P<0.001 by Kruskal-Wallis test with post hoc Dunn's test.

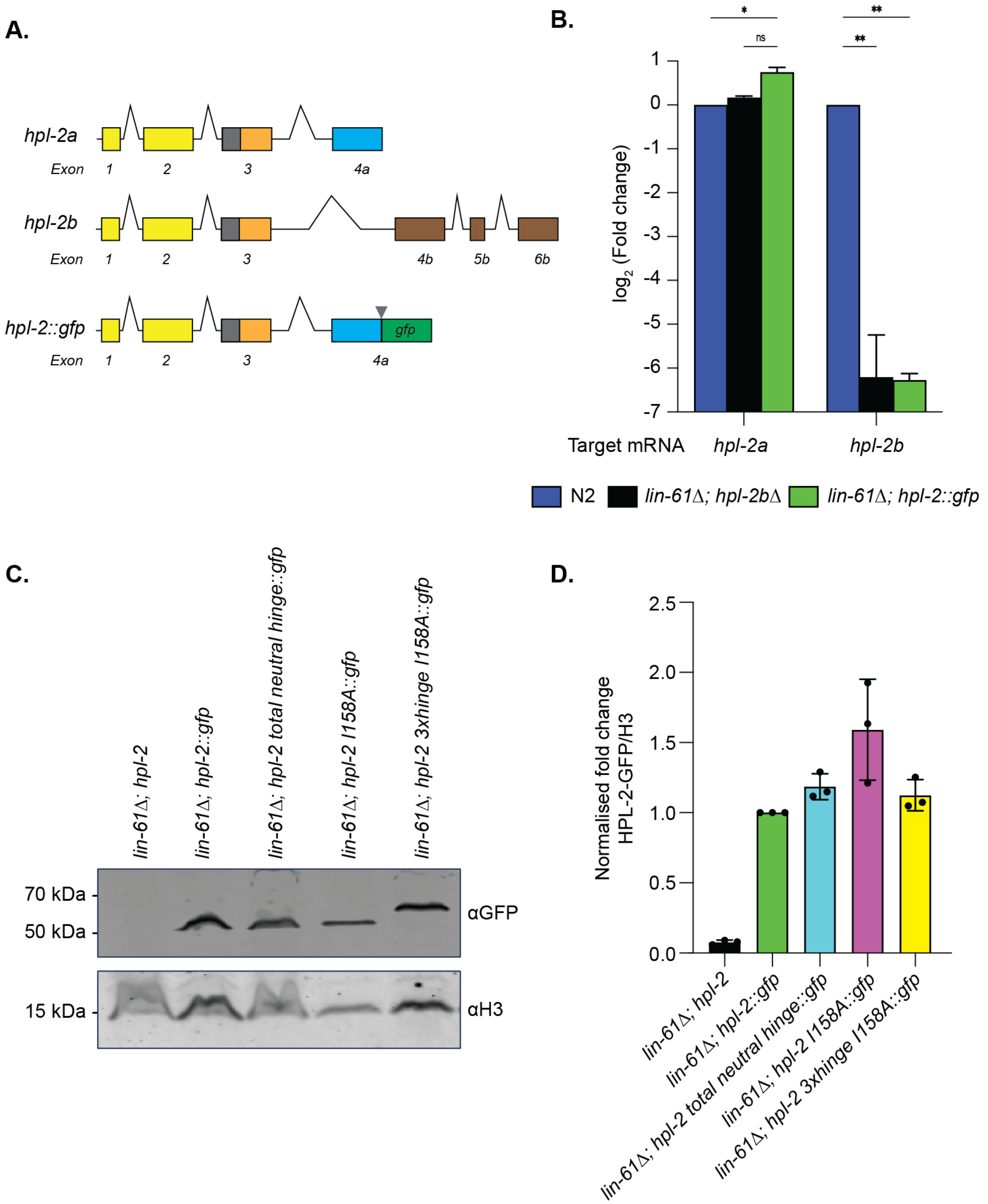


**Figure S15. Characterization of HPL-2::GFP wild-type and mutant *C. elegans* embryos**

**(A)** Scheme of the *hpl-2* gene locus that encodes for two splicing variants, *hpl-2a* (top), and *hpl-2b* (middle). Differential splicing of the ncRNA originating from the locus results in mRNAs that share the first three exons, corresponding to CD (yellow), hinge region (grey) and N-terminal portion of the CSD (orange) but differ in the sequences corresponding to the C-terminal portion and C-terminal extension of the CSD (exon 4 onwards; *hpl-2a,* orange; *hpl2b,* brown). A sequence encoding a GFP tag was introduced at the 5' end of exon 4A thereby eliminating the differential splicing event giving raise to HPL-2B. The strains used in this study therefore exclusively express wild-type HPL-2A::GFP or mutant fusion proteins. For simplicity, we annotate the corresponding *C. elegans* strains here as derivates of *hpl-2::gfp* and the resulting proteins as derivates of HPL-2::GFP. **(B)** The relative expression of the *hpl-2a* and *hpl2-b* mRNAs in *C. elegans* embryos of the indicated genetic background were measured by RT-qPCR and normalized to the level in the wild-type (N2) condition. Housekeeping gene *rpl-26* was analysed as an internal reference. N = 3 biological replicates; error bars indicate standard error of the mean. **(C)** Lysates of *C. elegans* embryos of the indicated genetic background were analysed by western blotting. The running position of molecular weight markers are indicated on the left. **(D)** Quantification of western blot analyses as in panel (C). Data represent the average fold enrichment of each HPL-2 mutant relative to H3, normalised to HPL-2::GFP expression in lane 2 in panel (C). N = 3 independent experiments; error bars indicate standard deviation.

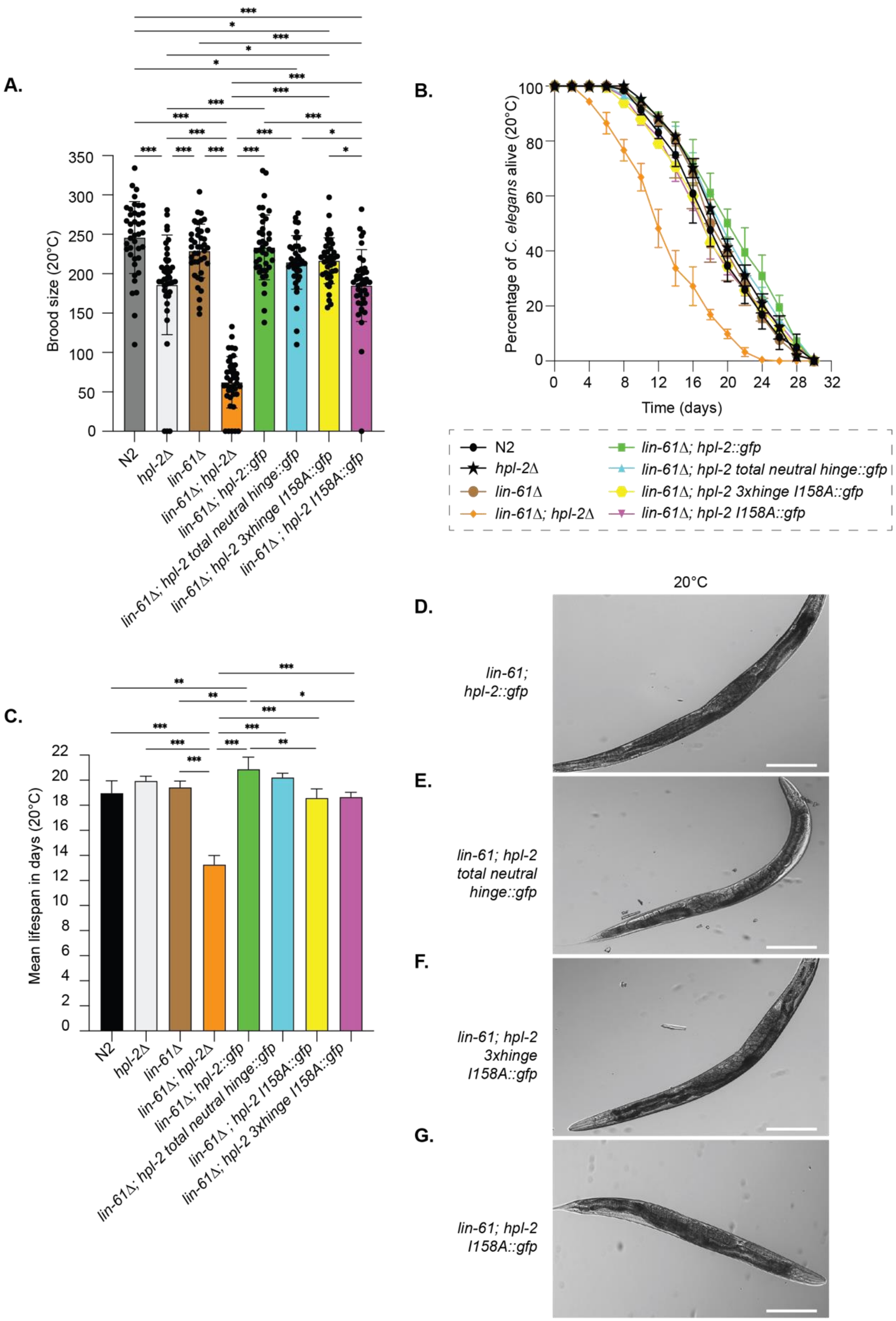
A.
Brood size (20°C)
350
300
250
200
150
100
50
0
N2
hpl-2Δ
lin-61Δ
lin-61Δ; hpl-2Δ
lin-61Δ; hpl-2::gfp
lin-61Δ; hpl-2 total neutral hinge::gfp
lin-61Δ; hpl-2 3xhinge I158A::gfp
lin-61Δ ; hpl-2 I158A::gfp
B.
Percentage of C. elegans alive (20°C)
Time (days)
N2
hpl-2Δ
lin-61Δ
lin-61Δ; hpl-2Δ
lin-61Δ; hpl-2::gfp
lin-61Δ; hpl-2 total neutral hinge::gfp
lin-61Δ; hpl-2 3xhinge I158A::gfp
lin-61Δ; hpl-2 I158A::gfp
C.
Mean lifespan in days (20°C)
N2
hpl-2Δ
lin-61Δ
lin-61Δ; hpl-2Δ
lin-61Δ; hpl-2::gfp
lin-61Δ; hpl-2 total neutral hinge::gfp
lin-61Δ ; hpl-2 I158A::gfp
lin-61Δ; hpl-2 3xhinge I158A::gfp
20°C
D.
lin-61; hpl-2::gfp
E.
lin-61; hpl-2 total neutral hinge::gfp
F.
lin-61; hpl-2 3xhinge I158A::gfp
G.
lin-61; hpl-2 I158A::gfp

**Figure S16. Effects of HPL-2 mutations on *C. elegans* fertility, lifespan, and vulval development at 20°C**

**(A)** Mean brood size of non-sterile *C. elegans* expressing HPL-2::GFP wild-type or mutant proteins in the background of *lin-61Δ* at 20°C. Data are representative of 40 worms analysed in four independent experiments. Error bars indicate standard deviation. *P<0.05, **P < 0.01, ***P < 0.001 by Kruskal–Wallis test with post hoc Dunn's test. **(B)** Lifespan curves of *C. elegans* expressing HPL-2::GFP wild-type or mutant proteins in the background of *lin-61Δ* at 20°C. The fraction of surviving worms (in percent) is plotted over time; Day 0 corresponds to the L4-to-adult molt. Data represent 225 worms from three independent experiments. Colour codes for genotypes are shown below the curves. **(C)** Mean lifespan (in days) of *C. elegans* expressing HPL-2::GFP wild-type or mutant proteins in the background of *lin-61Δ* at 20°C. Survival functions were estimated using the Kaplan–Meier method, and mean survival times were calculated as the area under the survival curve. Error bars denote standard error of the mean (SEM). *P < 0.05, **P < 0.01, ***P < 0.001 by Mantel–Cox log-rank test. **(D–G)** Representative differential interference contrast (DIC) microscopy images of *C. elegans* of the indicated genotypes at 20°C. Scale bars: 100 μm.

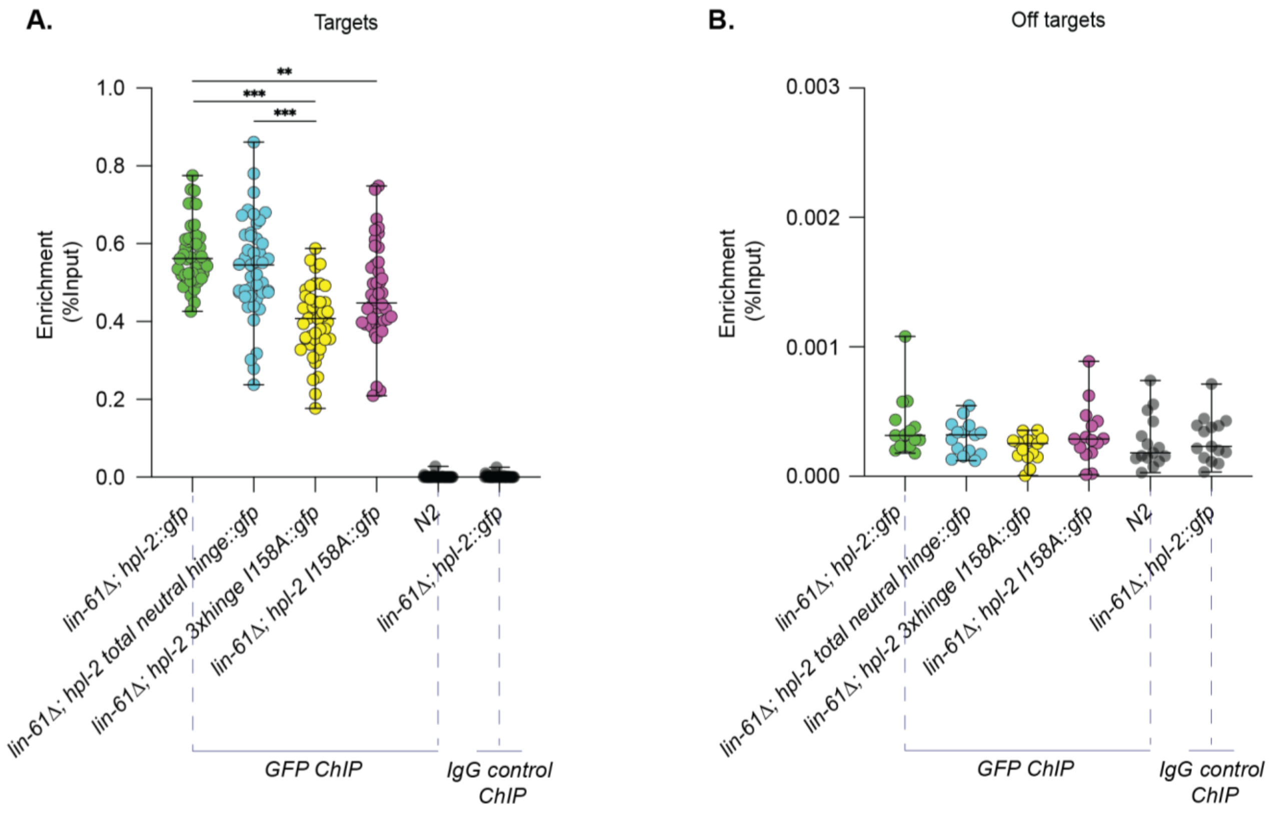


**Figure S17. HPL-2 mutant proteins deficient in dimerization retain substantial association with heterochromatin regions in *C. elegans* embryos**

**(A, B)** GFP ChIP-qPCR analysis of HPL-2 chromatin occupancy in *C. elegans* embryos expressing HPL-2::GFP wild-type or mutant proteins in the *lin-61Δ* background. Wild-type N2 embryos and IgG ChIP from *lin-61Δ; hpl-2::gfp* embryos were included as negative controls. Enrichment is shown as percentage of input recovered of 15 different target (A) and 5 different off-target (B) loci. Target loci were selected based on overlapping HPL-2 and H3K9me2 ChIP-seq enrichment in *C. elegans* embryos (taken from dataset GSE271919), whereas off-target loci were selected from chromosomal centre regions lacking enrichment for either factor. The analysis shown is absed on three independent biological replicates for each strain. $^{**}P < 0.01$, $^{***}P < 0.001$ by Kruskal–Wallis test with post hoc Dunn's test and Benjamini–Hochberg correction for multiple testing. Non-significant comparisons and comparisons with the negative controls (N2 and IgG ChIP) are not shown.

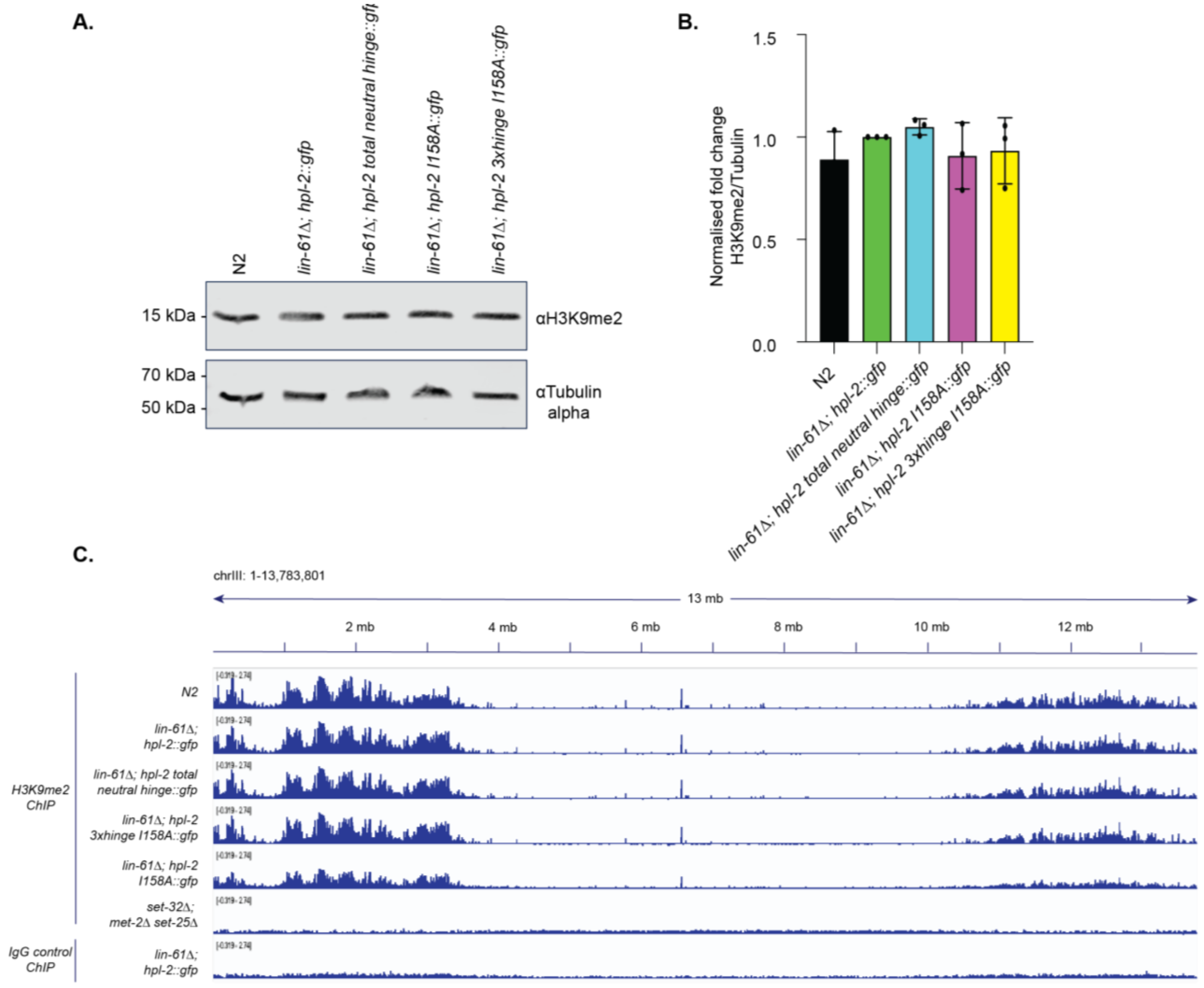


**Figure S18. Analysis of H3K9me2 levels in *C. elegans* embryos expressing HPL-2 mutant proteins**

**(A)** Lysates of *C. elegans* embryos of the indicated genetic background were analysed by western blotting. The running position of molecular weight markers are indicated on the left. N2, wild type. **(B)** Quantification of the western blot analyses as in panel (A). Data represent the average fold enrichment of H3K9me2 in each HPL-2 mutant relative to Tubulin-alpha, normalised to HPL-2 expression in lane 2 in panel (A). N = 3 independent experiments; error bars indicate standard deviation. **(C)** Representative Integrative Genomics Viewer (IGV) browser snapshot showing H3K9me2 ChIP-seq signal across chromosome III in the indicated *C. elegans* strains. Tracks are displayed using the same genomic coordinates and signal scale. The *set-32; met-2 set-25* strain [7] that is deficient in H3K9 KMTs and ChIP-seq performed with IgG serve as negative controls.

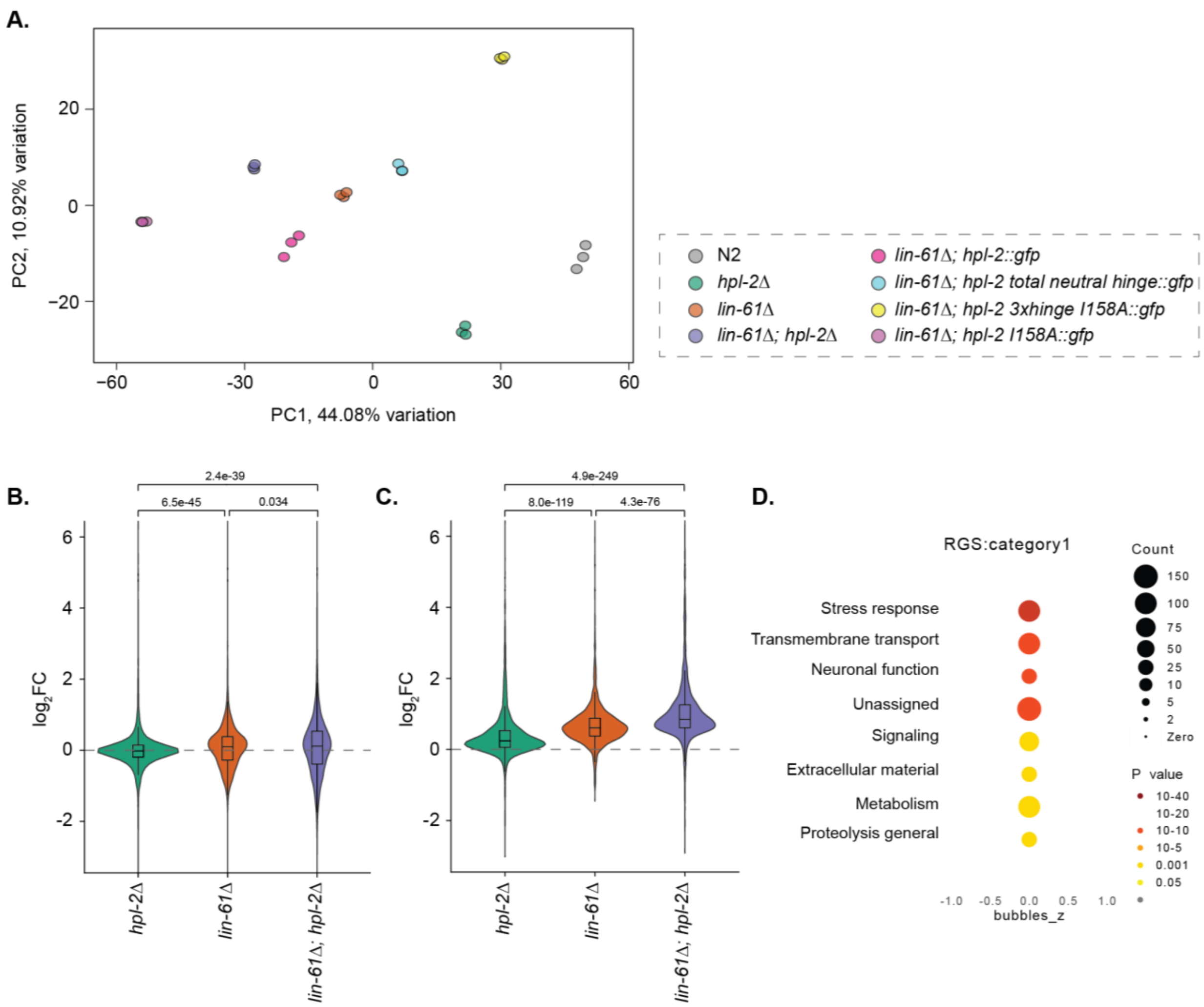


**Figure S19. Quality control and global differential gene expression analyses of RNA-seq datasets**

**(A)** Principal component analysis (PCA) of RNA-seq samples from all *C. elegans* strains analysed in this study. **(B)** Violin plot quantification of $log_2$ fold change ($log_2$FC) for all expressed genes (those not excluded by DESeq2 automatic filtering) located on the autosomal arms in embryos of the indicated mutant *C. elegans* strains, using N2 as the control. The violin plots show the distribution of $log_2$FC values, with the embedded boxplot indicating the interquartile range and median. Whiskers represent the data range. **(C)** Genes significantly upregulated in at least one sample (padj < 0.05; $log_2$FC > 0.5) relative to N2, quantified as violin plots with embedded boxplots showing the interquartile range and median. The FDR-adjusted p-values shown in (B, C) were obtained by a two-sided Wilcoxon rank sum test of all possible two-way comparisons of samples. Only significant (padj<0.05) comparisons are shown. (D) Bubble chart of enrichment categories for genes significantly upregulated on the autosomal arms in the embryos of the *lin-*

*61Δ; hpl-2 3xhinge I158A::gfp* relative to the control *lin-61Δ; hpl-2::gfp C. elegans* strain generated with WormCat 2.0. RGS: regulated gene set. Category 1 denotes the broad functional groups to which the enriched genes belong. Bubble size corresponds to the enrichment score.
